# Scalable Generation of Clinical-Grade Universal Human cDC1s Enables Potent Antitumor Immunotherapy

**DOI:** 10.64898/2026.01.14.699162

**Authors:** Can Liu, Siqi Bi, Wutao Chen, Ye Tian, Haowen Li, Guanhua Li, Honglin Li, You Wang, Li Wu, Haibo Zhou

**Affiliations:** Shanghai Institute of Rheumatology, Renji Hospital, School of Medicine, Shanghai Jiao Tong University (SJTUSM); Shanghai 200001, China; Department of Medical Oncology, Fudan University Shanghai Cancer Center, Department of Oncology, Shanghai Medical College, Fudan University, Shanghai 200032, China; Department of Obstetrics and Gynecology, Renji Hospital, School of Medicine, Shanghai Jiao Tong University (SJTUSM); Shanghai 200127, China; Hangzhou Institute of Medicine (HIM), Chinese Academy of Sciences, Hangzhou 310022, China; Department of Nephrology, The First Affiliated Hospital of Zhengzhou University; Zhengzhou 450052, China; Tsinghua-Peking Joint Center for Life Sciences, School of Basic Medical Science, Tsinghua Medicine, Tsinghua University, Beijing 100084, China

**Author notes:** These authors contributed equally (Can Liu, Siqi Bi, Wutao Chen, Ye Tian).

## Abstract

Despite decades of intensive clinical effort, monocyte-derived dendritic cell (DC) vaccines have not yet achieved sufficient clinical benefits in cancer therapy. Conventional type 1 DCs (cDC1s), a subset of antigen-presenting cells with superior cross-presentation capacity, are well established as pivotal mediators of antitumor immunity. However, their clinical translation has been hindered by the absence of scalable and efficient generation methods. Here, we develop a three-step strategy addressing this critical unmet need: a cost-effective, feeder cell-free, GMP-compatible approach enabling large-scale generation of cDC1s from human umbilical cord blood CD34^+^ hematopoietic stem and progenitor cells (CD34^+^ HSPCs). Starting from a single umbilical cord blood unit of CD34^+^ HSPCs, our method yields approximately 3.5×10^9^ pure cDC1s—sufficient for over 700 therapeutic doses. These cDC1s exhibit robust antigen cross-presentation activity and substantial production of antitumor cytokines. Critically, in various humanized tumor mouse models, they elicit significantly stronger antitumor efficacy than either moDCs or PD-1 monoclonal antibody (mAb) alone, pronouncedly remodel the tumor microenvironment (TME), and synergize with PD-1 mAbs to achieve enhanced therapeutic effects. Furthermore, our system recapitulates the complete *in vivo* DC differentiation trajectory: from CD34^+^ HSPCs through early-pre-DCs and pre-DCs to mature subsets (cDC1s, cDC2s, DC3s, mregDCs, ASDCs). This platform thus provides a powerful tool to dissect the regulatory mechanisms governing human DC subset specification. Overall, this work overcomes a long-standing bottleneck in cDC1-directed cancer immunotherapy and accelerates the clinical translation of DC subset-selective immunotherapeutic strategies.

## Introduction

Dendritic cells (DCs) are pivotal antigen-presenting cells that orchestrate both innate and adaptive immune responses in infections, chronic inflammatory diseases, and cancer(*1, 2*). They are further classified into distinct subsets, namely conventional DC subset (cDC) 1, cDC2, plasmacytoid DCs (pDCs), monocyte-derived DCs (moDCs), and the newly identified DC3(*1*). cDC1 and cDC2 are derived from common myeloid DC precursors(*3, 4*). In contrast, pDCs and DC3 primarily originate from the lymphoid pDC precursor (pre-pDC) and an independent Ly6c^+^ precursor within the macrophage-DC progenitor (MDP), respectively(*5, 6*). The development of cDCs and pDCs depends on fms related receptor tyrosine kinase 3 ligand (FLT3L), while that of DC3 relies on colony stimulating factor 2 (GM-CSF)(*1, 7*). The differentiation of cDC1 is regulated by a set of transcription factors (TFs), including interferon regulatory factor 8 (*IRF8*), ETS variant transcription factor 6 (*ETV6*), inhibitor of DNA binding 2 (*ID2*), basic leucine zipper ATF-like transcription factor 3 (*BATF3*) and zinc finger protein 366 (*ZNF366*)(*1*). In contrast, the differentiation of cDC2 is under the precise control of *IRF4* and zinc finger E-box binding homeobox 2 (*ZEB2*)(*1*). Additionally, the development of pDCs is modulated by transcription factor 4 (*TCF4*), whereas that of DC3 is governed by *IRF8* and KLF transcription factor 4 (*KLF4*)(*1, 6*).

cDC1s, which specialize in cross-presenting tumor antigens to cytotoxic CD8^+^ T cells, are critical for antitumor immunity(*8*). Their transcriptional signature and frequencies positively correlate with improved survival and treatment responsiveness across multiple human cancers(*9, 10*). cDC1s transport intact antigens to lymph nodes(*8*), and elicit long-lasting immune responses by activating CD4^+^ T cells(*11*). They outperform cDC2s in cross-presenting cell-associated antigens, and unlike moDCs, exogenous cDC1-based therapies do not rely on endogenous cDC1s(*12*). cDC1s are characterized by a high-level expression of C-Type lectin domain family 9 member A (CLEC9A), which binds to actin filaments exposed by necroptotic cells and subsequently induces phagosomal rupture, releasing dead-cell-associated antigens into the cytosol. These antigens are then processed via the major histocompatibility complex (MHC)-I pathway to prime antigen-specific CD8^+^ T cells(*8*). The crosstalk between cDC1 and other immune cells within the tumor microenvironment (TME) is critical for anti-tumor responses. cDC1-derived interleukin (IL)-12p70 can induce the cytotoxic activity of natural killer (NK) cells and CD8^+^ T cells, as well as stimulate these cells to produce IFN-γ(*13*). Additionally, cDC1-secreted chemokines C-X-C motif chemokine ligand (CXCL) 9 and CXCL10 recruit C-X-C motif chemokine receptor (CXCR) 3^+^ innate lymphocytes and T cells into tumors(*13*). Conversely, IFN-γ, produced by NK cells and T cells, can be sensed by intratumor cDC1s, promoting further IL-12p70 production(*13*). X-C motif chemokine ligand (XCL) 1 and C-C motif chemokine ligand (CCL) 5, secreted by tumor-infiltrating CD8^+^ T cells and NK cells, can recruit more cDC1 to the tumor sites, establishing a positive feedback loop(*13*). Notably, the C-C motif chemokine receptor (CCR) 7 negative cDC1 subset is essential for the terminal maturation of antigen-specific CD8^+^ T cells localized in the tertiary lymphoid structures within tumors(*14*).

Diverse studies have shown that, compared with moDCs, which are easily accessible and commonly used in clinical DC therapies, cDC1s possess significantly greater potential for tumor suppression in mouse model(*15*). Thus, cDC1s are regarded as the preferred subset for DC-based therapeutic strategies. However, existing protocols, aiming to enhance the induction efficiency of human cDC1, have not yet achieved the scale required for clinical application(*13*). Therefore, the development of a simple and feasible cDC1 induction system is critical for advancing cDC1-based cell therapies.

In this study, we have successfully established a serum-free production system for human cDC1 and cDC2. This system faithfully recapitulates the *in vivo* differentiation trajectory of cDCs and enables the generation of a substantial quantity of cDC1 cells from a single unit of umbilical cord blood (CB) - derived CD34^+^ stem cells, with this quantity being sufficient to support over 700 patient treatments. Significantly, these cDC1s generated *in vitro* have been validated to exhibit broad-spectrum anti-tumor efficacy across multiple humanized solid tumor models.

## Results

### *Ex Vivo* Generation of Human Conventional Dendritic Cells

Human CB derived CD34^+^ HSPCs obtained from healthy donors were cryogenically preserved and utilized to produce CB derived cDCs (CBcDCs) in a three-stage, 14-day process (Fig. 1A). During the first stage, the freeze-thawed CD34^+^ HSPCs underwent expansion in the (Good Manufacturing Practices) GMP-grade StemSpan™-AOF medium supplemented with FLT3L, stem cell factor (SCF), thrombopoietin (TPO) and StemRegenin (SR) 1 for 4 consecutive days. In the second stage, HSPCs were induced to differentiate into CBcDCs through a 10-day cultivation with different combinations of additives (Fig. 1A). Finally, cDC1s were purified and subsequently matured using Toll-like receptor (TLR) agonists.

**Fig. 1.**
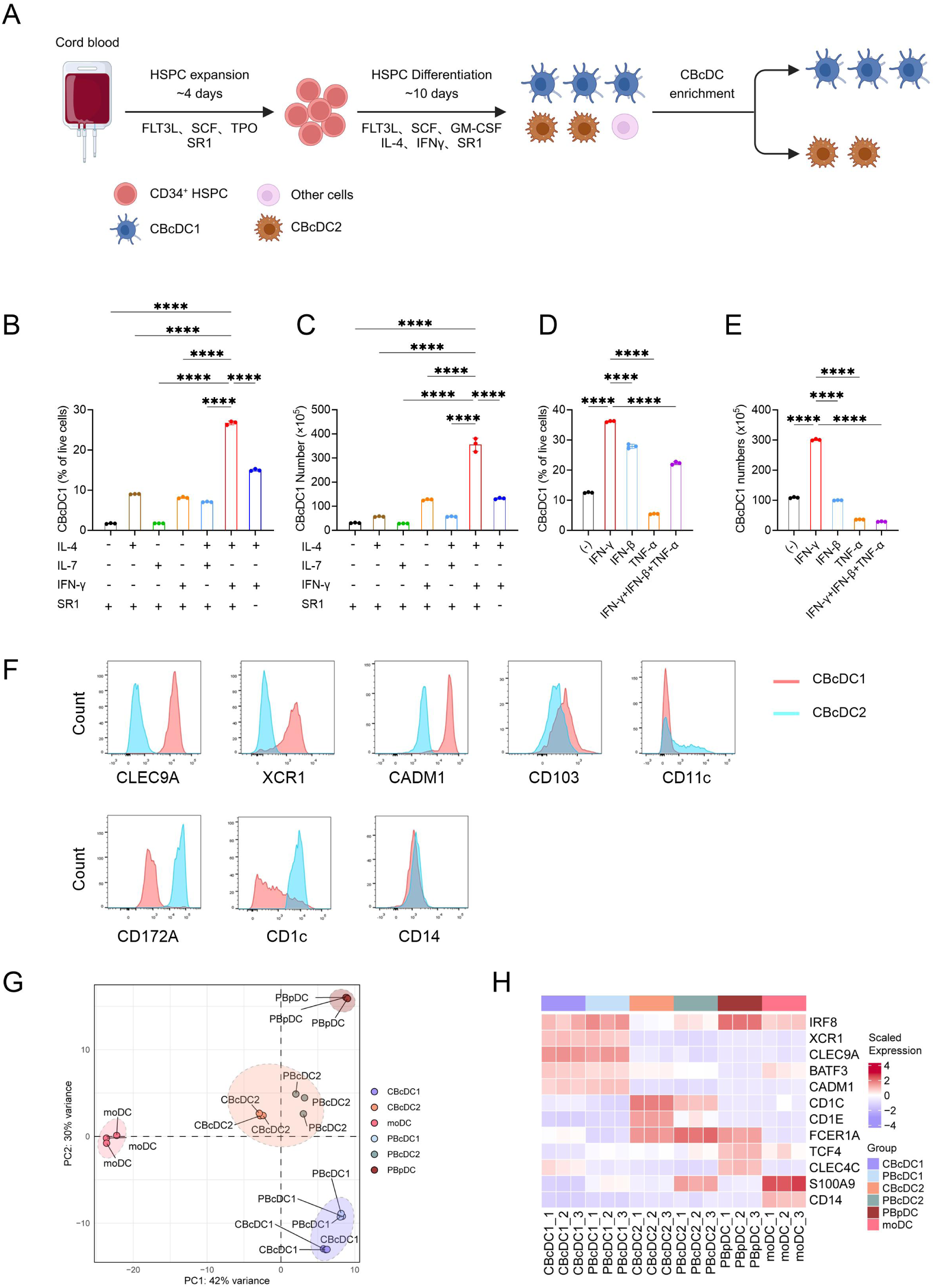
Allogeneic cDCs are Manufactured at High yield, Convenience and Robustness. (**A**) Schematic representation illustrating the generation of allogeneic CBcDCs from CB HSPCs utilizing GMP-compliant materials in a feeder cell-free system. (**B**, **C**) Impact of cytokine/small molecule compound combinations (IL-4, IL-7, IFN-γ, and SR1) on CBcDC induction. Induction efficiency (**B**) and absolute yield (**C**) of CBcDC1 cells generated from 1×10^5^ CB HSPC. (**D**, **E**) Identification of optimal cytokines for CBcDC1 induction. The induction efficiency (**D**) and the yield (**E**) of CBcDC1 produced from 1×10^5^ CB HSPC. N=3. (**B**-**E**) data are presented as the mean ± SD and were analyzed by one-way ANOVA with Dunnett’s multiple comparisons test. ****p<0.0001. (**F**) Expression of marker genes in CBcDC1 and CBcDC2. (**G**) Principal component analysis (PCA) of different DC subpopulations. PBcDC1, peripheral blood cDC1; PBcDC2, peripheral blood cDC2; PBpDC, peripheral blood pDC; moDC, monocyte derived DC. Triplicate samples were analyzed for each cell type. (**H**) Characteristic gene expression profiles of different DC subpopulations.

The cytokine combination of SCF, FLT3L and GM-CSF has been conventionally adopted for the *in vitro* induction of human cDCs(*16*). Consequently, we then set out to investigate how to further boost the induction efficiency of cDCs under this culture setting. The CD34^+^ CB stem cell derived cDC1_like cell (CBcDC1) was defined as Lineage^-^AXL^-^HLA-DR^+^CD123^-^CD141^high^CLEC9A^+^, while the CD34^+^ CB stem cell derived cDC2_like cell (CBcDC2) was defined as Lineage^-^AXL^-^HLA-DR^+^CD123^-^CD141^low/-^CD1c^+^ (fig. S1A). The supplementation of cytokines generally resulted in a reduction in the overall expansion fold of total cell numbers (fig. S1B). More precisely, the addition of IL-7 had minimal effect, whereas the addition of IL-4 or IFN-γ could augment the induction efficiency of CBcDC1 (Fig. 1B and C). To be specific, the proportion of CBcDC1 within total live cells increased from a mean value of 1.71% to 9.03% upon the addition of IL-4 and to 8.14% following the addition of IFN-γ (Fig. 1B). Co-application of IL-4 and IFN-γ substantially elevated the induction efficiency of CBcDC1 to 26.73% (Fig. 1B). Furthermore, SR1, an aryl hydrocarbon receptor antagonist that promotes the expansion of human HSCs(*17*), is also an essential component. Its absence reduced the induction efficiency of CBcDC1 to 15.0% (Fig. 1B). Correspondingly, the combined application of IFN-γ, IL-4 and SR1 resulted in an average 355.36-fold expansion in the number of CBcDC1, which was far higher than that achieved under other conditions (Fig. 1C). In contrast, the addition of IFN-γ and SR1, or the combined use of IL-4, IFN-γ and SR1 only slightly enhance the differentiation of CBcDC2 (fig. S1C and D). Previous reports have indicated that, aside from IFN-γ, type I IFN or tumor necrosis factor (TNF)-α can also facilitate the induction of cDC1(*18*). Therefore, we compared these three cytokines. The results indicated that the utilization of IFN-γ yielded the highest proportion and number of cDC1 (Fig. 1D and E). When further assessing the influence of basal medium on the differentiation of CBcDC1, we found that the StemSpan™-AOF medium held a substantial advantage over the X-VIVO 15 medium, which is prevalently utilized in clinical cell therapies (fig. S1E). Collectively, we concluded that the optimal condition for CBcDC1 differentiation involved supplementing StemSpan™-AOF medium with a combination of SCF, FLT3L, GM-CSF, IL-4, IFN-γ, and SR1 (G4γS condition).

It should be emphasized that the IFN-γ is a potent inducer of *IRF8*, a crucial TF promoting the differentiation and maintenance of cDC1(*19–22*). Our dynamic study confirmed that the IFN-γ was capable of inducing the expression of characteristic cDC1 genes, such as TFs (*IRF8*, *BATF3*, *ZNF366*, *ID2*), surface markers (*XCR1*, *CADM1*, *CLEC9A*) and the functional molecule (*WDFY4*), as well as the cDC2 gene *KLF4* and the DC progenitor gene *FLT3* (fig. S1F). Conversely, the addition of IFN-γ was unable to further induce other cDC1 gene (*NFIL3*), cDC2 genes (*IRF4*, *ZEB2*, *NOTCH2*, *SIRPA*, *MRC1*, *ITGAX*, and *CLEC10A*), DC progenitor genes (*CD34*, *CD33*, and *SPI1*) and the mregDC gene (*LAMP3*) (fig. S1F). Simultaneously, during the process of differentiation, as the expression of cDC genes was upregulated, the expression of the stem cell gene *CD34* and the pDC gene *TCF4* gradually declined (fig. S1F). Gene Ontology (GO) analysis revealed that the genes induced by IFN-γ were significantly enriched in the pathways associated with cDC1 functions, such as antigen processing and presentation pathways (fig. S1G).

Subsequently, we proceeded to detect the expression of characteristic surface markers in CBcDC subsets (Fig. 1F). The cDC1-signature makers, including CLEC9A, XCR1, and CADM1, were highly expressed in CBcDC1. In contrast, the cDC2-signature markers, such as CD172A and CD1c, were highly expressed in CBcDC2. Different from CBcDC2, CBcDC1 did not express the CD11c molecule (Fig. 1F). Additionally, the moDC-signature markers, for example, CD14, were not expressed in either CBcDC1 or CBcDC2.

To further verify the characteristics of the induced cells, we performed principal component analysis (PCA) on both the CBcDC subsets and those DC subsets sorted from adult peripheral blood. Results manifested that CBcDC1 was analogous to the endogenous cDC1 subset (PBcDC1) and CBcDC2 was analogous to the endogenous cDC2 subset (PBcDC2), while their relationship with pDC and moDC was much more distant (Fig. 1G). In line with this, the expression patterns of DC signatures demonstrated a closer similarity between CBcDC1 and PBcDC1, as well as between CBcDC2 and PBcDC2 (Fig. 1H).

Thereafter, we proceeded to investigate the impacts of TFs on the differentiation of CBcDCs. CRISPR/Cas9-mediated deletion of *IRF8* and *BATF3* attenuated the potential of HSPCs to differentiate into CBcDC1 (fig. S2A). The ablation of *BATF3* concurrently enhanced the predisposition for differentiation towards CBcDC2 (fig. S2B). While the perturbation of *IRF4* expression dampened the propensity for CBcDC2 production, the targeted deletion of *ZEB2* actively fostered the generation of CBcDC1 at the expense of CBcDC2 (fig. S2C and D). Moreover, the disruption of the *IFNGR1* signaling pathway exerted a highly specific inhibitory effect on the differentiation of CBcDC1 (fig. S2A and B). Conversely, the knockout of *TCF4*, a crucial TF that orchestrates pDC differentiation, had no effect on the differentiation of either CBcDC1 or CBcDC2 (fig. S2A and B). Ectopic expression of *BATF3* and *ZNF366*, two TFs critical for cDC1 terminal differentiation, further enhanced cDC1 induction efficiency to 87.8% and 79.7%, respectively (fig. S2E).

Ultimately, we probed into the efficacy of this refined approach across samples sourced from diverse donors. The total number of cells derived from the initial CD34^+^ cord blood HSPCs exhibited an expansion that ranged from a minimum of 769-fold to a maximum of 3,842-fold, with an average expansion of 2,043-fold (fig. S2F). For CBcDC1s, the induction efficiency ranged from 20.8% to 50.0% (mean: 33.46%; fig. S2G), with concomitant expansion of 200-fold to 1,614-fold (mean: 705-fold; fig. S2H). By contrast, application of this protocol to peripheral blood-derived HSPCs resulted in a marked reduction in cDC1 induction efficiency (fig. S2I).

### Identify the Subpopulations of CBcDC1 and CBcDC2

To delineate the functional properties of CBcDC1s, we systematically assessed their cytokine-secretion profiles in response to distinct TLR ligands. Stimulation of CBcDC1s with the TLR3 agonist Poly(I:C) alone elicited merely modest cytokine production. Strikingly, co-stimulation with either the TLR1/2 agonist Pam3CSK4 or the TLR7/8 agonist R848 induced robust secretion of IL-12p70, IFN-λ, TNF-α, IL-6 and CXCL10 (IP-10) (Fig. 2A). By contrast, co-stimulation with the TLR9 ligand CpG did not augment the production of these aforementioned cytokines (Fig. 2A). Subsequently, we designed experiments to evaluate the antigen presentation capacity of CBcDC1 (fig. S3A). The HLA-A*02 negative SKOV3 cell line was transfected with the human CTAG1B gene (encoding the NY-ESO-1 antigen) using a lentiviral vector to construct the HLA-A*02 negative SKOV3-CTAG1B cell line. The overexpression of CTAG1B was confirmed (fig. S3B). CBcDC1 derived from HLA-A*02:01 CB was utilized, and primary HLA-A*02 negative CD8^+^ T cells were engineered to overexpress the T-cell receptor (TCR) that specifically recognized the HLA-A*02:01 restricted NY-ESO-1_157-165_ MHC-peptide complex (fig. S3A). The expression of TCR was successfully detected on the surface of T cells with positive infection indicated by GFP fluorescence (fig. S3C). When incubated with the NY-ESO-1 peptides, CBcDC1 efficiently induced the proliferation of TCR-T cells without the addition of stimulants (Fig. 2B). Likewise, CBcDC1 displayed remarkable cross-presentation capacity, leading to the proliferation of TCR-T cells upon incubation with the CTAG1B protein (Fig. 2B). In contrast, the cross-presentation of antigens from live- or irradiated-SKOV3 cells by CBcDC1 necessitates extra innate immune activation (Fig. 2B). Of greater significance, the antigen presentation potential of CBcDC1s toward TCR-T cells is gradient-dependent. When CBcDC1s were co-incubated with peptides (Fig. 2C) or tumor cells (Fig. 2D), a stepwise increase in the CBcDC1: TCR-T cell ratio was accompanied by a progressive enhancement of TCR-T cell proliferation. Correspondingly, IFN-γ was also strongly induced during this process (fig. S3D and E). Meanwhile, the phagocytosis of cell-related antigens by CBcDC1 also gradually increased with the incubation time (fig. S3F). Overall, when co-cultured with live tumor cells, CBcDC1 relied on innate immune stimulation to efficiently cross-activate antigen-specific CD8^+^ T cells.

**Fig. 2.**
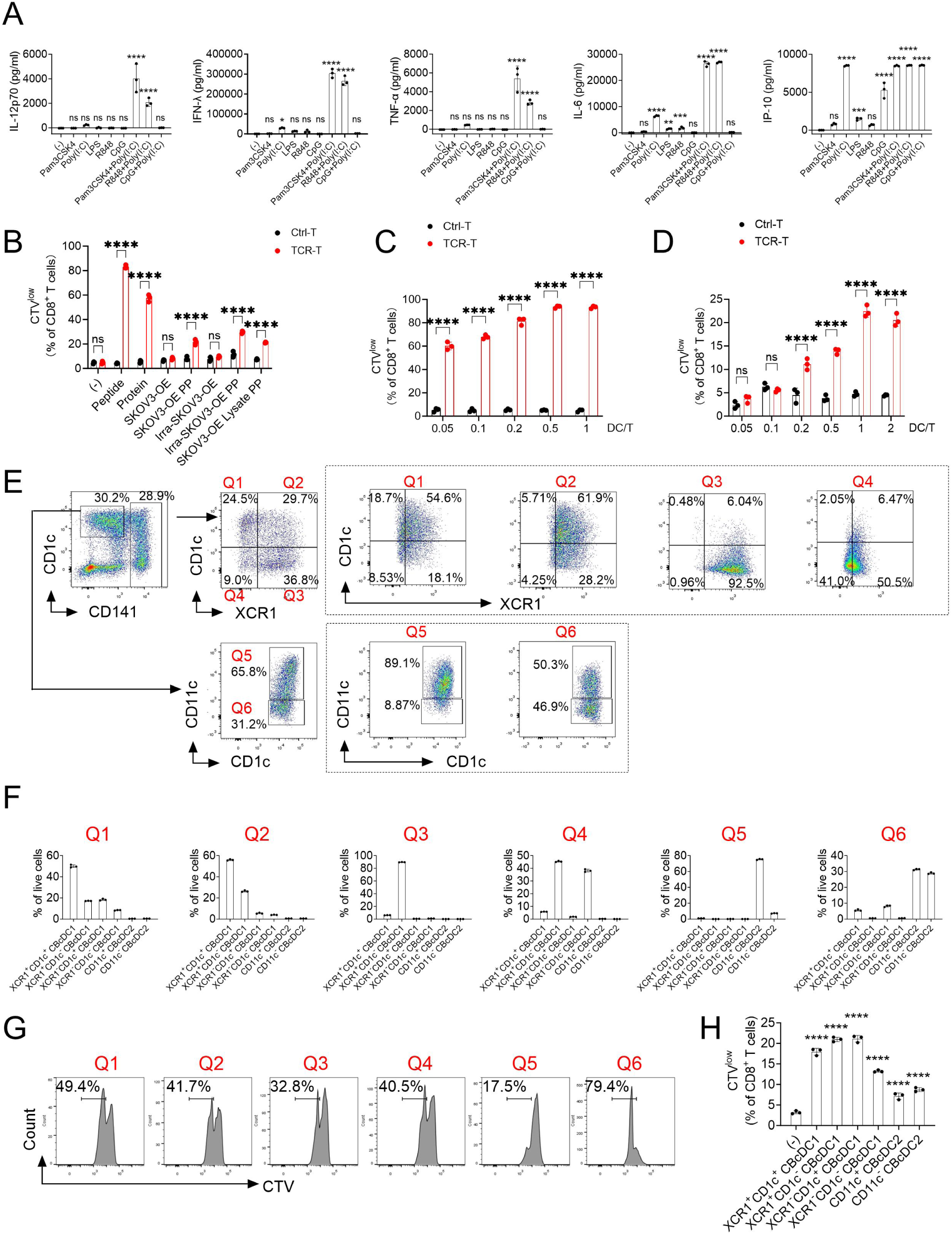
Identify the Subclusters of CBcDC1 and CBcDC2. (**A**) CBcDC1 were stimulated with the indicated stimulus, and the expression of IL-12p70, IFN-λ, TNF-α, IL-6 and IP-10 were evaluated. (**B**) The cross-presentation ability of CBcDC1. 5×10^4^ HLA-A*02:01 human CBcDC1 were co-cultured with 1×10^5^ HLA-A*02-negative human primary CD8^+^ T cells transduced with a TCR specific for the HLA-A*02:01-restricted NY-ESO-1 peptide (NY-ESO-1_157-165_ targeted TCR-T). Antigens from different sources were tested: Peptide, a pool of lyophilized 15-mer peptides covering the entire human NY-ESO-1 protein; Protein, CTAG1B protein; SKOV3-OE, HLA-A*02-negative SKOV3 cell line with CTAG1B overexpression; SKOV3-OE PP, HLA-A*02-negative SKOV3 cell line with CTAG1B overexpression and stimulated with Pam3CSK4 and Poly(I:C); Irra-SKOV3-OE, X-ray irradiated HLA-A*02-negative SKOV3 cell line with CTAG1B overexpression; Irra-SKOV3-OE PP, X-ray irradiated HLA-A*02-negative SKOV3 cell line with CTAG1B overexpression and stimulated with Pam3CSK4 and Poly(I:C); SKOV3-OE Lysate PP, whole cell lysate from HLA-A*02-negative SKOV3 cell line with CTAG1B overexpression and stimulated with Pam3CSK4 and Poly(I:C); CTV, CellTrace Violet. (**C**) The activation of TCR-T by CBcDC1 at varying DC:T cell ratios following peptide incubation. (**D**) The cross-activation of TCR-T by CBcDC1 at varying DC:T cell ratios following tumor cell (SKOV3-OE) incubation. (**E**) Representative flow cytometry plots showing the gating strategy for CBcDC subclusters and their differentiation potential. At day 8 of DC-oriented differentiation, four CBcDC1 subclusters were defined by XCR1 and CD1c expression, and two CBcDC2 subclusters by CD11c expression. These six CBcDC subclusters were sorted and cultured in differentiation medium for an additional 2 days to assess their differentiation capacity. Plots outside the dashed box show the initial differentiation states, and those inside show further differentiation outcomes after continued culture. Q1, XCR1^-^CD1c^+^ CBcDC1; Q2, XCR1^+^CD1c^+^ CBcDC1; Q3, XCR1^+^CD1c^-^ CBcDC1; Q4, XCR1^-^CD1c^-^ CBcDC1; Q5, CD11c^+^ CBcDC2; Q6, CD11c^-^ CBcDC2. (**F**) Quantification of the differentiation potential of CBcDC subclusters from panel E. (**G**) Proliferation potential of CBcDC subclusters during a 2-day redifferentiation process, evaluated by CTV labeling. (**H**) The cross-presentation ability of CBcDC subclusters. Both CBcDC1 and CBcDC2 subclusters were co-cultured with the CTAG1B-overexpressing HLA-A*02-negative SKOV3 cell in the presence of Pam3CSK4 and Poly(I:C) stimulation. Subsequently, CBcDCs were washed and incubated with the CTV-labeled NY-ESO-1_157-165_ targeted TCR-T for 48 hours. N=3. Data are presented as the mean ± SD: (**A)** and (**H**) were analyzed by one-way ANOVA with Dunnett’s multiple comparisons test; (**B**-**D**) by two-way ANOVA with Bonferroni post-hoc test. *p<0.05, **p<0.01, ***p<0.001, ****p<0.0001, ns, no significant.

When further delving into the CBcDC subsets, we found that CBcDC1 could be classified into 4 subpopulations according to the expression of XCR1 and CD1c. Moreover, CBcDC2 was also divided into CD11c^+^ and CD11c^-^ subpopulations (Fig. 2E). To elucidate the interrelationships among these subgroups, we individually sorted each subpopulation and subsequently induced their *in vitro* redifferentiation for 2 and 4 days (Fig. 2E and F, and fig. S4A). The XCR1^-^CD1c^+^ CBcDC1 were found to be able to give rise to XCR1^+^CD1c^+^, XCR1^-^CD1c^-^, and XCR1^+^CD1c^-^ CBcDC1s. Conversely, both XCR1^+^CD1c^+^ and XCR1^-^CD1c^-^ CBcDC1s were restricted to differentiating into XCR1^+^CD1c^-^ CBcDC1. Remarkably, the XCR1^+^CD1c^-^ CBcDC1 represented the terminal stage of CBcDC1 development and was devoid of the capacity to redifferentiate into other CBcDC1 subpopulations (Fig. 2E and F, and fig. S4A). For CBcDC2s, the CD11c^-^ subgroup was capable of differentiating into CD11c^+^ cells, along with a small proportion of CBcDC1s. In line with the redifferentiation ability, the proliferation potential varies across different subpopulations. Among CBcDC1 subgroups, XCR1^-^CD1c^+^ cells demonstrated the most robust proliferative capacity. This was succeeded by XCR1^+^CD1c^+^ and XCR1^-^CD1c^-^ CBcDC1s, while XCR1^+^CD1c^-^ cells exhibited the lowest proliferative potential. Within the CBcDC2 population, the CD11c^-^ cells displayed a substantially greater proliferation rate than their CD11c^+^ counterparts (Fig. 2G).

Subsequently, we characterized the functional profiles of distinct CBcDC subpopulations. Among the tested TLR ligands, Poly(I:C) induced CBcDC1 to secrete relative low levels of IL-12p70, TNF-α and IFN-λ (fig. S4B). Dual stimulation of CBcDC1 with Poly(I:C) combined with either Pam3CSK4 or R848 further enhanced the production of IL-12p70, TNF-α, IL-6, and IFN-λ, elevating their levels to a substantially greater extent compared with single-agent stimulation (fig. S4B). By contrast, combined stimulation with CpG and Poly(I:C) failed to enhance the production of these cytokines. CBcDC2 secreted IL-6 and IP-10 upon single Poly(I:C) stimulation, whereas only the CD11c^+^ subset of CBcDC2 produced these two cytokines following LPS stimulation (fig. S4B). Additionally, CBcDC2 produced negligible amounts of IL-12p70 and TNF-α upon stimulation with Pam3CSK4+Poly(I:C) or R848+Poly(I:C) combinations—markedly differing from CBcDC1 (fig. S4B). Moreover, both single Poly(I:C) stimulation and Poly(I:C)-based co-stimulation with Pam3CSK4 or R848 upregulated the expression of co-stimulatory molecules in CBcDC1 and CBcDC2 (fig. S4C).

When assessing the cross-presentation capacity of CBcDC1 subpopulations in response to cell-associated antigens, we found that all four subsets could cross-trigger TCR-T cell proliferation. Although CBcDC2 subpopulations exhibited detectable cross-presentation capacity following stimulation with Poly(I:C) and Pam3CSK4, their capacity was markedly weaker than that of CBcDC1 subpopulations (Fig. 2H).

### Establishment of the *in vitro* Ontogeny System for Human cDCs via ScRNA-seq

To conduct a comprehensive and in-depth exploration of the differentiation process of CBcDC, we employed single-cell RNA sequencing (scRNA-seq) technology to monitor the dynamic alterations of cell subpopulations across four different time points. An unbiased dimensionality-reduction clustering analysis identified nine predominant cell types (Fig. 3A-C and table S1 and 2). Specifically, the CBcDC1 cluster predominantly expressed characteristic genes of cDC1, such as *IRF8*, *XCR1*, *THBD*, *CLEC9A*, *BATF3*, *ZNF366*, *CADM1*, etc. (Fig. 3C and D, and fig. S5A). In contrast, the CBcDC2 cluster exhibited high-level expression of hallmarks of cDC2, including *CD1C*, *CD1E*, *CLEC4A*, *SIRPA* and *MRC1* (Fig. 3C and D, and fig. S5A). Notably, both CBcDC1 and CBcDC2 prominently expressed *ZBTB46*, a myeloid cDC marker typically lacking in moDCs (fig. S5A). With respect to the *LAMP3*^+^ DC cluster, in addition to the concurrent expression of marker genes specific to both cDC1 and cDC2, it demonstrated a pronounced expression of two mregDC-characteristic genes, namely *LAMP3* and *CCR7* (Fig. 3C and D, and fig. S5A). Both granulocyte progenitor (GP)_like and neutrophil_like clusters expressed granulocytic lineage genes *MPO*, *CSF3R* and *PRTN3*, with the former manifested higher expression of progenitor genes *CD34*, *PROM1* and *MLLT3* (Fig. 3C and D, and fig. S5A). The megakaryocyte/erythroid progenitors (MEP) cluster was delineated by a substantial expression of TFs *GATA2*, *KLF1*, *TAL1* and *GFI1B*, as well as the hemoglobin-related gene *TFRC* and *HBD* (Fig. 3C and D, and fig. S5A). In contrast, the HSPC cluster was typified by a prominently expression of HSC genes and the absence of lineage specific genes (Fig. 3C and D, and fig. S5A). The DC progenitor cluster was characterized by the expression of *FLT3* and *IRF8*, whereas the *CD1C*^+^ cell cluster predominantly expressed a fraction of cDC2 genes and was likely to be an intermediate precursor cell that bridged DC progenitors and CBcDC2 (Fig. 3C and D, and fig. S5A). As the differentiation progressed, the results clearly demonstrated that the proportions of progenitor cells gradually decreased while that of DCs gradually increased (Fig. 3B).

**Fig. 3.**
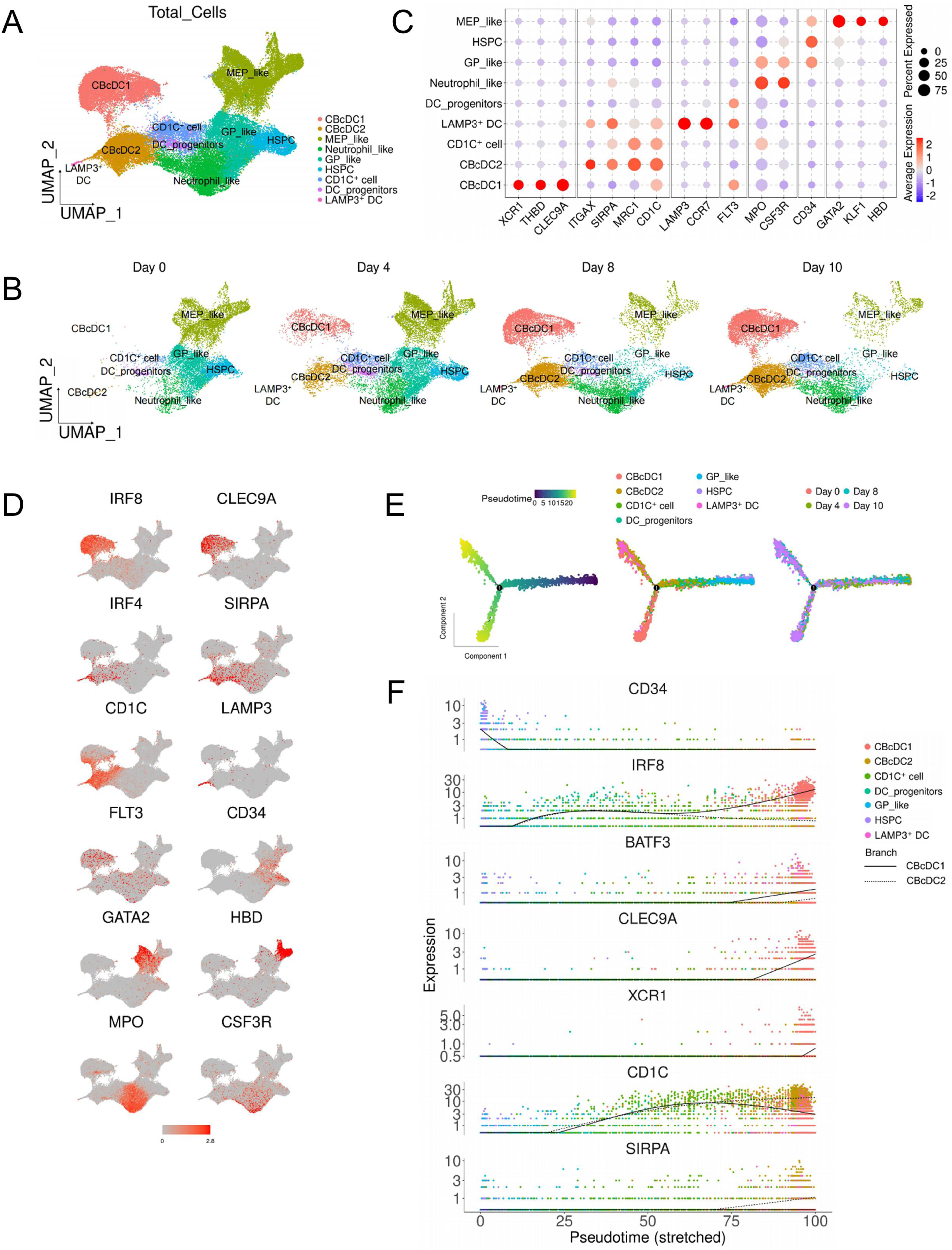
Unbiased scRNA-seq Analysis of Cellular Composition in the Differentiation System at Four Discrete Time Points. (**A**) Unbiased dimensionality reduction clustering identified nine different cell types. The UMAP plot shows the merged data from all four time points. Each point represents a single cell, with the colors denoting different cell types. (**B**) Four UMAP plots illustrate the dynamic changes in cell population composition at different time points during *in vitro* differentiation. Day 0 denotes the time point at the end of pre-expansion and the initiation of differentiation culture. (**C**) Dot plot showing the relative expression levels of marker genes used for cell type clustering. Each dot represents the expression of a specific marker gene within a given cluster. The size of the dot reflects the percentage of cells within the cluster expressing the gene, with larger dots indicating a higher proportion of cells expressing the gene. The color of the dot represents the average expression level of the gene, with red indicating higher expression and blue indicating lower expression. (**D**) UMAP plots depicting the expression of selected genes in each cell. The color scale represents the relative expression levels of these genes. (**E**) Pseudotime trajectory analysis was performed to infer the differentiation pathways of cell populations identified in the *in vitro* differentiation system. Left: A pseudotime trajectory plot illustrating the developmental progression of all cells, where each dot represents a single cell, and colors indicate the estimated pseudotime value. Darker colors correspond to early differentiation stages, while lighter colors indicate later stages. Middle: Cells are color-coded based on their respective cell types, showing the distribution of different cell populations along the inferred trajectory. Right: Cells are colored according to their sampling time points, illustrating their distribution along the inferred trajectory. (**F**) Expression patterns of selected genes along the branched pseudo time axis. The plots show the relative expression of selected genes across two distinct branches. The solid line represents the gene expression trend in CBcDC1, while the dashed line represents the trend in CBcDC2. Dots are colored according to cell type. The y-axis represents the relative expression levels of each gene.

To more comprehensively validate the characteristics of the DC subsets identified by our system, we respectively compared the gene profiles of these DC subsets with those of previously reported DC subsets sourced from human spleens, peripheral blood, and those engendered by different *in vitro* system. The CBcDC1 cluster was demonstrated to project onto the cDC1 and mitotic cDC1 clusters within the human spleen atlas (fig. S5B)(*23*). Concurrently, the CBcDC2 cluster was mapped precisely onto the *CLEC10A*^+^ and *CLEC10A*^-^ cDC2, mitotic cDC2 and ASDC clusters (fig. S5B). Moreover, the *LAMP3*^+^ DC was found to exhibit a similar plotting pattern to the *CCR7*^+^ cDC2 (fig. S5B). Notably, the CBcDC1 and CBcDC2 clusters had achieved high scores for the hallmarks of human peripheral blood cDC1 and cDC2, respectively (fig. S5C)(*24*). And CBcDC1 and CBcDC2 clusters were also demonstrated to be respectively enriched in the characteristic gene signatures of cDC1 and cDC2 stemming from distinct *in vitro* induction system (fig. S5C)(*25*). To provide a more comprehensive characterization, we scored all nine subsets against a panel of gene sets derived from diverse sources. Consistent with their identities, our induced CBcDC1 and CBcDC2 subsets exhibited the highest enrichment scores for cDC1- and cDC2-specific gene signatures, respectively, across all datasets analyzed (fig. S5D)(*24*). Significantly, *LAMP3*^+^ DC exhibited the highest expression of mregDC-characteristic gene set and also expressed cDC1 and cDC2 genes, suggesting a dual origin (fig. S5D)(*26*). *CD1C*^+^ cluster showed a slightly elevated expression of cDC2 signature genes, further validating its closer relation to cDC2 (fig. S5D). DC progenitor cells were found to highly express the gene set associated with early pre-DC, which has been demonstrated to possess the potential for differentiation into cDC1 and cDC2 (fig. S5D). Pseudo time trajectory analysis further confirmed that starting from HSPC, the cells differentiated through DC progenitors and finally developed into CBcDC1 and CBcDC2 (Fig. 3E and fig. S5E). Particularly, although the majority of *CD1C*^+^ cells tended to differentiate into CBcDC2, a small fraction of them differentiated along the CBcDC1 trajectory (Fig. 3E and fig. S5E). This finding clarifies our earlier observation that a small subset of CD11c^-^ CBcDC2 subpopulation, sorted using FACS, has the capacity to differentiate into CBcDC1 (Fig. 2F). From day 0 to day 10, the cell distribution closely followed the pseudo time line from the early to the late stage (Fig. 3E). Upon in-depth analysis of the expression profiles of characteristic genes along the branched pseudo time trajectory, as the expression of *CD34* declined, both *IRF8* and *CD1C* exhibited an initial upregulation in the differentiation branches of CBcDC1 and CBcDC2. Notably, the expression of *IRF8* continued to rise in the CBcDC1 branch, while it decreased in the CBcDC2 branch. Conversely, *CD1C* maintained a high-level expression in the CBcDC2 branch but experienced a reduction in the CBcDC1 branch (Fig. 3F). The expression of cDC1 genes *BATF3*, *CLEC9A*, and *XCR1* were found to be predominantly increased in the late stage of the CBcDC1 branch, and the cDC2 gene *SIRPA* showed exclusive upregulation in the late stage of the CBcDC2 branch (Fig. 3F).

Subsequently, we proceeded to conduct unbiased dimensionality reduction analysis on the CBcDC1 population. A total of five subclusters were identified within the CBcDC1 population (Fig. 4A and B). Specifically, CBcDC1_C0 displayed positive expression for both *CD1C* and *XCR1*, accompanied by a high-level expression of *CD33* and *SPI1*. CBcDC1_C1 was characterized by positive expression of *CD1C* and genes associated with mitosis. CBcDC1_C2 exhibited positive signals for *XCR1*, *FLT3*, and *SPI1*. CBcDC1_C3 was negative for both *CD1C* and *XCR1*, along with a moderate expression of mitosis genes. CBcDC1_C4 was positive for *CD1C*, *CD34*, *KIT*, and a fraction of cDC2-associated genes, representing the characteristics of early progenitors (Fig. 4B and fig. S6A). The expression patterns of cDC1 genes (*XCR1*, *CLEC9A* and *THBD*) across distinct subclusters aligned with those of the *IRF8* gene but deviated from those of the *BATF3* gene (Fig. 4B and fig. S6A). Pseudo time trajectory analysis revealed that the differentiation timeline followed a sequential path from C1 to C3, then to C0, and ultimately to C2, characterized by a transition from *CD1C*^+^ cells to *CD1C*^-^ cells and from *XCR1*^-^ cells to *XCR1*^+^ cells (Fig. 4C and fig. S6B). This finding was further supported by the observations that C1, C3 and C4 subclusters displayed higher scores of pre-cDC1 signature genes (Fig. 4D). More importantly, these outcomes were in alignment with the conclusions drawn from the previously described flow cytometry-based subpopulation discrimination (Fig. 2E and F). Along the differentiation trajectory of CBcDC1 subclusters, the expression of *IRF8* remained high levels, whereas that of *CLEC9A* and *XCR1* gradually increased. In contrast, the expression of *BATF3* showed a gradual decline. As for *CD1C*, its expression initially decreased before increasing (fig. S6C).

**Fig. 4.**
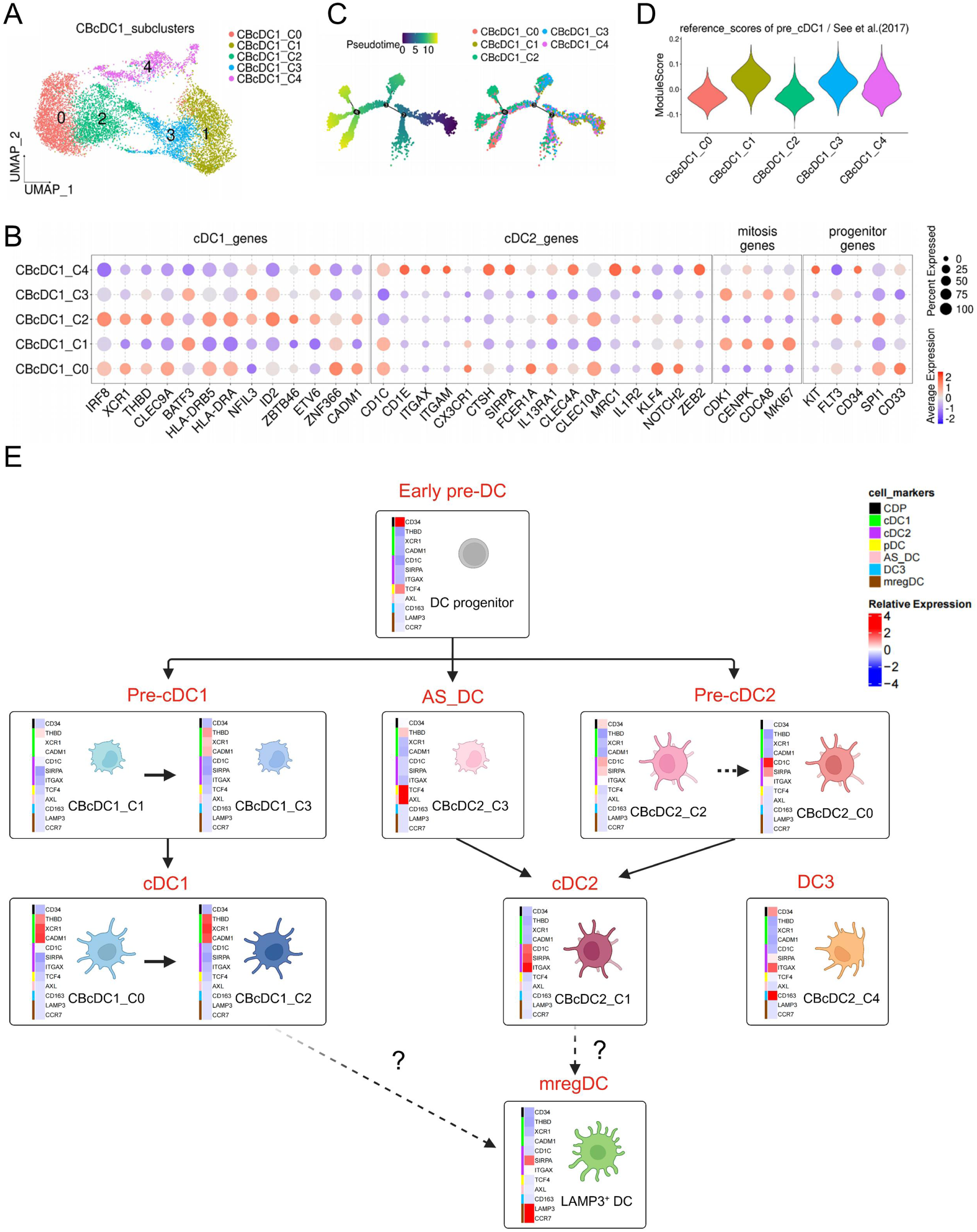
Unbiased Analysis of Cellular Composition within CBcDC1 Subclusters. (**A**) UMAP plot showing unsupervised cell clustering within CBcDC1. (**B**) Dot plot depicting the relative expression levels of selected marker genes across different CBcDC1 subclusters. (**C**) Pseudo-time trajectory analysis illustrating the inferred developmental progression of CBcDC1 subclusters. The left plot shows the pseudo-time trajectory. Right, cells are color-coded by their subcluster identities, showing their distribution along the inferred trajectory. (**D**) Violin plots showing the enrichment scores of human pre-cDC1 reference gene set across the five CBcDC1 subclusters. (**E**) Schematic representation of the hierarchy of cDC development *in vitro* as established in this study. The official names of cell types are indicated in red, while the corresponding subclusters identified in this study are shown in black. The heatmap provides a detailed overview of the relative expression levels of marker genes for CDP, cDC1, cDC2, pDC, ASDC, DC3, and mregDC across the identified clusters.

When conducting unbiased dimensionality-reduction clustering on the CBcDC2 population, we identified five subclusters (fig. S7A-C). The CBcDC2_C1 and CBcDC2_C4 subclusters tested positive for *ITGAX*. Notably, the CBcDC2_C4 subcluster also demonstrated a significantly higher expression level of DC3 signature genes, including *CD163* and *CD14* (fig. S7B and C). In contrast, the CBcDC2_C3 subcluster specifically expressed *AXL*, *SIGLEC6*, *TCF4* and *CX3CR1*, four marker genes for ASDC (fig. S7B and C). Meanwhile, both the CBcDC2_C0 and CBcDC2_C2 subclusters were negative for *ITGAX*. Particularly, the CBcDC2_C2 subcluster showed pronounced expression of mitosis-related genes (fig. S7B and C). *CLEC10A*, a phenotypic marker used to distinguish DC2 from DC3 subsets, was highly expressed in C0 and C1 subclusters but showed lowest expression in C4 (fig. S7B and C). The expression level of *SIRPA* genes was higher in C1 than in C0, whereas the expression level of *FCER1A* was higher in C0 than in C1 (fig. S7B and C). Pseudo time trajectory analysis further elucidated that the differentiation sequence adhered to a sequential progression from C2 to C0 and subsequently to C1 (fig. S7D). This finding was consistent with the conclusions drawn from flow cytometry analysis, which indicated that the *ITGAX*-negative fraction of CBcDC2 served as the precursor of the *ITGAX*-positive fraction of CBcDC2 (Fig. 2E and F). As the expression levels of *CD1C* and *CLEC10A* decline, the expression of *ITGAX* was found to be increased at a later stage (fig. S7E). Furthermore, both the C0 and C1 subclusters exhibited higher scores for the cDC2A signature gene sets. In contrast, the C3 subcluster expressed high levels of ASDC signature genes, and the C4 subcluster showed elevated expression of DC3 signature genes (fig. S7F).

In summary, scRNA-seq analysis demonstrated that we successfully established an *in vitro* ontogeny system for cDCs that closely mimicked the *in vivo* counterpart, comprising early pre-DC, pre-cDC1, pre-cDC2, cDC1, cDC2, DC3, ASDC and mregDC. Leveraging the advantages of this *in vitro* system, we were able to identify more refined DC precursor subclusters and cDC subclusters (Fig. 4E). Starting from the early pre-DC stage, the differentiation into cDC1 via pre-cDC1 was characterized by the progressive upregulation of cDC1-signature genes. Meanwhile, the differentiation into cDC2 via pre-cDC2 showed increasing expression of cDC2-specific markers. The pDC-signature TF *TCF4* is expressed in early pre-DC but lost in mature cDC, suggesting that early pre-DC might retain bipotent differentiation potential towards both cDC and pDC lineages. ASDCs, which are referred to as transitional DCs (tDCs) in some studies and serve as precursors of cDC2s, exhibited a hybrid gene signature combining features of cDC2 (e.g., *KLF4*), pDC (e.g., *TCF4*) and cDC1 (e.g., *THBD*) (Fig. 4E and fig. S7C).

### Allogeneic CBcDC1 Elicits Systemic Antitumor Immunity

Given that we have successfully generated a substantial number of CBcDC1 with potent functionality via *in vitro* system, we then set out to evaluate the anti-tumor potential of these cells by utilizing a humanized mouse model. The NCG (*NOD/ShiLtJGpt-Prkd^cem26Cd52^Il2rg^em26Cd22^/Gpt*) mouse is a severely immunodeficient mouse strain developed by using gene-editing technology to knock out the *Prkdc* (Protein kinase, DNA-activated, catalytic polypeptide) and *Il2rg* (Common γ-chain receptor) genes in *NOD/ShiltJGpt* mouse. The NCG-X mouse, a genetically modified variant of the NCG mouse harboring a homozygous *Kit*^W-41J^ mutation, can support the engraftment of huHSCs without the need for γ-radiation(*27, 28*). To generate humanized NCG-X (huNCG-X) mice, CD34^+^ HSPCs from a healthy donor with the HLA-A*02:01 genotype were injected intravenously into NCG-X juvenile mice. We have validated that the huNCG-X mice manifested remarkable humanization outcomes (fig. S8A-G). Specifically, they recapitulated a diverse array of human immune cell subsets in the spleen, such as B cells, CD4^+^ T cells, CD8^+^ T cells, NK cells, monocytes, cDC1, cDC2, and pDCs (fig. S8A-D). Notably, within the thymus of these mice, the developmental process of human T cell subsets has been successfully reconstituted (fig. S8E). Additionally, both human thymic HLA-A*02-expressing epithelial cells and DCs—key players in T cell clonal selection—were efficiently reconstituted, consistent with prior studies (fig. S8F and G)(*28, 29*).

At the 18th week following hematopoietic reconstitution, a melanoma xenograft model was established via subcutaneous injection of the A375 cell line, which harbors the HLA-A*02:01 genotype. Subsequently, on days 4, 7, and 10 after tumor implantation, intratumoral delivery of pre-activated CBcDC1 with the HLA-A*02:01 genotype was performed in the tumor-bearing mice (Fig. 5A). The assessment of immune responses was carried out on day 16 after tumor inoculation, whereas the observation of tumor growth was prolonged until day 20 post-tumor establishment (Fig. 5A). It was observed that the average tumor size in the CBcDC1 treatment group was approximately 22.3% of that in the control group. Notably, 1/3 of the mice in the treatment group (3 out of 9) experienced complete tumor regression before the end-point of the detection (Fig. 5B). Meanwhile, administration of CBcDC1 also led to a significant improvement in the survival rate of tumor-bearing mice (fig. S9A).

**Fig. 5.**
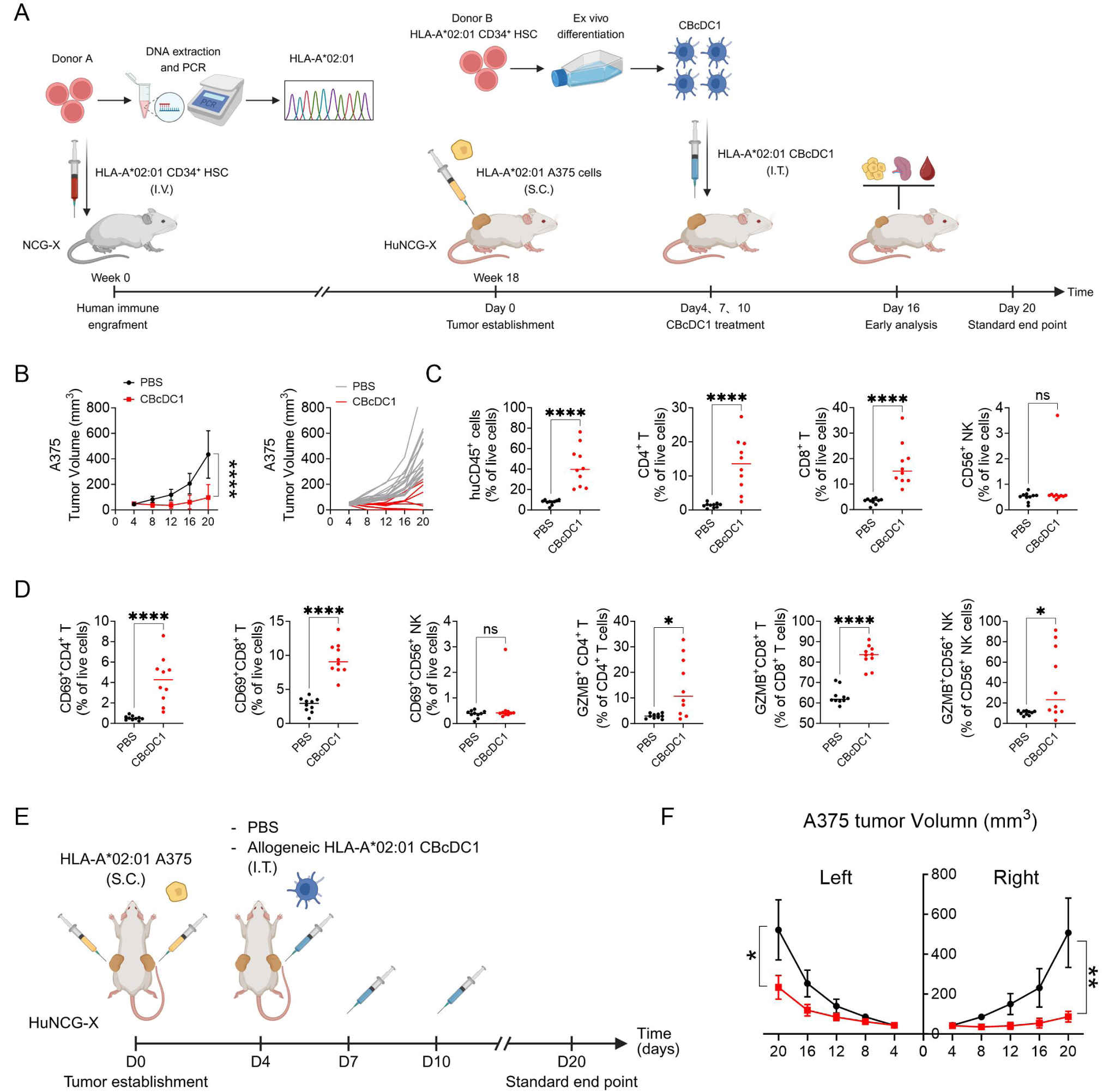
Allogeneic CBcDC1 Elicits Systemic Antitumor Immunity. (**A**) NCG-X mice were intravenously (I.V.) infused with human HLA-A*02:01 CD34^+^ CB HSPC (huNCG-X). Xenograft melanoma models were established by subcutaneously (S.C.) inoculation of the HLA-A*02:01 A375 cell line. Allogeneic HLA-A*02:01 human CBcDC1 (1×10^6^ cells per mouse per injection) were activated with Pam3CSK4 and Poly(I:C) for 4 hours, and subsequently intratumorally (I.T.) injected into mice at 4-, 7- and 10-days following tumor establishment. (**B**) Tumor growth kinetics, showing mean tumor volume (left) and growth trajectories of individual tumors (right). PBS, N=17; CBcDC1, N=9. (**C**, **D**) Frequency of the indicated cell types in A375 tumors on day 16 following tumor establishment. PBS, N=10; CBcDC1, N=10. (**E**) Allogeneic CBcDC1 induced systemic anti-tumor responses. HLA-A*02:01 huNCG-X mice with bilateral A375 tumors were vaccinated with activated allogeneic HLA-A*02:01 CBcDC1 (1×10^6^ cells per mouse per injection) at 4-, 7- and 10-days following tumor establishment. The right-sided tumor was treated with CBcDC1, while the left-sided tumor was intact. (**F**) Growth curves of bilateral tumor are shown. N=6. (**B**) and (**F**) data are presented as the mean ± SD and were analyzed by two-way ANOVA with Bonferroni post-hoc test; (**C**) and (**D**) were analyzed by Mann-Whitney U test. *p<0.05, **p<0.01, ****p<0.0001, ns, no significant.

Subsequently, we carried out a comprehensive exploration of the immunological alterations triggered by CBcDC1 therapy (fig. S9B-D and S10A-D)(*30*). The reconstitution efficiency of human CD45^+^ cells in the spleen was approximately 80%, and no significant difference was observed in the humanization efficiency between the control group and the treatment group (fig. S9B). Robust infiltration of human CD45^+^ cells into the tumor was observed, which included CD4^+^ and CD8^+^ T cells (Fig. 5C). The proportions of both CD69^+^CD4^+^ and CD69^+^CD8^+^ activated T cells were also significantly elevated within the tumors (Fig. 5D). In line with this, the percentages of cytotoxic granzyme B (GZMB)^+^CD4^+^ and GZMB^+^CD8^+^ T cells were elevated following CBcDC1 administration (Fig. 5D). While the overall frequency of NK cells within the tumor remained unaltered, cytotoxic GZMB^+^ NK cells were significantly enriched following treatment with CBcDC1 (Fig. 5C and D). Within the spleen, the percentages of GZMB^+^CD4^+^ T cells, GZMB^+^CD8^+^ T cells and GZMB^+^ NK cells were also elevated (fig. S9B). Consistent with these findings, the levels of IFN-γ, IL-12p70, and IL-6 in the peripheral serum were detected to be elevated following treatment with CBcDC1 (fig. S9C). Meanwhile, an expansion (from 1.44% to 3.03%) of antigen-specific CD8^+^ T cells recognizing the HLA-A*02:01 NY-ESO-1_157-165_ tetramer was noted in the tumors of the treatment group (fig. S9D).

We then proceeded to evaluate the systemic anti-tumor potential of CBcDC1. The A375 cell line was subcutaneously inoculated on both flanks of huNCG-X mice to establish a melanoma model (Fig. 5E). Either PBS or allogeneic HLA-A*02:01 CBcDC1s were delivered through intratumoral injection into the right-sided tumor on days 4, 7, and 10 after tumor inoculation, whereas the contralateral tumor was left intact (Fig. 5E). We observed concurrent suppression of bilateral tumor growth in the treatment group, with a stronger inhibitory effect on right-sided tumors (82.9% vs. 55.2% inhibition, right vs. left) (Fig. 5F).

### CBcDC1 Predominantly Localize to the Tumor Microenvironment *In Situ*

To gain insight into the function of CBcDC1, we investigated the distribution of CBcDC1 and its immediate impact on remodeling the tumor microenvironment. HLA-A*02:01 A375 cells engineered to express GFP were employed to establish the melanoma model (fig. S11A). And, allogeneic HLA-A*02:01 CBcDC1 were activated and stained with CellTrace™ Violet (CTV) to distinguish them from endogenous cDC1. Exogenous CBcDC1 were found to predominantly localize within the tumor and do not spread to other tissue sites (fig. S11B and C). The *in-situ* phagocytosis of tumor-associated antigens by CBcDC1 can also be detected (fig. S11D). Meanwhile, the proportions of activated lymphoid cells, including GZMB^+^IFN-γ^+^CD8^+^ T cells and GZMB^+^CD56^+^ NK cells, were elevated in CBcDC1 treated tumors (fig. S11E). Alongside these phenotypic alterations, the levels of IL-12p70, IFN-γ, IFN-λ and IL-6 in the serum were increased (fig. S11F). Therefore, CBcDC1 exerts a systemic function in ameliorating the tumor microenvironment.

### Allogeneic CBcDC1 remodels the tumor microenvironment

To comprehensively delineate the ameliorative effects of CBcDC1 on the tumor microenvironment, we performed scRNA-seq of intratumoral CD45^+^ cells at 12 days post A375 tumor implantation (Fig. 6A). Equal numbers of sorted human CD45^+^ cells were pooled separately from 5 control mice and 5 CBcDC1-treated mice for scRNA-seq and TCR sequencing. After quality control, a total of 19,006 cells were retained, comprising 8,915 cells in the control group and 10,091 cells in the CBcDC1-treated group (table S3). The cells were dimensionally reduced and clustered into seven major populations according to the common characteristic gene sets (Fig. 6B and fig. S12A). These populations encompassed B cells (n=2,163), CD4^+^ T cells (n=1,494), CD8^+^ T cells (n=11,251), cDC1 (n=455), *LAMP3*^+^ DC (n=154), pDC (n=1,410), monocytes and macrophages (n=2,079) (Fig. 6B and table S3). When compared with the control group, the CBcDC1 treatment group showed a striking increase in the proportions of CD8^+^ T cells (from 40.5% in the control to 75.7% in the treatment) and CD4^+^ T cells (from 3.4% in the control to 11.7% in the treatment). Conversely, there was a significant decline in the relative proportions of B cells (dropping from 16.9% in the control to 6.5% in the treatment) and myeloid cells, with monocytes and macrophages showing a particularly pronounced decrease (from 20.4% in the control to 2.6% in the treatment) (Fig. 6B).

**Fig. 6.**
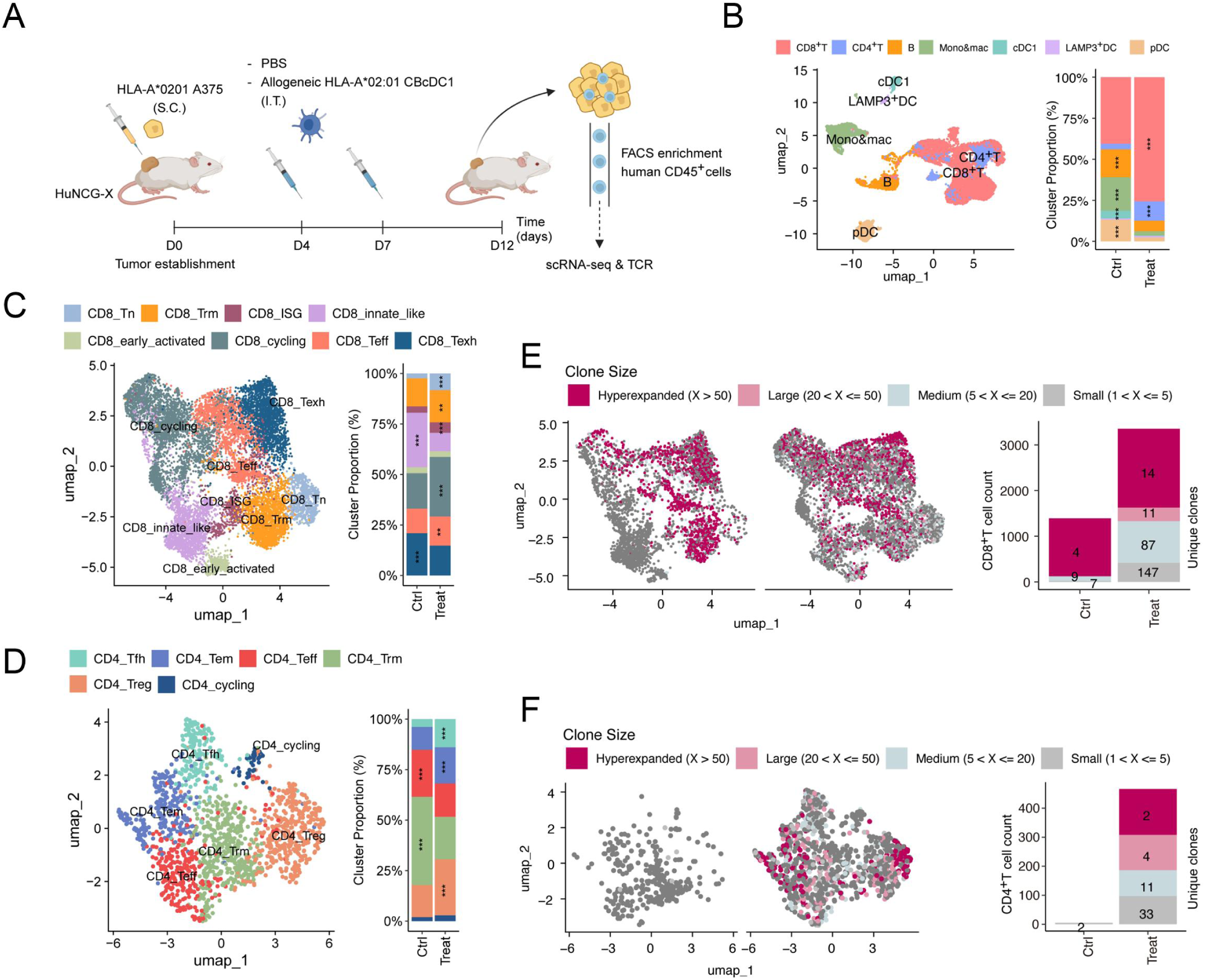
Allogeneic CBcDC1 Remodels the Tumor Microenvironment. (**A**) HLA-A*02:01 A375 tumor-bearing mice were treated with CBcDC1 on the 4th and 7th days following tumor establishment. Single cells were prepared from the tumors of PBS-treated (Ctrl) or HLA-A*02:01 CBcDC1-treated (Treat) HLA-A*02:01 huNCG-X mice on day 12. Subsequently, equal numbers of human CD45^+^ cells from 5 mice in each group were pooled, isolated using FACS, and then subjected to scRNAseq analysis. (**B**, left) Unbiased dimensionality reduction clustering identified different immune cell types. (**B**, right) The proportion of different immune cell types in Ctrl and Treat groups. (**C**, **D**) Unbiased analysis of the composition of the CD8^+^ T cell (**C**) and CD4^+^ T cell (**D**) subclusters. (**E**) UMAP plots illustrating the distribution of TCR clonotypes in CD8^+^ T cells, color-coded by clonotype size: small (2-5 cells), medium (6-20 cells), large (21-50 cells), and hyperexpanded (>50 cells). Single-cell TCR clonotypes were excluded from the analysis. Bar plots depicting the clonotype distribution of CD8^+^ T cells. (**F**) UMAP plots showing the clonotype sizes of CD4^+^ T cell. Bar plots displaying the clonotype distribution of CD4^+^ T cells. (**B**-**D**) data were analyzed using the Pearson’s chi-squared (χ²) test. **p<0.01, ***p<0.001.

Subsequently, the additional dimensionality-reduction analysis was conducted on both CD8⁺ T cells and CD4⁺ T cells (Fig. 6C and D, and fig. S12B and C). Eight distinct cell subclusters were discriminated within the CD8⁺ T cell population and six distinct cell subsets were identified within the CD4⁺ T cell population (Fig. 6C and D, and fig. S12B and C).

Compared with the control group, the CD8^+^ T cell subset in the treatment group exhibited increased proportions of naive T cells (Tn), resident memory T cells (Trm), type I IFN-inducible gene (ISG)-high-expressing T cells, cycling T cells and effector T cells (Teff) (Fig. 6C). In contrast, CBcDC1 therapy reduced the relative proportion of exhausted CD8^+^ T cells (Texh) (Fig. 6C). For the CD4^+^ T cell subset, the treatment group showed elevated relative proportions of follicular helper T cell (Tfh), effector memory T cells (Tem), and regulatory T cells (Treg), whereas the relative proportions of Teff and Trm were decreased (Fig. 6D). Analysis of gene expression further revealed that CBcDC1 administration significantly upregulated the expression levels of *GZMB*, *PRF1*, *IFNG*, *ICOS*, and *PDCD1* across most CD8^+^ T cell subclusters (fig. S12D). *TCF7*—a TF critical for determining the magnitude and durability of anti-tumor immune responses—was found to be upregulated in CD8^+^ Texh and CD8^+^ Tn (fig. S12D). Similarly, *GZMB*, *IFNG*, *TNF*, *ICOS*, and *PDCD1* expression trended upward in CD4^+^ T cell subsets (including Teff, Trm and Tem), albeit without statistical significance (fig. S13A). In-depth phenotypic characterization of expanded Treg subsets within the cDC1-treated cohort revealed a marked enrichment of *IL10*⁺ Tregs—a functionally specialized subset that was otherwise undetectable in control groups (fig. S13B–D). Defined by high expression of effector molecules including *GZMB, IL10, ICOS* and *CXCR3* (fig. S13E), these IL10⁺ Tregs are endowed with potent and robust antitumor activity(*31*). Therefore, these findings indicate that CBcDC1 mediates its antitumor effects, at least in part, via the induction of this distinct Treg subset. More importantly, anti-tumor-associated pathways were comprehensively upregulated across most cell populations in the treatment group. Notably, the cytotoxic T lymphocyte (CTL) pathway, IL-12 pathway, cGAS-STING pathway, and IFN-α, IFN-γ, and type III IFN signaling pathways exhibited substantial enhancement (fig. S13F). To evaluate whether the clonal diversity of T cells was expanded, we analyzed full-length αβ T cell receptor (TCR) sequences from single T cells, where each clonotype was defined as a TCR sequence variant represented by two or more cells. In CBcDC1-treated tumors, clonal diversity was enhanced by over 10-fold in CD8^+^ T cells and 25-fold in CD4^+^ T cells (Fig. 6E and F). Collectively, these findings strongly indicate that CBcDC1 markedly augments T cell clonal diversity and effectively reshapes the tumor microenvironment, shifting it from an immunosuppressive to an immunocompetent state.

To validate the cytotoxic potential of intratumoral CD8^+^ T cells expanded by CBcDC1s, we isolated human CD8+ T cells from tumors of control mice and CBcDC1-treated mice (fig. S14A). These cells were then adoptively transferred into NCG-X recipient mice bearing pre-established A375 tumors (fig. S14A). We observed that CD8^+^ T cells from CBcDC1-treated mice exhibited significantly enhanced antitumor activity, with tumor growth inhibition rates of 45.2% for T cells from control tumors versus 94.1% for those from CBcDC1-treated tumors (fig. S14B). Furthermore, the critical role of CD8^+^ T cells was underscored by the fact that when CD8^+^ T cells were depleted *in vivo*, the anti-tumor activity of allogeneic CBcDC1 cells was notably reduced (fig. S14C-E). In contrast, CD4^+^ T cell ablation only partially impaired the anti-tumor capacity of CBcDC1 (fig. S14C-E).

### CBcDC1 is superior in tumor control than immune checkpoint inhibitor and moDC

We next conducted a comparison of the therapeutic efficacy between CBcDC1 and anti-PD-1 monoclonal antibody (pembrolizumab) within the melanoma model (fig. S15A). Monotherapy with pembrolizumab inhibited tumor volume growth by 41.8%. In contrast, CBcDC1 treatment achieved an 80.1% reduction in tumor volume. Significantly, the concomitant administration of pembrolizumab and CBcDC1 led to an 87.3% suppression of tumor growth (fig. S15B).

To assess whether CBcDC1 possesses superior tumor-controlling capacity compared to moDCs, allogeneic HLA-A*02:01 CBcDC1 and moDC were generated *in vitro*. Subsequently, CBcDC1s were activated with Pam3CSK4 and Poly(I:C), whereas moDCs were stimulated with LPS—with both DC subsets pulsed with tumor lysate during the activation process. It was found that treatment with moDCs led to an average 43.4% suppression of tumor volume. In contrast, CBcDC1-based treatment achieved a remarkable 76.5% suppression rate of tumor volume (fig. S15C).

### CBcDC1 possesses comprehensive anti-tumor capabilities in both breast cancer and ovarian cancer

To investigate if CBcDC1 has pan-antitumor capacity, we built breast and ovarian cancer xenograft models in huNCG-X mice, utilizing HLA-A*02:01 MDA-MB-231 cell line and HLA-A*02:01 SKOV3 cell line respectively. Pre-activated HLA-A*02:01 CBcDC1s were administered intratumorally on days 4, 7, and 10 following tumor establishment. It was discovered that the application of CBcDC1 led to a significant 74.4% reduction in breast-tumor volume. Remarkably, complete tumor regression was achieved in 1/3 (2 out of 6) of the mice in treatment group (Fig. 7A). Alongside this phenotype, there was an increase in the intratumoral infiltration of CD45^+^ cells, including CD4^+^ T cells, CD8^+^ T cells and NK cells (Fig. 7B-E). The activation of these lymphocytes also increased in the treatment group, as evidenced by the elevated proportion of the CD69^+^ cell subset (Fig. 7F-H). In addition, there were remarkable augmentations in the percentages of IFN-γ^+^CD4^+^ T cells, TNF-α^+^CD4^+^ T cells and GZMB^+^CD8^+^ T cells (Fig. 7I-K).

**Fig. 7.**
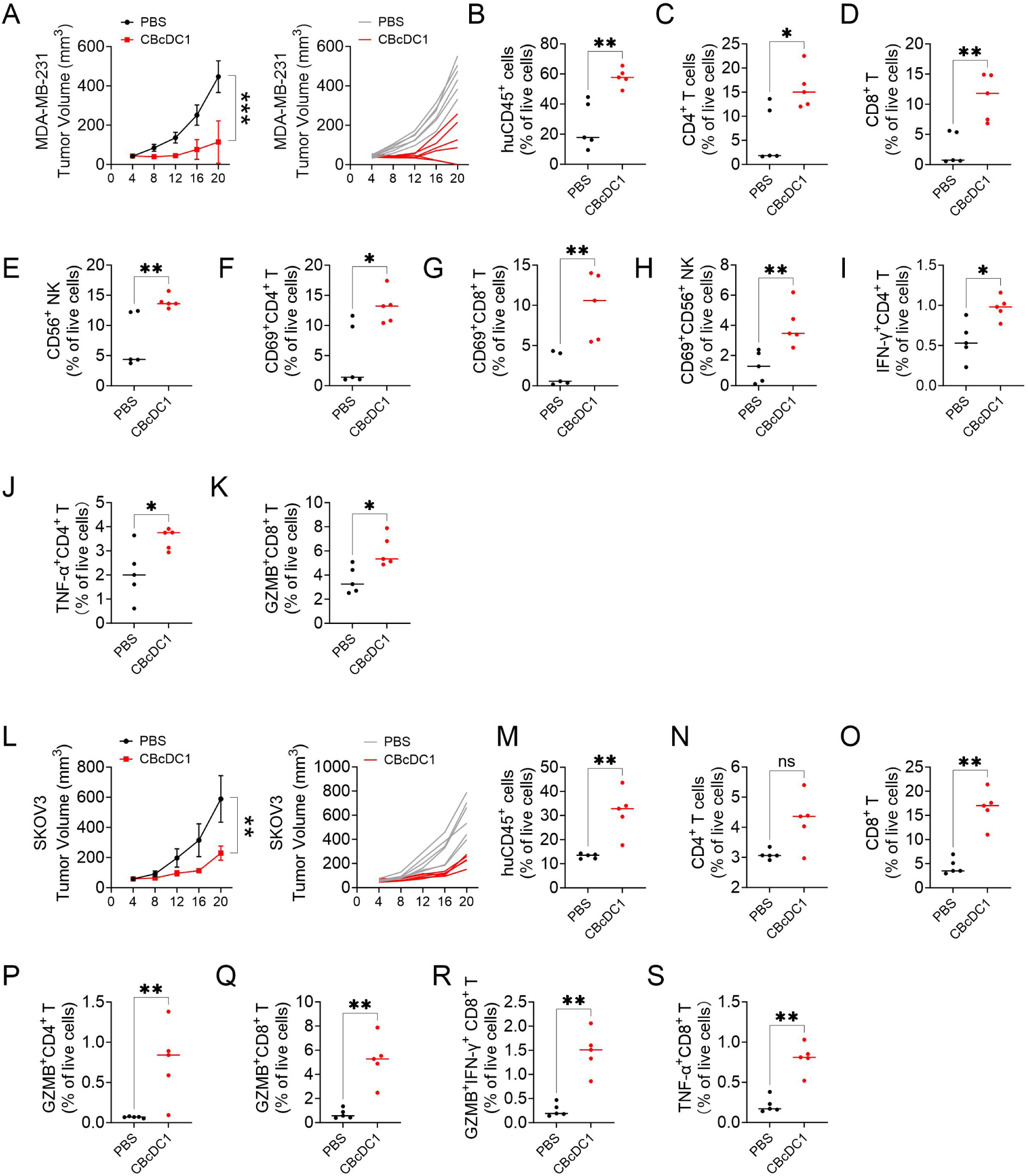
CBcDC1 Possesses Comprehensive Anti-tumor Capabilities in Breast Cancer and Ovarian Cancer. (**A**) Breast cancer models were established by S.C. injection of HLA-A*02:01 MDA-MB-231 cells. Activated allogeneic HLA-A*02:01 CBcDC1 (1×10^6^ cells per mouse per injection) were I.T. injected into mice model at 4-, 7- and 10-days following tumor establishment. Tumor growth kinetics, showing mean tumor volume (left) and growth trajectories of individual tumors (right). N=6. (**B**-**K**) Frequency of the indicated cell types in the tumor at 16 days following tumor establishment. N=5. (**L**) Ovarian cancer models were established by S.C. injection of HLA-A*02:01 SKOV3 cell line. Activated allogeneic HLA-A*02:01 CBcDC1 (1×10^6^ cells per mouse per injection) were I.T. injected into mice model at 4-, 7- and 10-days following tumor establishment. SKOV3 tumor growth curves showing mean tumor size (left) and growth of individual tumors (right). PBS, N=6; CBcDC1, N=5. (**M**-**S**) Frequency of the indicated cell types in the tumor of at 16 days following tumor establishment. N=5. (**A**) and (**L**) data are presented as the mean ± SD and were analyzed by two-way ANOVA with Bonferroni post-hoc test; (**B**-**K**) and (**M**-**S**) were analyzed by Mann-Whitney U test. *p<0.05, **p<0.01, ***p<0.001, ns, no significant.

Similarly, CBcDC1 is capable of exerting an analogous anti-tumor effect within the established ovarian cancer model. Compared with the control group, the tumor volume in the treatment group was reduced by an average of 61.1% (Fig. 7L). Meanwhile, an increase in the intratumoral infiltration of CD45^+^ cells, including CD8^+^ T cells, was observed post CBcDC1 inoculation (Fig. 7M-O). Specifically, there were significant elevations in the proportions of GZMB⁺CD4⁺ T cells, GZMB⁺CD8⁺ T cells, GZMB⁺IFN-γ⁺CD8⁺ T cells, and TNF-α⁺CD8⁺ T cells (Fig. 7P-S). These findings strongly suggests that CBcDC1 exhibits comprehensive anti-tumor capabilities.

## Discussion

In this study, we established a highly efficient and straightforward approach for the generation of human cDCs, specifically cDC1, through the utilization of GMP-grade reagents, offering the prerequisites for the large-scale manufacturing and clinical application of cDC1. Furthermore, we corroborated that these allogeneic cDC1s are distinguished by a remarkably comprehensive anti-tumor effectiveness, which substantially outperforms those of moDC and PD-1 antibody.

The *in vitro* differentiation system of murine DCs has been well-developed. Through the exclusive application of FLT3L, large amounts of both cDC and pDC subsets can be concurrently generated(*32*). However, FLT3L alone can only induce an extremely low proportion and a minuscule number of human DCs. Relative to the initial stem cell count, a 21-day differentiation in Yssel’s medium supplemented with 10% human AB serum, FLT3L and TPO yielded simultaneous expansions of cDC1 (≈2.9-fold), CD1c^+^ DC (≈5.67-fold) and pDC (≈4.79-fold) from G-CSF-mobilized adult bone marrow HSPCs, as reported by Proietto et al(*33*). In contrast, after 7 days of pre-amplification in Stempan medium supplemented with SCF, FLT3L, IL-3, and TPO, followed by 11 days of induced differentiation in RPMI 1640 medium containing SCF, FLT3L, GM-CSF, IL-4, and 10% FCS, Balan et al. obtained approximately 18-fold expansion of XCR1^+^ cDC1 relative to the number of CD34^+^ CB stem cells(*16*). Afterward, the same group further employed murine OP9 feeder cells overexpressing the Notch ligand to enhance the yield of cDC1 and pDC(*25*). Here, we achieved a higher expansion efficiency (average 705-fold, up to 1,614-fold) via a feeder cell-free system containing SR1 and IFN-γ within 2 weeks. SR1 played a crucial role in sustaining the continuous proliferation of stem cells throughout the expansion and differentiation stages, whereas IFN-γ substantially promoted the differentiation towards cDC1 by activating IRF-8. Therefore, when starting with one unit of cord blood containing an initial count of 5×10^6^ CD34^+^ stem cells, we can obtain 1-8×10^9^ (average of 3.5×10^9^ cells) CBcDC1 cells. Given that a typical dosage of DC vaccine is set at 5×10^6^ cells, a single unit of cord blood is sufficient to support 200-1600 treatment courses, with an average of roughly 700 treatments. Notably, our strategy achieves high amplification efficiency via a convenient and readily implementable method. While our findings clearly demonstrate that targeted overexpression of *BATF3* or knockout of *ZEB2* markedly enhances the induction efficiency of cDC1s to over 87% and 60%, respectively, a comprehensive evaluation of the potential adverse effects and increased costs associated with genetic modification reveals that our current strategy is fully capable of meeting the stringent requirements for the clinical translation of cDC1s(*34*).

Recent advances leveraging scRNA-seq have expanded our understanding of DC subset heterogeneity and established a paradigm for differentiation of human DCs. The cDC1 and cDC2 subsets are originated from CDP. Specifically, the cDC1 subset differentiates via the pre-cDC1 stage, and the cDC2 subset via the pre-cDC2 stage, both strictly dependent on FLT3L(*1, 3*). Conversely, DC3 derives from a DC3-committed progenitor within the CLEC12A^+^ fraction of granulocyte, monocyte, and dendritic cell progenitor (GMDP)(*7*). DC3 development acutely relies on GM-CSF and occurs independently of CDP and the monocyte-restricted progenitor (cMoP)(*7*). ASDCs share transcriptional programs with both pDCs and cDCs, and have been confirmed to be cDC2 precursors(*35*). Our findings further reveal that the *in vitro* differentiation system employed herein exhibits remarkable congruence with the endogenous DC differentiation pathways. Moreover, in contrast to *ex vivo* single-cell analyses of human specimens—limited by the number of obtainable cells—this *in vitro* system enables the resolution of finer details of the differentiation process. Significantly, our observations demonstrate that gene editing of specific TFs elicits corresponding alterations in the differentiation trajectories of DC subsets, which closely recapitulates the results from murine studies. Therefore, these findings provide a robust platform for comprehensive explorations of the underlying mechanisms governing the differentiation processes of DC subsets, especially the human-specific regulatory modulators, including enhancers and non-coding RNAs.

The challenge of generating large quantities of cDCs has constrained their translational and clinical application. Consequently, while substantial clinical data have been reported for monocyte-derived dendritic cells (moDCs), the therapeutic efficacy of cDCs in cancer patients remains largely underexplored. Prior studies leveraging FLT3L have provided preliminary evidence supporting the antitumor activity of cDC1s(*36–38*). Furthermore, several research groups have harnessed autologous cDC1s and/or cDC2s — directly isolated from peripheral blood — for therapeutic interventions, yielding early evidence of antitumor activity in immune checkpoint inhibitor-refractory melanoma patients(*39*). More recently, a strategy that directly reprograms tumor cells into cDC1 *in situ* via genetic modification has demonstrated efficacy in murine models(*40*). This approach circumvented the *in vitro* production procedure of cDC1 and was anticipated to offer prompt treatment options for cancer patients. In contrast, our method presents a sophisticated therapy featuring precisely quantifiable dosing tailored for cDC1-based treatment. Moreover, the application of allogeneic cDC1 has surmounted the disadvantages of exorbitant costs and inconvenience associated with autologous cell therapies. The preparation procedure that eschews the utilization of feeder cells and genetic manipulation also ensures the utmost safety of the treatment. Nevertheless, it should be noted that the host-versus-graft-reactions, which are potentially induced by allogeneic cDC1, will constrain the persistence and effectiveness of the cDC1 therapy. This limitation might be mitigated through more precise HLA matching, or alternatively, by engineering the knockout of endogenous HLA molecules in allogeneic cDC1s followed by direct loading with patient-derived tumor peptide-HLA complexes to prevent rejection. This approach leverages a well-characterized MHC cross-dressing mechanism that enables direct activation of CD8^+^ T cells independent of intracellular antigen processing(*41, 42*).

The anti-tumor mechanism of cDC1 has been intensively explored. It shuttles between the tumor and the draining lymph nodes, maintaining a reservoir of proliferative tumor-antigen specific TCF-1^+^CD8^+^ T cells, which drive the response to immune checkpoint blockade(*43*). Moreover, the immunostimulatory CCR7^-^ cDC1 in tumors provides local niches for expansion and effector differentiation of tumor-infiltrating stemlike TCF-1^+^CD8^+^ T cells. Meanwhile, the early priming of CD4^+^ T cells against tumor-derived antigens is also dependent on cDC1s, which sustain robust antitumor responses during both primary and secondary challenges(*11*). Our flow cytometry analyses and single-cell sequencing data demonstrate that CBcDC1-based therapy elicits robust T cell infiltration and expansion within tumors, with a notable predominance of effector CD8^+^ T cells and *TCF1*^+^ CD8⁺ T cells. Furthermore, CD8^+^ T cells demonstrated substantially elevated expression of anti-tumor factors relative to their CD4⁺ T cell counterparts. In line with these observations, *in vivo* depletion of CD8⁺ T cells nearly abrogated the anti-tumor efficacy of CBcDC1, while depletion of CD4⁺ T cells only partially impaired its anti-tumor function. Our proposed mechanism is distinct from the direct cDC1 reprogramming strategies, which highlight the critical role of CD4⁺ T cells in their therapeutic approach(*40*).

cDC1s have been demonstrated to exhibit superior anti-tumor efficacy compared with moDCs and cDC2s(*15, 44*). This advantage is attributed to its specific expression of genes including *CLEC9A*, *WDFY4*, and *XCR1*, which endow cDC1 with a stronger capacity to take up, process, and cross-present cell-associated antigens for T cell activation relative to both moDCs and cDC2s(*12, 45–47*). Furthermore, the *BATF3* gene—specifically and highly expressed in cDC1—confers additional anti-tumor benefits through a cross-presentation-independent tumor rejection mechanism(*48*). Notably, the functional efficacy of widely used moDC vaccines has been confirmed to rely on host endogenous cDC1 cells(*12*). In our study, we observed that CBcDC1 subsets generated in our system possess a more robust ability to activate CD8^+^ T cells compared with CBcDC2 subsets. Concomitantly, CBcDC1s exert a more prominent inhibitory effect on tumor growth than moDCs. Additionally, the induced CBcDC1 subsets retain proliferative capacity, implying their potential for sustained effects that exceed those of terminally differentiated endogenous cDC1s.

The antitumor activity of cDC1-based monotherapy or its combinatorial regimens has been confirmed in a variety of murine tumor models, including melanoma, osteosarcoma, colon adenocarcinoma, immunogenic fibrosarcoma and pancreatic ductal adenocarcinoma (PDAC)(*12, 15, 22, 44, 49*). Building on our preclinical findings, we have recently initiated a clinical trial investigating CBcDC1-mediated therapy for solid tumors. The conduct of this trial is expected to yield more precise data on the therapeutic efficacy and underlying mechanisms of universal cDC1s, which in turn will address the limitations inherent in humanized mouse experiments. More interestingly, dysregulated cDC1 function has been identified to correlate with immune microenvironment perturbations and impaired antitumor potential induced by aging(*50*). Restoring the functional capacity of cDC1s could contribute to the recovery of antitumor vitality in elderly individuals. This finding further substantially broadens the application scope of cDC1-based strategies.

## References and Notes

1. S. L. Nutt, M. Chopin, Transcriptional Networks Driving Dendritic Cell Differentiation and Function. Immunity 52, 942–956 (2020).

2. S. K. Wculek et al., Dendritic cells in cancer immunology and immunotherapy. Nat Rev Immunol 20, 7–24 (2020).

3. A. Schlitzer et al., Identification of cDC1- and cDC2-committed DC progenitors reveals early lineage priming at the common DC progenitor stage in the bone marrow. Nat Immunol 16, 718–728 (2015).

4. J. Lee et al., Restricted dendritic cell and monocyte progenitors in human cord blood and bone marrow. J Exp Med 212, 385–399 (2015).

5. R. J. Dress et al., Plasmacytoid dendritic cells develop from Ly6D(+) lymphoid progenitors distinct from the myeloid lineage. Nat Immunol 20, 852–864 (2019).

6. Z. Liu et al., Dendritic cell type 3 arises from Ly6C(+) monocyte-dendritic cell progenitors. Immunity 56, 1761–1777.e1766 (2023).

7. P. Bourdely et al., Transcriptional and Functional Analysis of CD1c(+) Human Dendritic Cells Identifies a CD163(+) Subset Priming CD8(+)CD103(+) T Cells. Immunity 53, 335–352.e338 (2020).

8. E. Kvedaraite, F. Ginhoux, Human dendritic cells in cancer. Sci Immunol 7, eabm9409 (2022).

9. J. P. Böttcher et al., NK Cells Stimulate Recruitment of cDC1 into the Tumor Microenvironment Promoting Cancer Immune Control. Cell 172, 1022–1037.e1014 (2018).

10. S. Spranger, D. Dai, B. Horton, T. F. Gajewski, Tumor-Residing Batf3 Dendritic Cells Are Required for Effector T Cell Trafficking and Adoptive T Cell Therapy. Cancer Cell 31, 711–723.e714 (2017).

11. S. T. Ferris et al., cDC1 prime and are licensed by CD4(+) T cells to induce anti-tumour immunity. Nature 584, 624–629 (2020).

12. S. T. Ferris et al., cDC1 Vaccines Drive Tumor Rejection by Direct Presentation Independently of Host cDC1. Cancer Immunol Res 10, 920–931 (2022).

13. S. Zhang, M. Chopin, S. L. Nutt, Type 1 conventional dendritic cells: ontogeny, function, and emerging roles in cancer immunotherapy. Trends Immunol 42, 1113–1127 (2021).

14. P. Meiser et al., A distinct stimulatory cDC1 subpopulation amplifies CD8(+) T cell responses in tumors for protective anti-cancer immunity. Cancer Cell 41, 1498–1515.e1410 (2023).

15. Y. Zhou et al., Vaccine efficacy against primary and metastatic cancer with in vitro-generated CD103(+) conventional dendritic cells. J Immunother Cancer 8, (2020).

16. S. Balan et al., Human XCR1+ dendritic cells derived in vitro from CD34+ progenitors closely resemble blood dendritic cells, including their adjuvant responsiveness, contrary to monocyte-derived dendritic cells. J Immunol 193, 1622–1635 (2014).

17. A. E. Boitano et al., Aryl hydrocarbon receptor antagonists promote the expansion of human hematopoietic stem cells. Science 329, 1345–1348 (2010).

18. F. F. Rosa, et al., Single-cell transcriptional profiling informs efficient reprogramming of human somatic cells to cross-presenting dendritic cells. Sci Immunol 7, eabg5539 (2022).

19. G. Schiavoni et al., ICSBP is essential for the development of mouse type I interferon-producing cells and for the generation and activation of CD8alpha(+) dendritic cells. J Exp Med 196, 1415–1425 (2002).

20. T. Lança et al., IRF8 deficiency induces the transcriptional, functional, and epigenetic reprogramming of cDC1 into the cDC2 lineage. Immunity 55, 1431–1447.e1411 (2022).

21. P. H. Driggers et al., An interferon gamma-regulated protein that binds the interferon-inducible enhancer element of major histocompatibility complex class I genes. Proc Natl Acad Sci U S A 87, 3743–3747 (1990).

22. U. Cytlak et al., Differential IRF8 Transcription Factor Requirement Defines Two Pathways of Dendritic Cell Development in Humans. Immunity 53, 353–370.e358 (2020).

23. C. C. Brown et al., Transcriptional Basis of Mouse and Human Dendritic Cell Heterogeneity. Cell 179, 846–863.e824 (2019).

24. P. See et al., Mapping the human DC lineage through the integration of high-dimensional techniques. Science 356, (2017).

25. S. Balan et al., Large-Scale Human Dendritic Cell Differentiation Revealing Notch-Dependent Lineage Bifurcation and Heterogeneity. Cell Rep 24, 1902–1915.e1906 (2018).

26. K. Mulder et al., Cross-tissue single-cell landscape of human monocytes and macrophages in health and disease. Immunity 54, 1883–1900.e1885 (2021).

27. K. N. Cosgun et al., Kit regulates HSC engraftment across the human-mouse species barrier. Cell Stem Cell 15, 227–238 (2014).

28. D. P. Chupp et al., A humanized mouse that mounts mature class-switched, hypermutated and neutralizing antibody responses. Nat Immunol 25, 1489–1506 (2024).

29. S. Chakrabarti et al., Bone Marrow-Derived Cells Contribute to the Maintenance of Thymic Stroma including TECs. J Immunol Res 2022, 6061746 (2022).

30. A. Saxena, P. K. Dagur, A. Biancotto, Multiparametric Flow Cytometry Analysis of Naïve, Memory, and Effector T Cells. Methods Mol Biol 2032, 129–140 (2019).

31. X. Huang et al., Opposing functions of distinct regulatory T cell subsets in colorectal cancer. Immunity, (2025).

32. S. H. Naik et al., Cutting edge: generation of splenic CD8+ and CD8-dendritic cell equivalents in Fms-like tyrosine kinase 3 ligand bone marrow cultures. J Immunol 174, 6592–6597 (2005).

33. A. I. Proietto, D. Mittag, A. W. Roberts, N. Sprigg, L. Wu, The equivalents of human blood and spleen dendritic cell subtypes can be generated in vitro from human CD34(+) stem cells in the presence of fms-like tyrosine kinase 3 ligand and thrombopoietin. Cell Mol Immunol 9, 446–454 (2012).

34. T. T. Liu et al., Ablation of cDC2 development by triple mutations within the Zeb2 enhancer. Nature 607, 142–148 (2022).

35. A. C. Villani et al., Single-cell RNA-seq reveals new types of human blood dendritic cells, monocytes, and progenitors. Science 356, (2017).

36. N. Bhardwaj et al., Flt3 ligand augments immune responses to anti-DEC-205-NY-ESO-1 vaccine through expansion of dendritic cell subsets. Nat Cancer 1, 1204–1217 (2020).

37. L. Hammerich et al., Systemic clinical tumor regressions and potentiation of PD1 blockade with in situ vaccination. Nat Med 25, 814–824 (2019).

38. G. D. Hogg et al., Combined Flt3L and CD40 agonism restores dendritic cell-driven T cell immunity in pancreatic cancer. Sci Immunol 10, eadp3978 (2025).

39. J. K. Schwarze et al., Intratumoral administration of CD1c (BDCA-1)(+) and CD141 (BDCA-3)(+) myeloid dendritic cells in combination with talimogene laherparepvec in immune checkpoint blockade refractory advanced melanoma patients: a phase I clinical trial. J Immunother Cancer 10, (2022).

40. E. Ascic et al., In vivo dendritic cell reprogramming for cancer immunotherapy. Science 386, eadn9083 (2024).

41. E. Duong et al., Type I interferon activates MHC class I-dressed CD11b(+) conventional dendritic cells to promote protective anti-tumor CD8(+) T cell immunity. Immunity 55, 308–323.e309 (2022).

42. B. W. MacNabb et al., Dendritic cells can prime anti-tumor CD8(+) T cell responses through major histocompatibility complex cross-dressing. Immunity 55, 982–997.e988 (2022).

43. J. M. Schenkel et al., Conventional type I dendritic cells maintain a reservoir of proliferative tumor-antigen specific TCF-1(+) CD8(+) T cells in tumor-draining lymph nodes. Immunity 54, 2338–2353.e2336 (2021).

44. I. Heras-Murillo et al., Immunotherapy with conventional type-1 dendritic cells induces immune memory and limits tumor relapse. Nat Commun 16, 3369 (2025).

45. D. J. Theisen et al., WDFY4 is required for cross-presentation in response to viral and tumor antigens. Science 362, 694–699 (2018).

46. D. Sancho et al., Identification of a dendritic cell receptor that couples sensing of necrosis to immunity. Nature 458, 899–903 (2009).

47. B. G. Dorner et al., Selective expression of the chemokine receptor XCR1 on cross-presenting dendritic cells determines cooperation with CD8+ T cells. Immunity 31, 823–833 (2009).

48. D. J. Theisen et al., Batf3-Dependent Genes Control Tumor Rejection Induced by Dendritic Cells Independently of Cross-Presentation. Cancer Immunol Res 7, 29–39 (2019).

49. K. K. Mahadevan et al., Type I conventional dendritic cells facilitate immunotherapy in pancreatic cancer. Science 384, eadh4567 (2024).

50. A. C. Y. Chen et al., The aged tumor microenvironment limits T cell control of cancer. Nat Immunol 25, 1033–1045 (2024).

51. Q. Pan et al., Phase 1 clinical trial to assess safety and efficacy of NY-ESO-1-specific TCR T cells in HLA-A∗02:01 patients with advanced soft tissue sarcoma. Cell Rep Med 4, 101133 (2023).

