## Supplementary material for "Scalable Generation of Clinical-Grade Universal Human cDC1s Enables Potent Antitumor Immunotherapy": fig. S1-S15, supplemental Table 1-5

### Materials and Methods

#### Mice

NCG-X mice（Strain NO. T003802）were purchased from GemPharmatech (Nanjing, China). Humanized mice were generated by grafting 3-week-old NCG-X mice with $CD34^+$ HSPC. Each mouse received $2\times10^5$ freshly isolated or frozen-thawed $huCD34^+$ cells that carried the HLA-A*02:01 genotype. All the mice employed in the experiments were housed in a specific-pathogen-free (SPF) animal facility at Renji Hospital.

#### Human Samples

$HuCD34^+$ hematopoietic stem cells (HSCs) were carefully isolated from umbilical cord blood of full-term, normal male and female newborns delivered by cesarean section, after getting informed consent from healthy puerperae of various ages. Human peripheral blood derived-DC subsets were isolated from healthy adult human donors. All human experiments were conducted in accordance with the declaration of Helsinki Principles. Written informed consent was obtained from participants prior to inclusion in the study. Human studies were approved by the ethics committee of Renji Hospital.

The $CD34^+$ cells were purified using CliniMACS® CD34 Reagent CR/GMP (220-001-831, Miltenyi Biotech) according to the manufacturer's instructions, yielding at least 99% purity $huCD34^+$ cell preparations. Immediately after purification, the freshly isolated $huCD34^+$ cells underwent HLA genotyping. Subsequently, cells with the HLA-A*02:01 genotype were resuspended in stem cell expansion medium for the induction of cDC1 cells; or cryopreserved in CryoStor CS10 (210102, Biolife Solutions) and stored in liquid nitrogen for later applications. Human primary $CD8^+$ T cells (HLA-A*02-negative) were isolated from PBMC using the human naive $CD8^+$ T cell isolation kit (130-093-244, Miltenyi Biotec).

#### Cell culture

The A375, SKOV3 and MDA-MB-231 cell lines were obtained from the National Collection of Authenticated Cell Cultures. The A375 cell line was cultivated in Dulbecco's Modified Eagle Medium (DMEM, Thermofisher), supplemented with 10% FBS (Thermofisher) and 1% penicillin-streptomycin (PS, Thermofisher). The SKOV3 cell line was maintained in McCOY's 5A medium (Thermofisher), with the addition of 10% FBS (Thermofisher) and 1% PS (Thermofisher). The MDA-MB-231 cell line was grown in Leibovitz L-15 medium (Thermofisher), which was supplemented with 10% FBS (Thermofisher) and 1% PS (Thermofisher). Human primary $CD8^+$ T cells were cultured in RPMI 1640 medium, supplemented with 10% FBS (Thermofisher), 1% PS and 100ng/ml huIL-2 (BT-002-GMP, R&D systems).

#### DC induction, isolation and activation

$HuCD34^+$ cells were first cultured in GMP-compliant StemSpan™-AOF medium (#100-0130, STEMCELL Technologies) with 100 ng/ml Recombinant Human SCF GMP Protein (BT-SCF-GMP, R&D systems), 100 ng/ml Recombinant Human Thrombopoietin/TPO GMP Protein (BT-TPO-GMP, R&D systems), 100 ng/ml Recombinant Human Flt-3 Ligand/FLT3L GMP Protein (BT-FT3L-GMP, R&D systems), and 1 μM SR1 (#72342, STEMCELL Technologies) for 4 days. Then, they were transferred to fresh StemSpan™-AOF medium supplemented with 100 ng/ml

SCF, 100 ng/ml FLT3L, 1 μM SR1, 5 ng/ml Recombinant Human IFN-gamma GMP Protein (285-GMP, R&D systems), 2.5 ng/ml Recombinant Human GM-CSF GMP Protein (215-GMP, R&D systems), and 2.5 ng/ml Recombinant Human IL-4 GMP Protein (BT-004-GMP, R&D systems) for 10 days, with the medium refreshed every 4 days. Regarding to X-VIVO 15 conditions, the StemSpan™-AOF medium was replaced with the Lonza X-VIVO 15 medium (BEBP02-061Q, Lonza Bioscience). For the purpose of treating the tumor model, CBcDC1 cell were purified with the CliniMACS® CD141 (BDCA-3) Reagent CR/GMP kit (220-001-832, Miltenyi Biotech) at a dosage twice the amount recommended in the instruction manual. The purity of CBcDC1 is higher than 98%. Then, CBcDC1 underwent a 6-hour stimulation process with 1 μg/ml Poly(I:C) (tlrl-pic, Invivogen) and 1 μg/ml Pam3CSK4 (tlrl-pms, Invivogen), and subsequently, these activated cells were applied to tumor treatment. MoDCs were generated by culturing primary monocytes in RPMI 1640 medium supplemented with 100 ng/ml Recombinant Human GM-CSF GMP Protein (215-GMP, R&D systems) and 20 ng/ml Recombinant Human IL-4 GMP Protein (BT-004-GMP, R&D systems) for 5 days. Subsequently, moDCs ($CD11c^+HLA\text{-}DR^+$) were isolated, stimulated with 100 ng/ml LPS (tlrl-eblps; Invivogen) for 12 hours, and pulsed with tumor lysate concomitantly during activation. Peripheral blood DC subsets were isolated via FACS based on the following immunophenotypic markers: PBcDC1, $Lin^-CD123^-AXL^-HLA\text{-}DR^+CD11c^+CD1c^-SIRP\alpha^-CD141^+CLEC9A^+$; PBcDC2, $Lin^-CD123^-AXL^-HLA\text{-}DR^+CD11c^+CD1c^+SIRP\alpha^+CD141^-CLEC9A^-$; PBpDC, $Lin^-AXL^-HLA\text{-}DR^+CD11c^-CD1c^-SIRP\alpha^-CD141^-CLEC9A^-CD123^+BDCA2^+$.

During the investigation of the subpopulations, both the CBcDC1 ($Lin^-CD123^-AXL^-CD141^+$) and CBcDC2 ($Lin^-CD123^-AXL^-CD141^{low/-}CD1c^{high}$) subclusters were isolated using FACS. Then, they were stimulated either individually with 1 μg/ml Poly(I:C), 1 μg/ml Pam3CSK4, 5 μg/ml R848 (tlrl-r848-1, Invivogen), 100 ng/ml LPS (tlrl-eblps, Invivogen), or 1μM ODN2216 (tlrl-2216, Invivogen), or in combinations thereof, for the time periods specified in the experimental design.

**NY-ESO-$1_{157\text{-}165}$ restricted TCR T cell construction**

The TCR sequence that has been reported to specifically recognize the structure of HLA-A*02:01/NY-ESO-$1_{157\text{-}165}$ complex was utilized(*51*). The coding region of TCR was chemically synthesized and subsequently cloned into a lentiviral vector pRRLSIN-cPPT-SFFV-MCS-3FLAG-E2A-EGFP-SV40-puromycin (Genechem Co.,Ltd., Shanghai). Primary HLA-A2 negative $CD8^+$ T cells were pre-expanded with human CD3/CD28 T cell activation beads (422604, Biolegend) for 2 days, followed by transfection with lentivirus at a multiplicity of infection (MOI) of 50. Three days later, $EGFP^+$ T cells were isolated by FACS.

**Cell preparation and flow cytometry analysis**

Tissues were enzymatically digested with 1 mg/ml collagenase IV (C5138, Sigma-Aldrich) and 100 μg/ml DNase I (10104159001, Roche), treated with red cell lysis buffer (00-4333-57, ThermoFisher), and subjected to FACS analysis by Fortessa instrument (BD bioscience). For cell surface staining, cells were first blocked with FcR Blocking Reagent (130-059-901, Miltenyi Biotec) for 10 minutes at 4 °C, followed by staining with fluorochrome-conjugated monoclonal antibodies in Cell Staining Buffer (420201, BioLegend) for 1 hour at 4 °C. Regarding intracellular FACS analysis, the cells were first incubated with the Cell Activation Cocktail containing Brefeldin A (423304, Biolegend) at 37 °C for 6 hours. Subsequently, the BD

Cytofix/Cytoperm™ Fixation/Permeabilization Kit (554714, BD bioscience) was applied for sample processing. Finally, the cells were stained with relevant antibodies in BD Perm/Wash buffer (554723, BD bioscience) at 4 °C for 1 hour. The antibodies and reagents utilized in this study are listed in Supplementary Table 4. Data analysis was performed by FlowJo software 10.7.1.

**Tumor model establishment and treatment**

Three-week-old NCG-X mice were employed to establish the HuNCG-X mouse model. Specifically, $2\times10^5$ $CD34^+$ umbilical cord blood stem cells sourced from a healthy donor with the HLA-A*02:01 genotype were administered via the tail vein to each mouse. Mice with the proportion of human PBMCs exceeding 60% in the peripheral blood at week 14 were selected for subsequent experiments. Eighteen weeks after hematopoietic reconstitution, the tumor xenograft model was established by subcutaneously injecting the specified cell lines ($2\times10^6$ cells per mouse), which carried the HLA-A*02:01 genotype. Subsequently, on days 4, 7, and 10 following tumor implantation, pre-activated CBcDC1 cells with the HLA-A*02:01 genotype were intratumorally injected into tumor-bearing mice at a dose of $1\times10^6$ viable cells per mouse per administration. The immune responses were evaluated on day 16 after tumor inoculation, while the observation of tumor growth was extended until day 20 after tumor establishment.

**T cell depletion *in vivo***

$CD8^+$ T cells and $CD4^+$ T cells were depleted using the anti-CD8 antibody (clone OKT-8, Bio X Cell) and anti-CD4 antibody (clone OKT-4, Bio X Cell), respectively. These antibodies were diluted in PBS and administered via intraperitoneal injection. Injections were administered for 3 consecutive days immediately following tumor cell inoculation, and subsequently repeated every 3 days for a total duration of 20 days (100 μg per mouse per application). An IgG2a isotype antibody was employed as a control in this experiment.

**Generation of $EGFP^+$ SKOV3 cell line with or without HLA-A*02:01 genotype, and endocytosis experiments**

The coding region of HLA-A*02:01 was chemically synthesized and cloned into a lentiviral vector pRRLSIN-cPPT-CMV-MCS-3FLAG-E2A-EGFP-SV40-puromycin (Genechem Co.,Ltd., Shanghai). Subsequently, SKOV3 cells were transfected either the empty lentivirus or the lentivirus overexpressing HLA-A*02:01 at a MOI of 1:50. The $EGFP^+$ SKOV3 cells were sorted using FACS.

For endocytosis studies, $1\times10^5$ CBcDC1 cells were co-cultured with $2\times10^4$ $EGFP^+$ SKOV3 cells. The co-culture was carried out in the presence of 1 μg/ml Poly(I:C) and 1 μg/ml Pam3CSK4 for 0, 6 and 12 hours. Subsequently, the endocytosis of the GFP antigen was detected by flow cytometry.

**CRISPR/CAS9 mediated gene editing**

Isolated $CD34^+$ CB HSCs were first expanded in the StemSpan™-AOF medium with 100 ng/ml SCF, 100 ng/ml TPO, 100 ng/ml FLT3L, and 1 μM SR1 for 2 days. Subsequently, the sgRNA-Cas9 protein complex (GenCrispt) were introduced into HSCs via electroporation with the Neon Transfection Kit (Thermofisher) according to the protocol manual. After that, the cells continued to expand for another two days and then started to differentiate into cDCs. The sgRNA sequences designed to target each genes are presented in Supplementary Table 5.

**TFs overexpression**

The coding regions of human *IRF8* (NM_002163), *BATF3* (NM_018664) or *ZNF366* (NM_152625) were synthesized and cloned into the multiple cloning sites of the EF1α-MCS-Flag-PGK-EGFP lentiviral plasmid (Tsingke Biotech, China). Viral particles for overexpression were transduced at a MOI of 150 on day 3 of CB-HSPC pre-expansion, followed by continued induction of stem cell differentiation into cDCs.

**DC redifferentiation and proliferation assay**

CBcDC subpopulations were isolated on day 8 after cDC induction. For the redifferentiation assay, the isolated subpopulations were cultured in StemSpan™-AOF medium supplemented with 100 ng/ml SCF, 100 ng/ml FLT3L, 1 μM SR1, 5 ng/ml IFN-γ, 2.5 ng/ml GM-CSF, and 2.5 ng/ml IL-4 for another 2 or 4 days. For the proliferation assay, the subpopulations were stained with CellTrace™ Violet (CTV, Thermofisher) and then cultured in the same supplemented StemSpan™-AOF medium for 2 days.

**ScRNA-seq**

Cell suspensions were loaded meticulously onto Chromium microfluidic chips with the 3' v3.1 chemistry, followed by single-cell barcoding on a 10×Chromium Controller (10×Genomics). Barcoded cells were subjected to reverse transcription, and sequencing libraries were prepared with reagents from the Chromium Single Cell 3' v3.1 kit (10×Genomics), strictly adhering to the manufacturer's protocols. Paired-end sequencing was performed on an Illumina NovaSeq 6000 platform in accordance with standard operating procedures provided by Illumina.

Raw sequencing reads were quality-controlled and summarized using Fastp. For downstream analyses, raw base call (BCL) files generated by the Illumina sequencer were converted to FASTQ format, which served as input for Cell Ranger Count (10× Genomics). Reads were demultiplexed and aligned to the reference genome using the Cell Ranger pipeline (https://support.10xgenomics.com/single-cell-gene-expression/ software/pipelines/latest/what-is-cell-ranger) with default parameters. Unless specified otherwise, all subsequent single-cell transcriptomic analyses were executed using Cell Ranger and Seurat (version 5.1.0). Briefly, unique molecular identifiers (UMIs) were quantified for each gene and cell barcode (post Cell Ranger filtering) to generate digital gene expression matrices. A secondary filtration step was applied in Seurat to retain high-quality cells and genes: only genes detected in ≥3 cells, and cells expressing ≥200 genes, were included in downstream analyses.

Differential gene expression analysis between experimental groups was performed on the Seurat-filtered expression matrices using the edgeR package, enabling the identification of region-specific marker genes. Cellular differentiation trajectories were reconstructed with Monocle3 (version 3.34.0), which infers developmental trajectories from the initial to terminal cellular states.

Gene Ontology (GO) and Kyoto Encyclopedia of Genes and Genomes (KEGG) enrichment analyses of marker genes were conducted using the clusterProfiler R package (version 4.14.6), with correction for gene-length bias. Terms with a nominal P-value < 0.05 were considered statistically significant.

T cell receptor (TCR) sequencing data from treatment and control groups were processed and annotated using Cell Ranger V(D)J, and imported into scRepertoire (version 2.2.1) for clonotype analysis. TCR annotations were merged using the combineTCR function with parameters removeNA = TRUE, removeMulti = TRUE, and filterMulti = TRUE—a step that excluded entries with missing annotations and multi-mapped TCR chains. TCR clonotype metadata (including clone size) were then integrated into Seurat objects via the combineExpression function for joint transcriptomic and TCR repertoire analysis.

**Smart-RNA sequencing**

For each sample, $1\times10^4$ cells were utilized to generate full-length cDNA using the Single Cell Full Length mRNA-Amplification Kit (Vazyme, Nanjing, China). Subsequently, next-generation libraries were prepared by employing the TruePrep DNA Library Prep Kit V2 for Illumina (Vazyme, Nanjing, China). The quality of the library was evaluated using the Bioanalyzer 2100 (Agilent, Santa Clara, CA, USA). The RNA libraries were sequenced on the Novaseq 6000 system (Illumina, San Diego, CA, USA). Genes with a $p$ value less than 0.05 as determined by DEseq2 analysis and a $\log_2$ fold-change greater than 1 were selected for further investigation. GO analysis was carried out to identify the enriched pathways of the selected genes.

**Cytokine detection**

The levels of TNF-α, IL-6, IL-12p70, IP-10, and IFN-γ were respectively determined using the BD™ Cytometric Bead Array (CBA) Human TNF, IL-6, IL-12p70, IP10 and IFN-γ Flex Sets (BD Bioscience). The levels of IFN-λ were detected by the Human IL-29 ELISA Kit (absin, China).

**Antigen presentation assay of CBcDC**

Control or NY-ESO-$1_{157\text{-}165}$ TCR-T cells were initially labeled with CellTrace™ Violet (ThermoFisher). Subsequently, $5\times10^4$ CBcDCs were incubated for 12 hours with 1 µg/ml NY-ESO-1 peptides (Miltenyi Biotech) or 10 µg/ml CTAG1B protein (Genwiz) with no stimulation, or 100 µg/ml cell lysate prepared from HLA-A*02-negative SKOV3 cells with overexpressed CTAG1B in the presence of 1 µg/mL Poly(I:C) and 1 µg/mL Pam3CSK4 stimulation. After that, the DCs were washed and co-incubated with Ctrl-T or TCR-T cells at a DC-to-T cell ratio of 1:2 for 48 hours. In the context of cell-associated antigen cross-presentation, $2\times10^4$ live or X-ray-irradiated HLA-A*02-negative SKOV3 cells overexpressing CTAG1B were cocultured with $5\times10^4$ CBcDCs for 12 hours in the presence or absence of 1 µg/mL Poly(I:C) and 1 µg/mL Pam3CSK4 stimulation. Subsequently, CBcDCs were isolated, washed and co-incubated with $1\times10^5$ CTV-labeled Ctrl-T or TCR-T cells at a DC-to-T cell ratio of 1:2 for 48 hours, and the proliferation of T cells was detected by flow cytometry. For the titration assay, $5\times10^4$ CBcDC1s were co-incubated with viable HLA-A*02-negative SKOV3-CTAG1B cells for 12 hours in the presence of 1 µg/mL poly(I:C) and 1 µg/mL Pam3CSK4 stimulation, or alternatively co-incubated with 1 µg/mL NY-ESO-1 peptides for 12 hours. The CBcDC1s were then washed and subsequently co-cultured with CTV-labeled Ctrl-T or TCR-T cells at the indicated ratios for 48 hours.

**Statistics**

Statistical analyses were conducted using GraphPad Prism software (version 10.1.2; GraphPad Software Inc.). Normality of data was assessed using the Shapiro-Wilk test. For comparisons between two independent groups: Normally distributed data with equal variances were analyzed using unpaired two-tailed t-tests; non-normally distributed data were analyzed using the Mann-Whitney U test. For comparisons among three or more independent groups: Normally distributed data with equal variances were analyzed using one-way analysis of variance (one-way ANOVA) with Dunnett's multiple comparisons test. Two-way ANOVA with Bonferroni post hoc test was applied for multi-group comparisons involving two independent variables, to account for multiple comparisons. Data are presented as mean ±SD and N represents the total number of animals or biological replicates. Survival curves were analyzed via the log-rank (Mantel-Cox) test, and dynamic tumor growth changes were assessed using two-way ANOVA with multiple comparisons. For single-cell RNA sequencing (scRNA-seq) data, differences in the proportion of each T cell subcluster between the treatment and control groups were compared using Pearson's chi-squared ($\chi^2$) test, implemented with the prop.test function in R. Within each T cell subcluster, differential gene expression analysis between the treatment and control groups was performed on scRNA-seq data using the FindMarkers function in Seurat, with the Wilcoxon rank-sum test specified (test.use = "wilcox"). Statistical significance was defined as $*p < 0.05$, $**p < 0.01$, $***p < 0.001$, and $****p < 0.0001$. The specific statistical tests used for each experiment are detailed in the corresponding figure legends.

**Data availability**

ScRNA-seq data corresponding to CBcDC differentiation and tumor-infiltrating immune cells were deposited in the Genome Sequence Archive for Human (GSA-Human; https://ngdc.cncb.ac.cn/gsa/) under the accession numbers HRA011482 and HRA011459. RNA sequencing (RNA-seq) data of distinct DC subsets were deposited in the same repository with the accession number HRA011154.

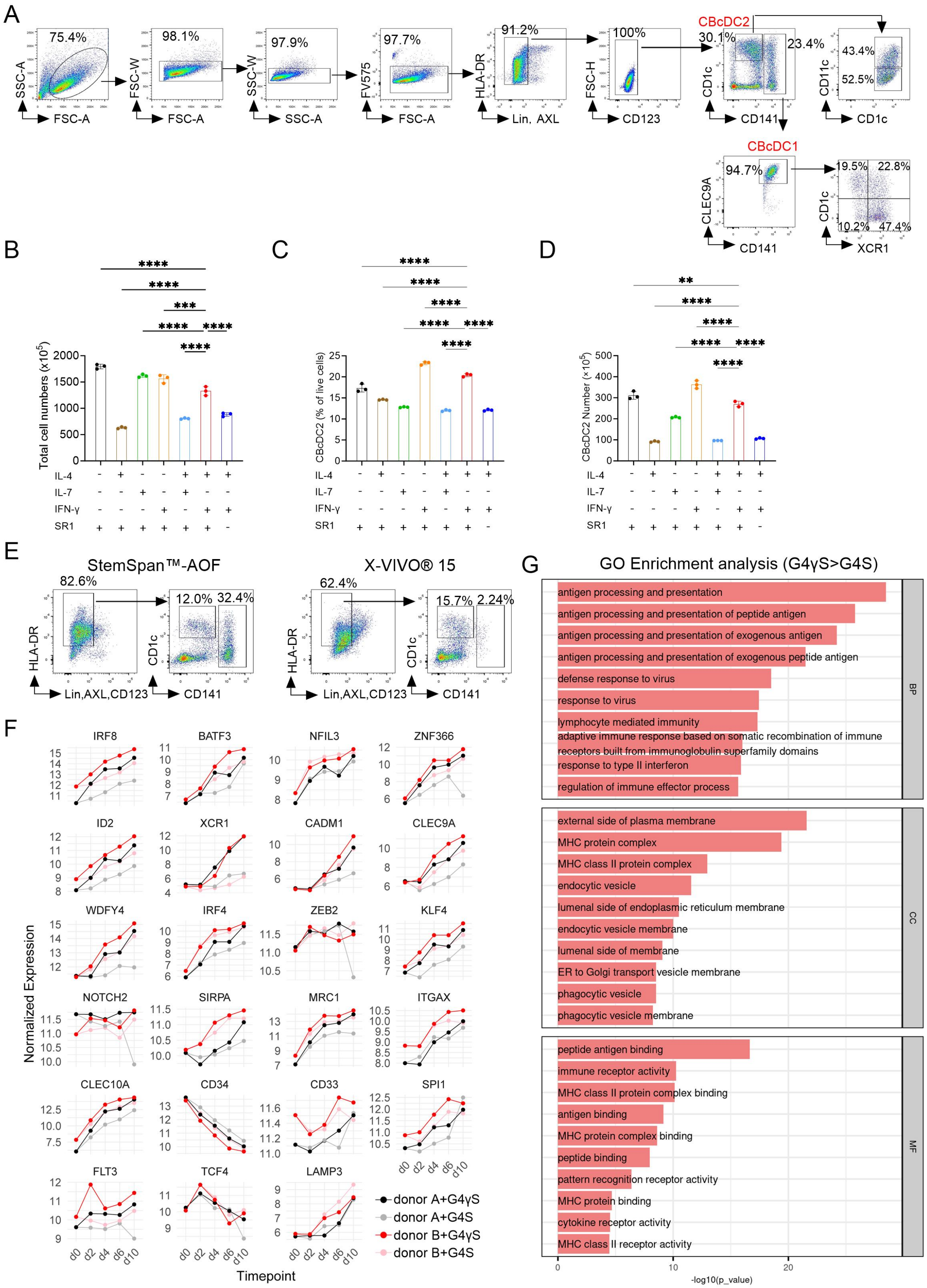

A
75.4%
SSC-A
FSC-A
98.1%
FSC-W
FSC-A
97.9%
SSC-W
SSC-A
97.7%
FV575
FSC-A
91.2%
HLA-DR
Lin, AXL
100%
FSC-H
CD123
CBcDC2
30.1%
23.4%
CD1c
CD141
43.4%
52.5%
CD11c
CD1c
CBcDC1
94.7%
CLEC9A
CD141
19.5%
22.8%
10.2%
47.4%
CD1c
XCR1
B
Total cell numbers (x10^5)
IL-4
IL-7
IFN-γ
SR1
C
CBcDC2 (% of live cells)
D
CBcDC2 Number (x10^5)
E
StemSpan™-AOF
82.6%
12.0%
32.4%
HLA-DR
Lin,AXL,CD123
CD1c
CD141
X-VIVO® 15
62.4%
15.7%
2.24%
F
IRF8
BATF3
NFIL3
ZNF366
ID2
XCR1
CADM1
CLEC9A
WDFY4
IRF4
ZEB2
KLF4
NOTCH2
SIRPA
MRC1
ITGAX
CLEC10A
CD34
CD33
SPI1
FLT3
TCF4
LAMP3
Normalized Expression
Timepoint
d0 d2 d4 d6 d10
donor A+G4γS
donor A+G4S
donor B+G4γS
donor B+G4S
G
GO Enrichment analysis (G4γS>G4S)
antigen processing and presentation
antigen processing and presentation of peptide antigen
antigen processing and presentation of exogenous antigen
antigen processing and presentation of exogenous peptide antigen
defense response to virus
response to virus
lymphocyte mediated immunity
adaptive immune response based on somatic recombination of immune receptors built from immunoglobulin superfamily domains
response to type II interferon
regulation of immune effector process
BP
external side of plasma membrane
MHC protein complex
MHC class II protein complex
endocytic vesicle
lumenal side of endoplasmic reticulum membrane
endocytic vesicle membrane
lumenal side of membrane
ER to Golgi transport vesicle membrane
phagocytic vesicle
phagocytic vesicle membrane
CC
peptide antigen binding
immune receptor activity
MHC class II protein complex binding
antigen binding
MHC protein complex binding
peptide binding
pattern recognition receptor activity
MHC protein binding
cytokine receptor activity
MHC class II receptor activity
MF
-log10(p_value)

**Fig. S1. Optimization of Culture Conditions for Inducing CBcDCs.** (**A**) Representative flow cytometry analysis for identification of CBcDC subclusters. (**B**) Expansion of total cells from $1 \times 10^5$ CB HSPC cultured with various combinations of IL-4, IL-7, IFN-γ, and SR1. (**C**, **D**) Effect of different combinations (IL-4, IL-7, IFN-γ, and SR1) on the induction of CBcDC2. Induction efficiency (**C**) and yield (**D**) of CBcDC2 generated from an input of $1 \times 10^5$ CB HSPC. N=3. (**B**-**D**) data are presented as the mean ± SD and were analyzed by one-way ANOVA with Dunnett's multiple comparisons test. **p<0.01, ***p<0.001, ****p<0.0001. (**E**) Impact of different culture media on CBcDC induction. (**F**) Dynamic change of marker genes expression during CBcDC induction. (**G**) GO enrichment analysis of pathways upregulated in CBcDCs induced with IFN-γ supplementation compared to those without.

<u>Related to Fig. 1.</u>

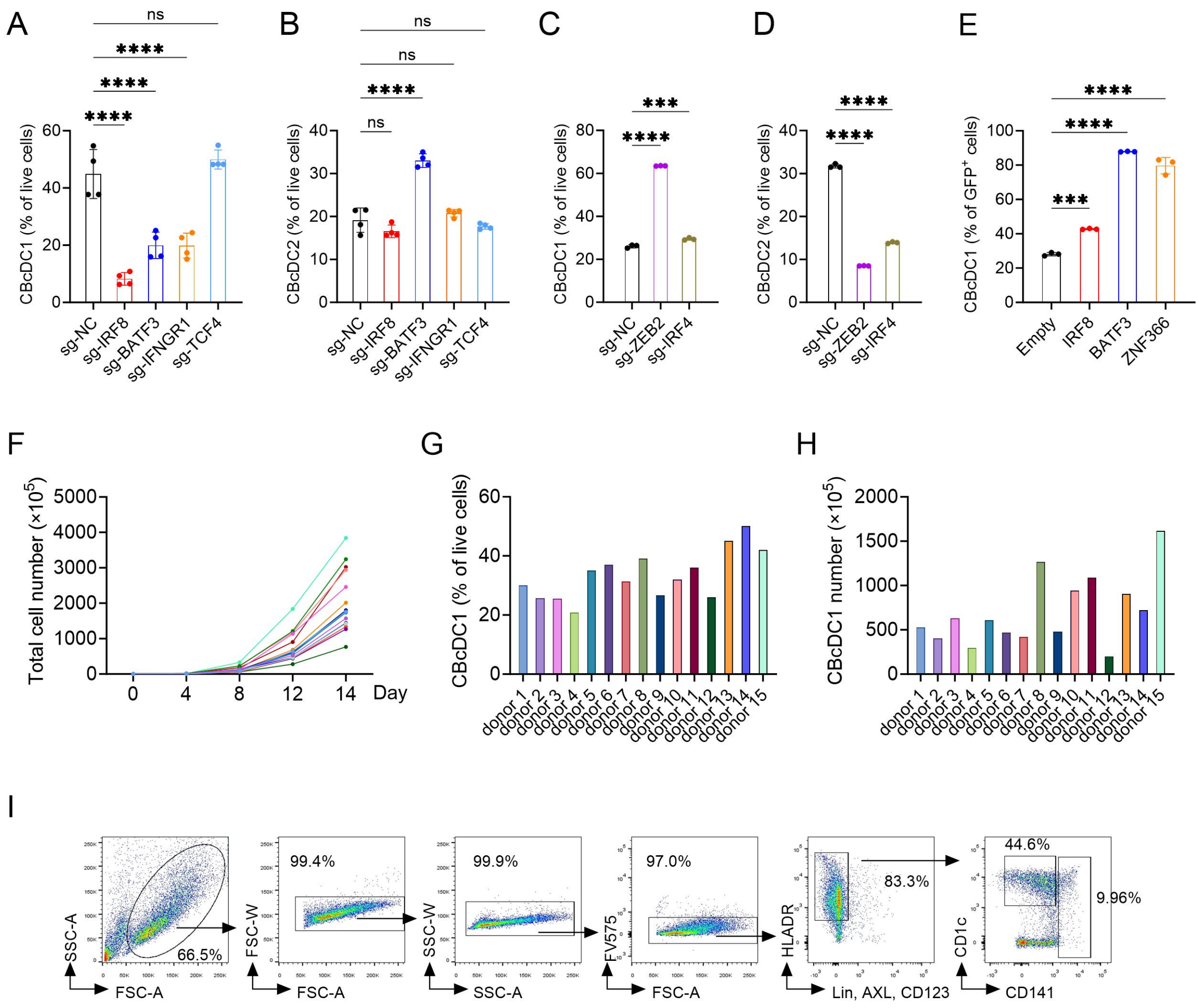


**Fig. S2. The Generalizability of This Induction System for cDC1 Expansion.** (**A**, **B**) Impact of *IRF8*, *BATF3*, *IFNGR1* or *TCF4* knockout on CBcDC1 (**A**) and CBcDC2 (**B**) differentiation. (**C**, **D**) Effect of *ZEB2* or *IRF4* knockout on CBcDC1 (**C**) and CBcDC2 (**D**) differentiation. (**E**) Influence of *IRF8*, *BATF3* or *ZNF366* overexpression on CBcDC1 differentiation. N=3 or 4. (**A**-**E**) data are presented as the mean ± SD and were analyzed by one-way ANOVA with Dunnett's multiple comparisons test. ***p<0.001, ****p<0.0001, ns, no significant. (**F**) Expansion kinetics of total cell numbers during CBcDC1 induction, starting from $1\times10^5$ $CD34^+$ HSPCs (15 different donors). (**G**) The induction efficiency of CBcDC1. (**H**) Quantification of CBcDC1 yield from $1\times10^5$ $CD34^+$ HSPCs. (**I**) Generation of cDC from $CD34^+$ HSPCs isolated from adult peripheral blood using the G4γS culture system.

Related to Fig. 1.

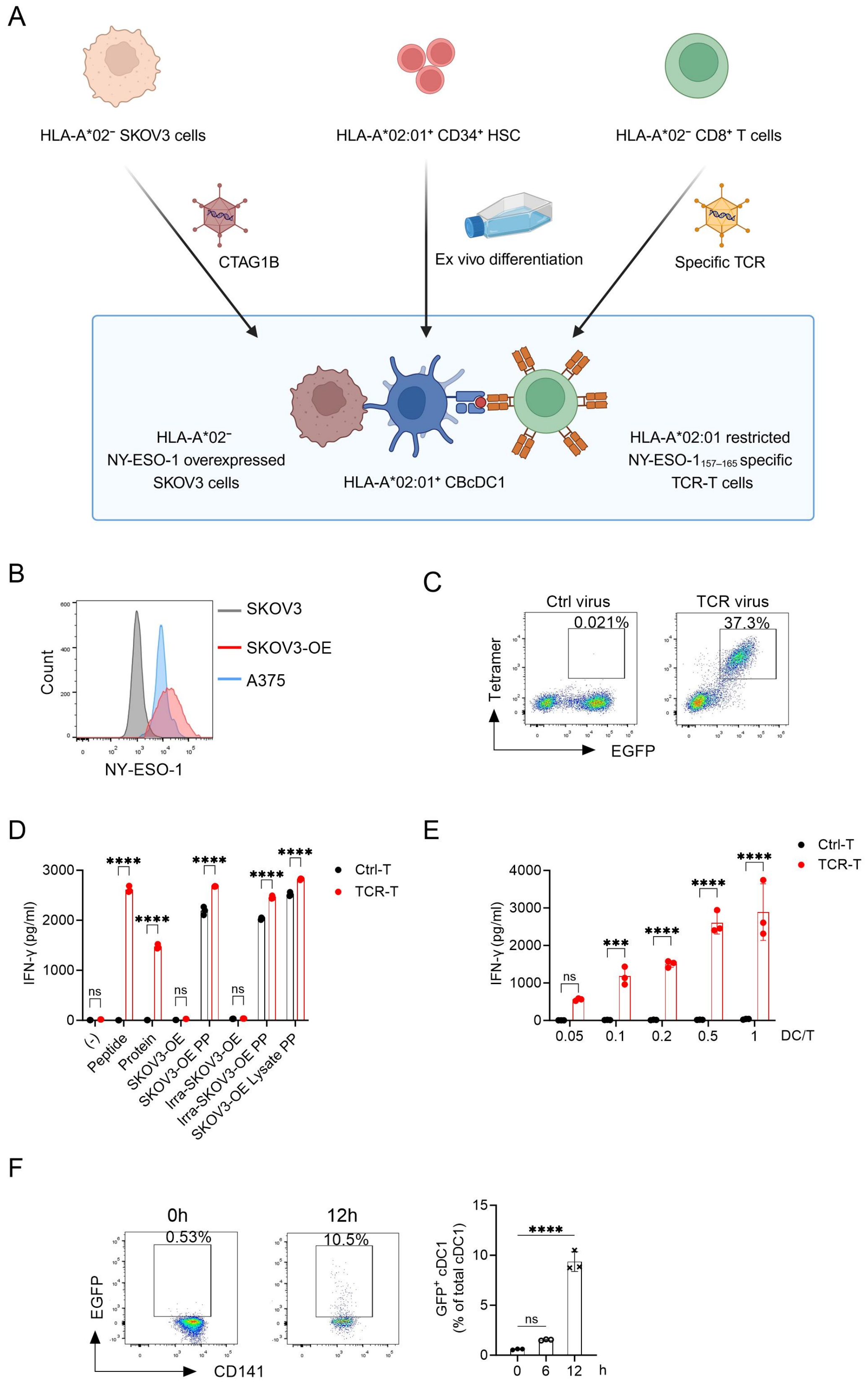
A
HLA-A*02⁻ SKOV3 cells
HLA-A*02:01⁺ CD34⁺ HSC
HLA-A*02⁻ CD8⁺ T cells
CTAG1B
Ex vivo differentiation
Specific TCR
HLA-A*02⁻
NY-ESO-1 overexpressed
SKOV3 cells
HLA-A*02:01⁺ CBcDC1
HLA-A*02:01 restricted
NY-ESO-1157–165 specific
TCR-T cells
B
Count
NY-ESO-1
SKOV3
SKOV3-OE
A375
C
Ctrl virus
TCR virus
0.021%
37.3%
Tetramer
EGFP
D
IFN-γ (pg/ml)
Ctrl-T
TCR-T
ns
****
(-)
Peptide
Protein
SKOV3-OE
SKOV3-OE PP
Irra-SKOV3-OE
Irra-SKOV3-OE PP
SKOV3-OE Lysate PP
E
IFN-γ (pg/ml)
Ctrl-T
TCR-T
ns
***
****
0.05
0.1
0.2
0.5
1
DC/T
F
0h
12h
0.53%
10.5%
EGFP
CD141
GFP⁺ cDC1
(% of total cDC1)
ns
****
0
6
12
h

**Fig. S3. CBcDC1s Exhibit Potent Antigen Cross-Presentation Capacity.** (**A**) Schematic diagram illustrating the setup for the cross-presentation experiment. (**B**) The expression of NY-ESO-1 in SKOV3 cell line with CTAG1B overexpression. A375 cell line was used as positive control. (**C**) The expression of TCR specifically recognized the HLA-A*02:01/NY-ESO-$1_{157\text{-}165}$ complex. (**D**) IFN-γ secretion by TCR-T cells upon cross-activation by CBcDC1s. (**E**) IFN-γ secretion by TCR-T cells cross-activated by NY-ESO-1 peptide-pulsed CBcDC1s at varying DC:T ratios. (**F**) Phagocytosis of CBcDC1. CBcDC1 was incubated with $EGFP^+$ SKOV3 cells for 0, 6 and 12 hours to evaluate cellular antigen uptake. N=3. (**D**-**F**) data are presented as the mean ± SD. (**D**) and (**E**) data were analyzed by two-way ANOVA with Bonferroni post-hoc test; (**F**) by one-way ANOVA with Dunnett's multiple comparisons test. ***$p<0.001$, ****$p<0.0001$, ns, no significant.

<u>Related to Fig. 2.</u>

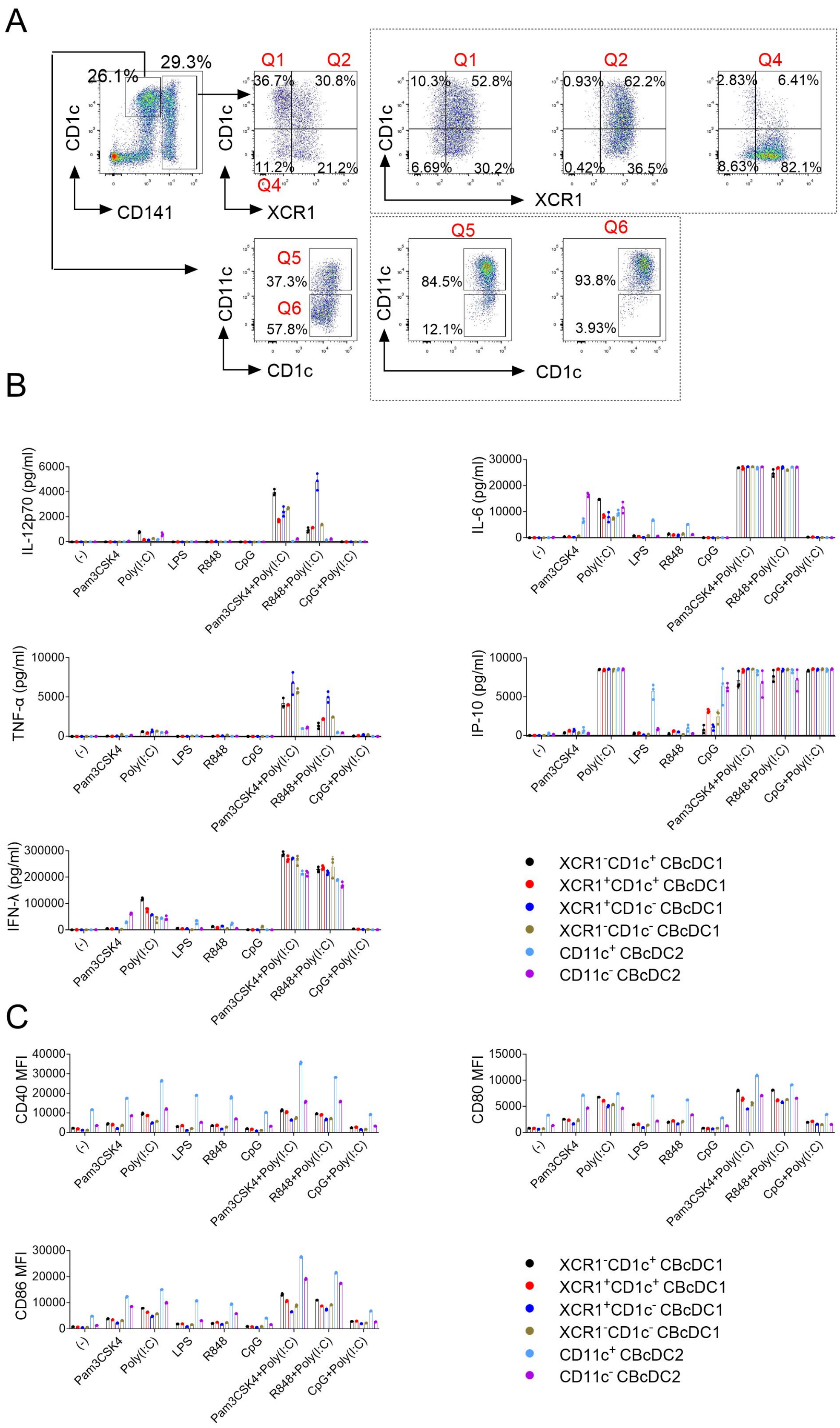
A
B
C
CD1c
CD141
26.1%
29.3%
Q1
Q2
Q4
36.7%
30.8%
11.2%
21.2%
XCR1
10.3%
52.8%
6.69%
30.2%
0.93%
62.2%
0.42%
36.5%
2.83%
6.41%
8.63%
82.1%
CD11c
Q5
Q6
37.3%
57.8%
84.5%
12.1%
93.8%
3.93%
IL-12p70 (pg/ml)
IL-6 (pg/ml)
TNF-α (pg/ml)
IP-10 (pg/ml)
IFN-λ (pg/ml)
CD40 MFI
CD80 MFI
CD86 MFI
(-)
Pam3CSK4
Poly(I:C)
LPS
R848
CpG
Pam3CSK4+Poly(I:C)
R848+Poly(I:C)
CpG+Poly(I:C)
XCR1⁻CD1c⁺ CBcDC1
XCR1⁺CD1c⁺ CBcDC1
XCR1⁺CD1c⁻ CBcDC1
XCR1⁻CD1c⁻ CBcDC1
CD11c⁺ CBcDC2
CD11c⁻ CBcDC2

**Fig. S4. Functional Heterogeneity of CBcDC Subclusters.** (**A**) At day 8 of DC-directed differentiation, four CBcDC1 subclusters were defined by XCR1 and CD1c expression, and two CBcDC2 subclusters by CD11c expression. Sorted CBcDC subclusters were cultured in differentiation medium for an additional 4 days to assess their differentiation capacity. Plots outside the box show initial differentiation states, whereas those inside show further differentiation outcomes following extended culture. (**B**, **C**) Functional characteristics of CBcDC subclusters. The expression of cytokines/chemokine (**B**) and co-stimulatory molecules (**C**) in CBcDC subclusters following treatment with the indicated stimulus. N=3.
Related to Fig. 2.

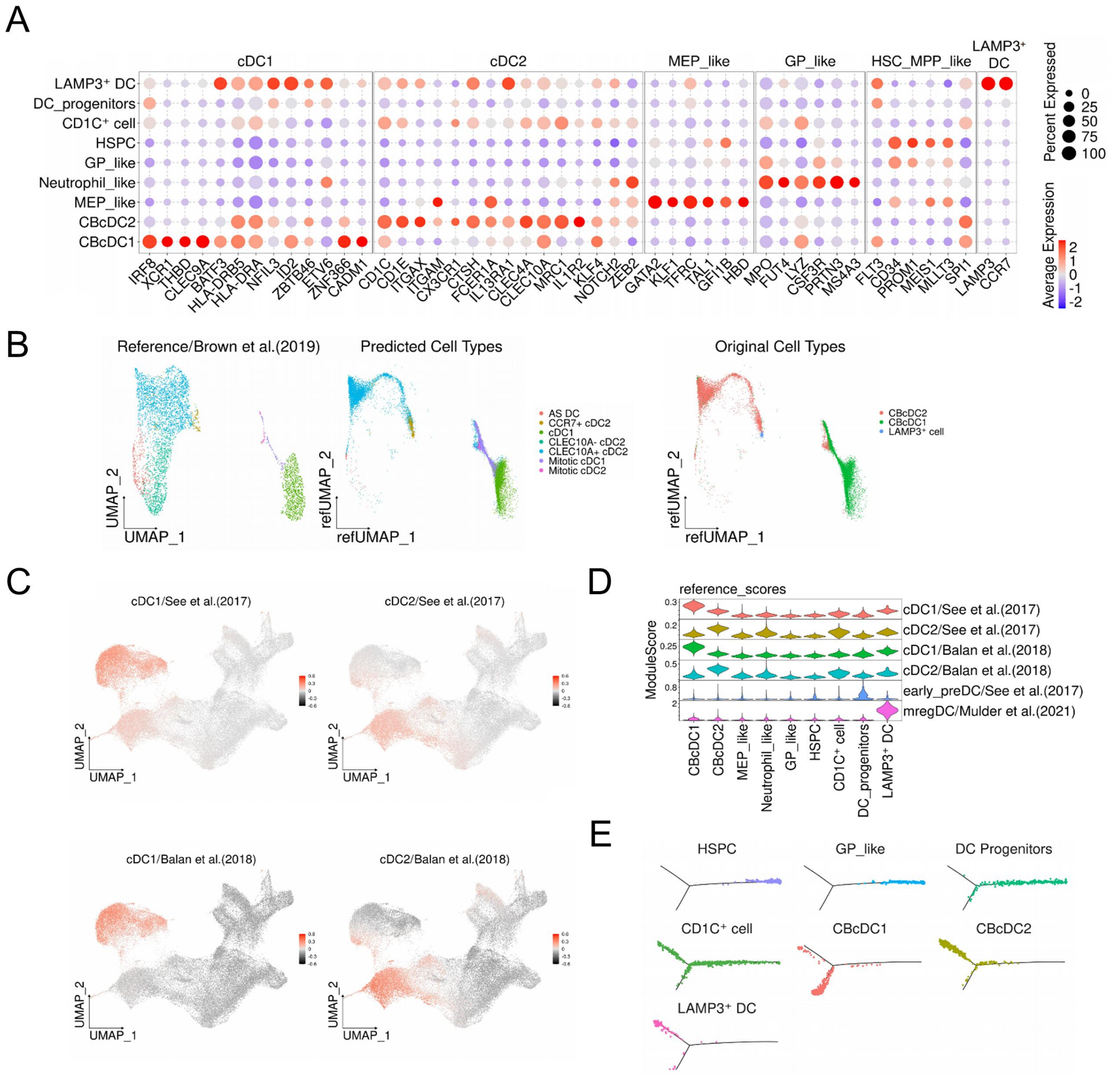


**Fig. S5. Induced DC Subsets Exhibit High Similarity to Corresponding Endogenous DC Subsets.** (**A**) Dot plot showing the relative expression levels of selected genes across different clusters. (**B**) UMAP plots depicting the projection of CBcDC1, CBcDC2, and LAMP3$^{+}$ DC cells from this study onto the human splenocyte atlas generated by Brown et al. The left plot displays the original atlas of DCs from Brown et al. The middle plot shows the DCs from this study mapped onto the atlas of Brown et al. with their cellular annotation. The plot on the right illustrates the DCs from this study mapped onto the atlas of Brown et al., but with our cellular annotation. (**C**) UMAP plots showing the enrichment scores of human cDC1 and cDC2 reference gene sets. Each dot represents a single cell, and the color of the dot indicates the enrichment score. (**D**) Violin plots showing the enrichment scores of human cDC1, cDC2, early pre-DC and mregDC reference gene sets across the nine cell populations of CB-derived cells. (**E**) Individual trajectory plots displaying the distribution of each cell population from Fig. 3E along the pseudo time axis.

<u>**Related to Fig. 3.**</u>

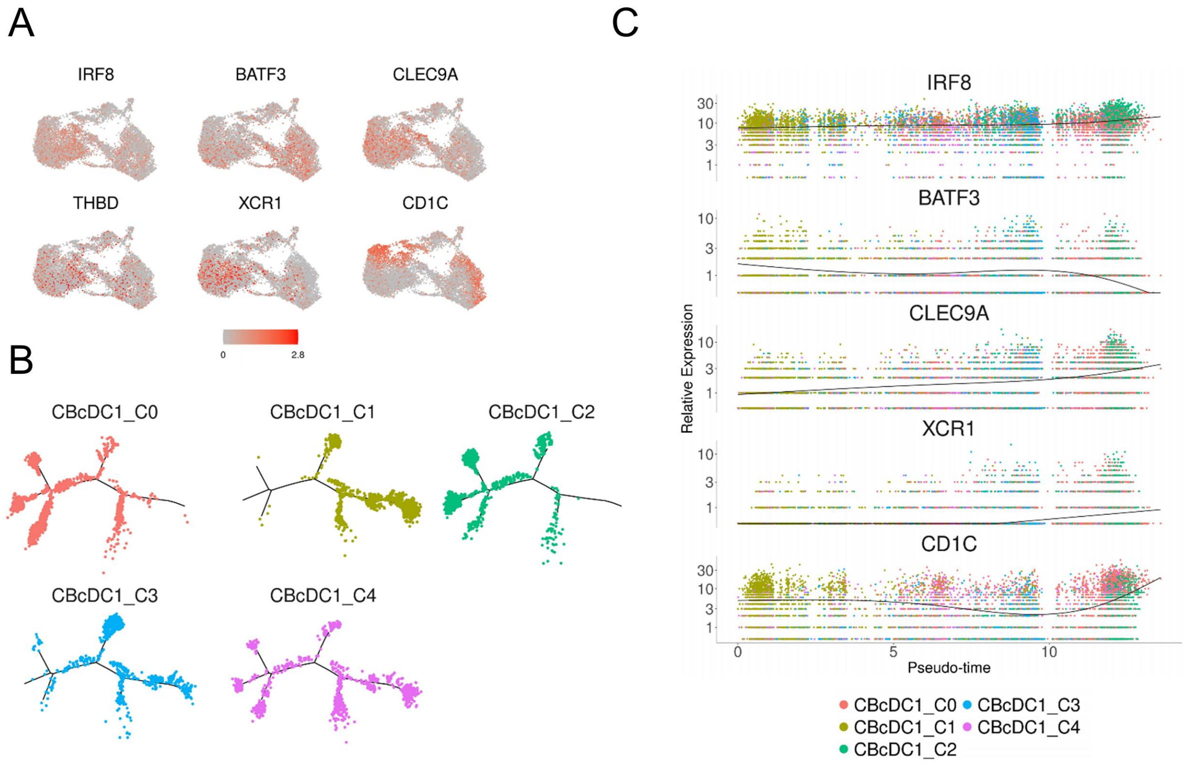


**Fig. S6. Characteristics of the CBcDC1 Subclusters.** (**A**) UMAP plots illustrating the expression patterns of selected genes within CBcDC1 subclusters, with the color scale indicating relative expression levels. (**B**) Individual trajectory plots displaying the distribution of each CBcDC1 subcluster along the pseudo time axis. (**C**) Relative expression of key genes across CBcDC1 subclusters. Each dot represents an individual cell, colored by subcluster identity. The overlaid smooth curve depicts the overall temporal trend in gene expression within the total CBcDC1 population.

<u>Related to Fig. 4.</u>

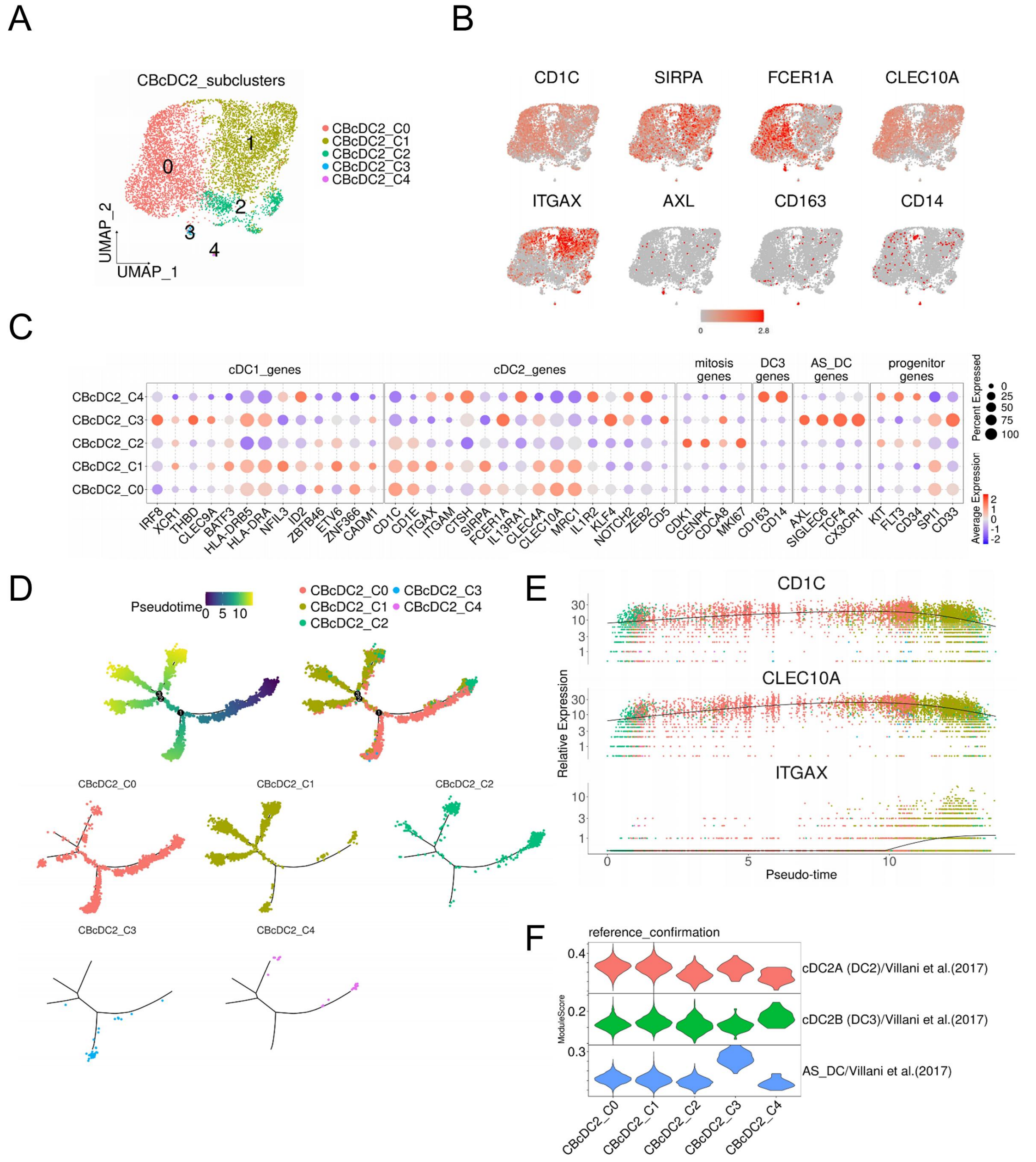


**Fig. S7. Unbiased Analysis of the Composition of the CBcDC2 Subclusters.** (**A**) UMAP plot showing unsupervised subclusters within the CBcDC2. (**B**) The UMAP plots showing the expression of selected genes in each cell. (**C**) Dot plot depicting the relative expression levels of selected genes across different CBcDC2 subclusters. (**D**) Pseudo time trajectory plot showing the progression of cells along the inferred developmental timeline. The upper two plots show the pseudo time trajectory of CBcDC2, with different colors representing pseudo time (top left) and different CBcDC2 subclusters (top right). The five lower plots show distribution of each subcluster along the pseudo time trajectory separately. (**E**) Relative expression of key genes across CBcDC2 subclusters. Each dot represents an individual cell, colored by subcluster identity. The overlaid smooth curve depicts the overall temporal trend in gene expression within the total CBcDC2 population. (**F**) Violin plots showing the enrichment scores of human cDC2A, cDC2B,

and AS_DC reference gene sets across each CBcDC2 subcluster.
Related to Fig. 4.

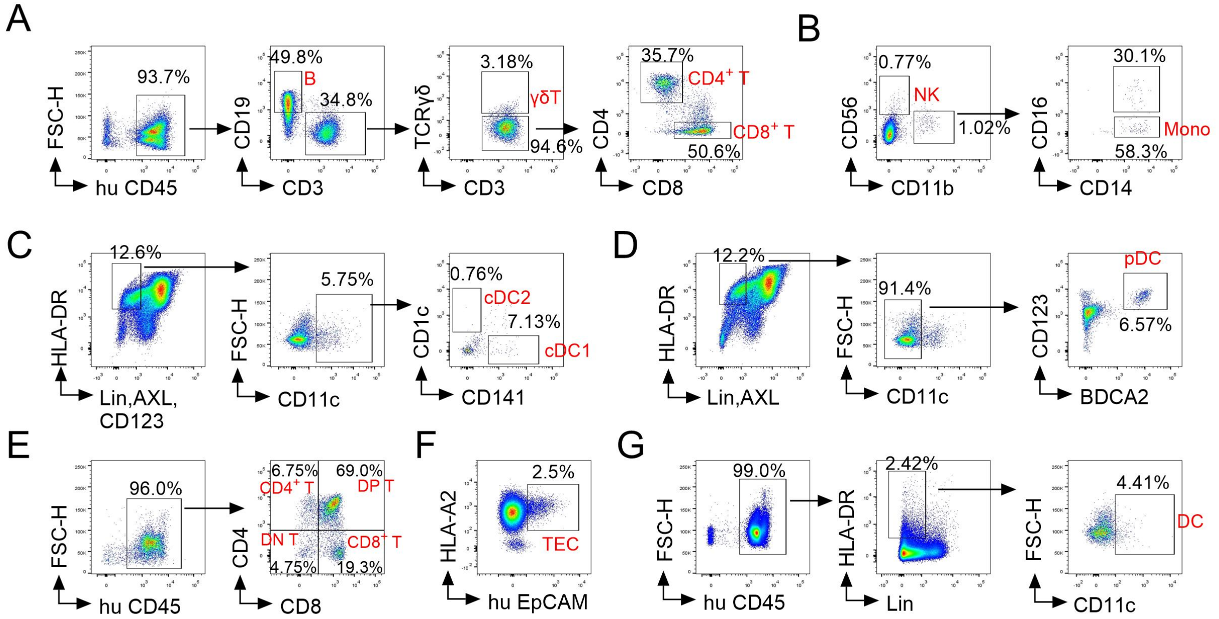


**Fig. S8. Validation of the Reconstitution of huNCG-X mice.** CD34$^+$ HSPCs obtained from a healthy donor with the HLA-A*02:01 genotype were infused into NCG-X mice. Analysis of these mice was conducted 18 weeks following the injection. (**A**) The proportion of human B cells and T cells in the spleen. (**B**) The proportion of NK cells and monocytes (gated on human CD45$^+$) in the spleen. (**C**) The proportion of cDC1 and cDC2 (gated on human CD45$^+$) in the spleen. (**D**) The proportion of pDC (gated on human CD45$^+$) in the spleen. (**E**) The proportion of T cells in the thymus. (**F**) The proportion of human epithelial cells in the thymus (TECs). (**G**) The proportion of human DCs in the thymus.

Related to Fig. 5.

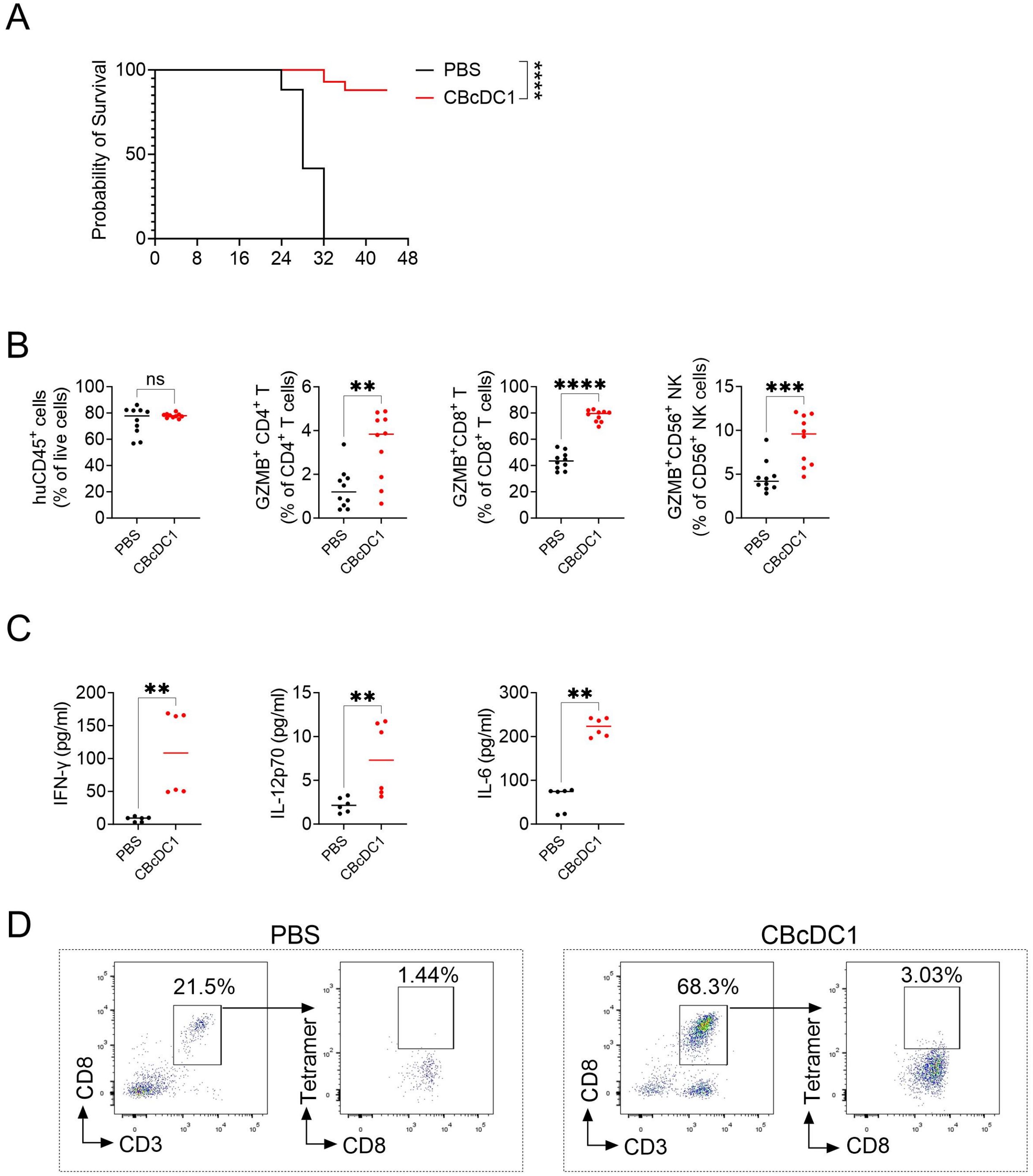


**Fig. S9. Therapeutic Efficacy of CBcDC1s in Melanoma Models.** (**A**) Kaplan–Meier survival curves of experimental mice over time. PBS, N=17; CBcDC1, N=9. Survival analysis was performed by log-rank Mantel-Cox test. (**B**) Frequency of the indicated cell types in the spleen of huNCG-X mice bearing A375 tumors. N=10. (**C**) Quantitative analysis of serum levels of IFN-γ, IL-12p70, and IL-6 in huNCG-X mice. N=6. (**B**) and (**C**) data were analyzed by Mann-Whitney U test. **$p<0.01$, ***$p<0.001$, ****$p<0.0001$, ns, no significant. (**D**) The expansion of HLA-A*02:01 restricted NY-ESO-$1_{157\text{-}165}$ targeted TCR T cells *in vivo*. Tumor single cells were first gated as human CD45$^+$ cells, and then gated as CD3$^+$CD8$^+$HLA-A*02:01/NY-ESO-1 tetramer$^+$

cells.
Related to Fig. 5.

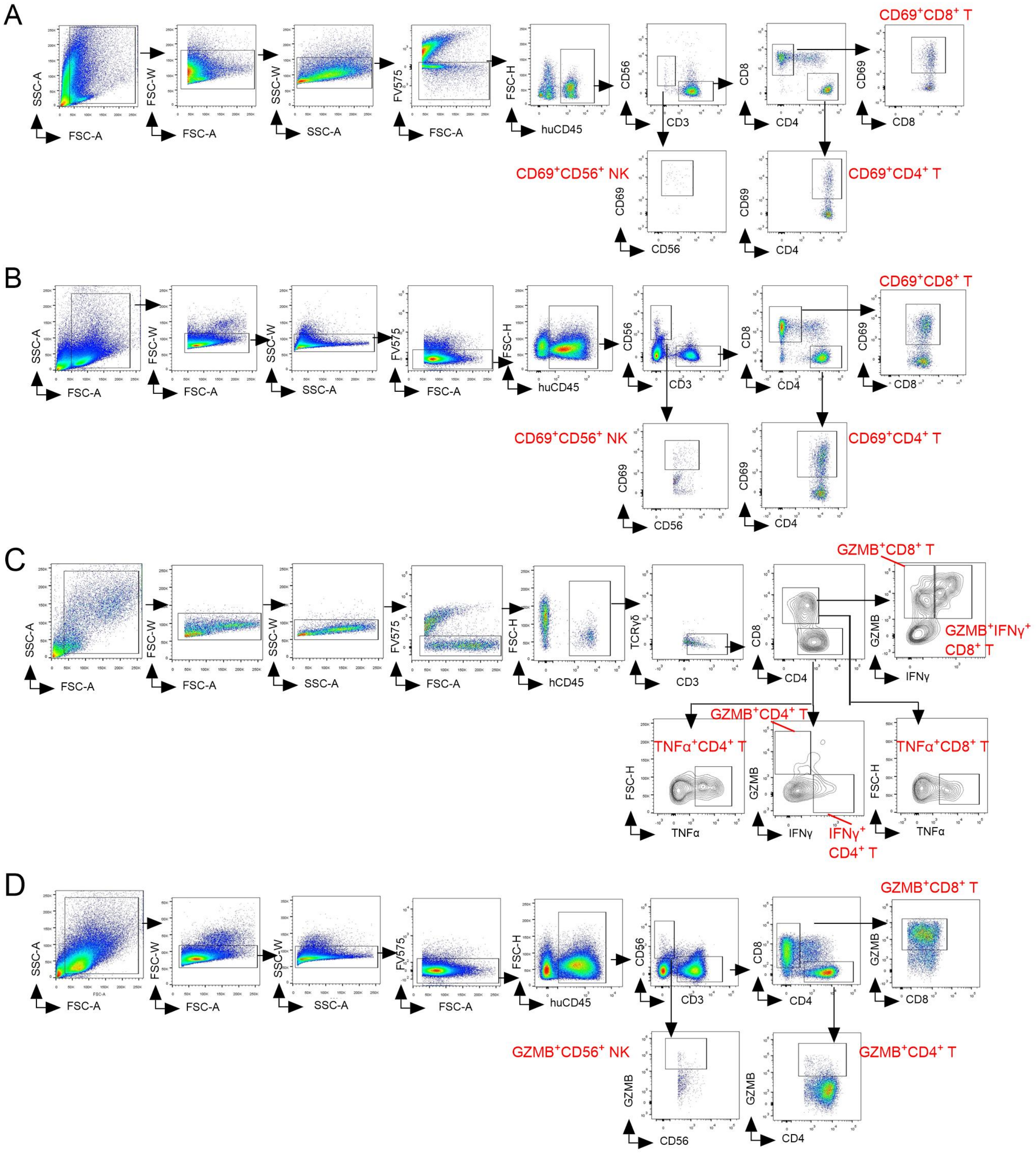


**Fig. S10. Gating Strategies for Activated and Cytotoxic T/NK Cells in Tumor and Spleen.** Gating strategies for CD69-positive T and NK cells in tumor (**A**) and spleen (**B**), cytotoxic T and NK cells in tumor (**C**) and spleen (**D**).

Related to Fig. 5.

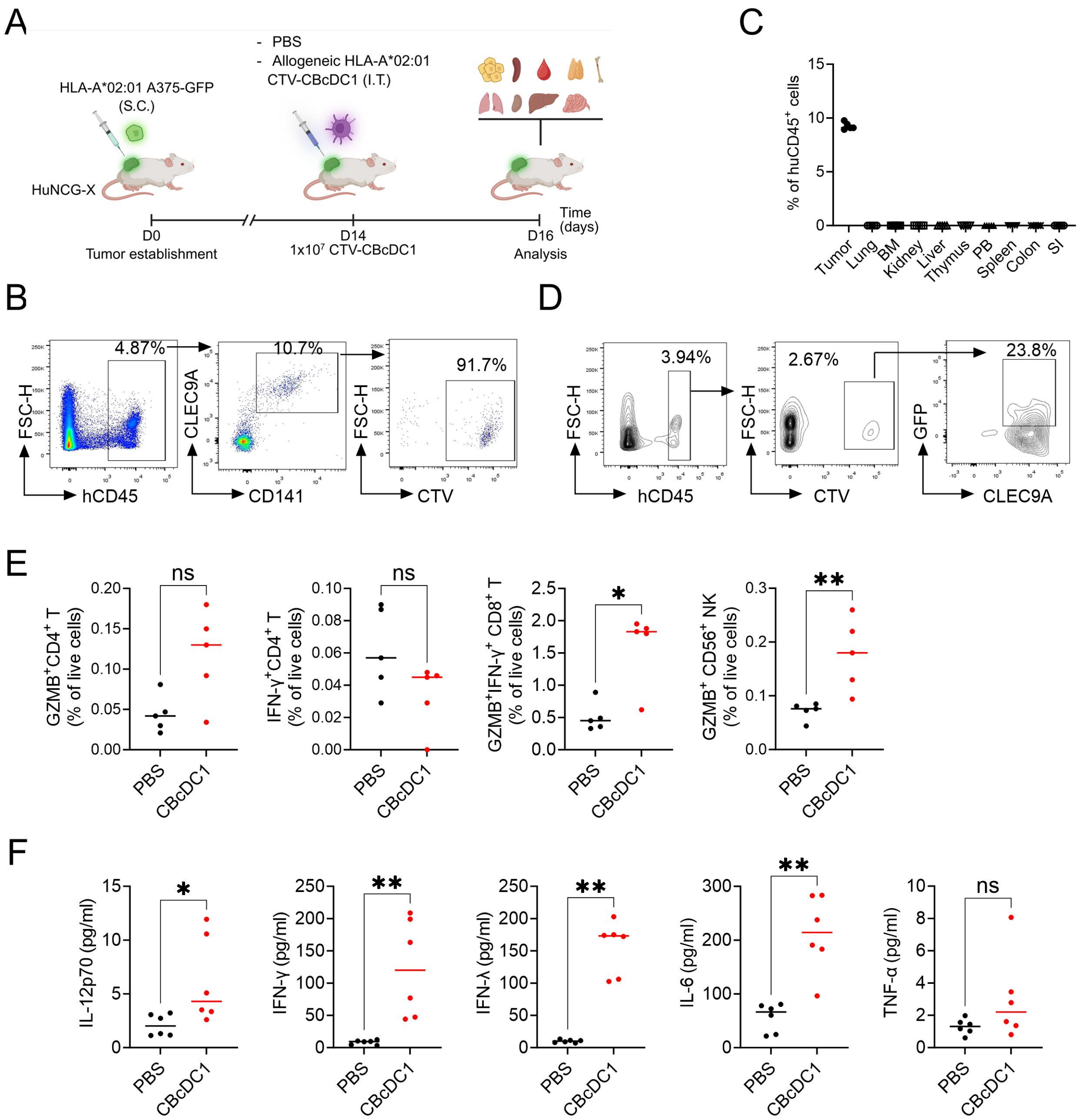


**Fig. S11. CBcDC1 Predominantly Functions *in-situ* within Tumors.** (**A**) Schematic illustration showing activated allogeneic HLA-A*02:01 CBcDC1 ($1\times10^7$ cells) labeled with CTV, administered via I.T. injection at 14 days post of establishment of $GFP^+$ HLA-A*02:01 A375 tumors. (**B**) *In vivo* distribution of CBcDC1 was analyzed 48 hours after injection. Gating strategies for detecting exogenously injected CBcDC1 in various tissue are presented. (**C**) Quantitative statistical analysis of the distribution of exogenously CBcDC1 in various tissues. (**D**) Analysis of the phagocytic activity of exogenous CBcDC1 *in vivo*. (**E**) Frequency of the indicated cell types in the tumor. N=5. (**F**) Serum levels of IL-12p70, IFN-γ, IFN-λ, IL-6 and TNF-α in huNCG-X mice. N=6. (**E**) and (**F**) data were analyzed by Mann-Whitney U test. $*p<0.05$, $**p<0.01$, ns, no significant.

<u>Related to Fig. 5.</u>

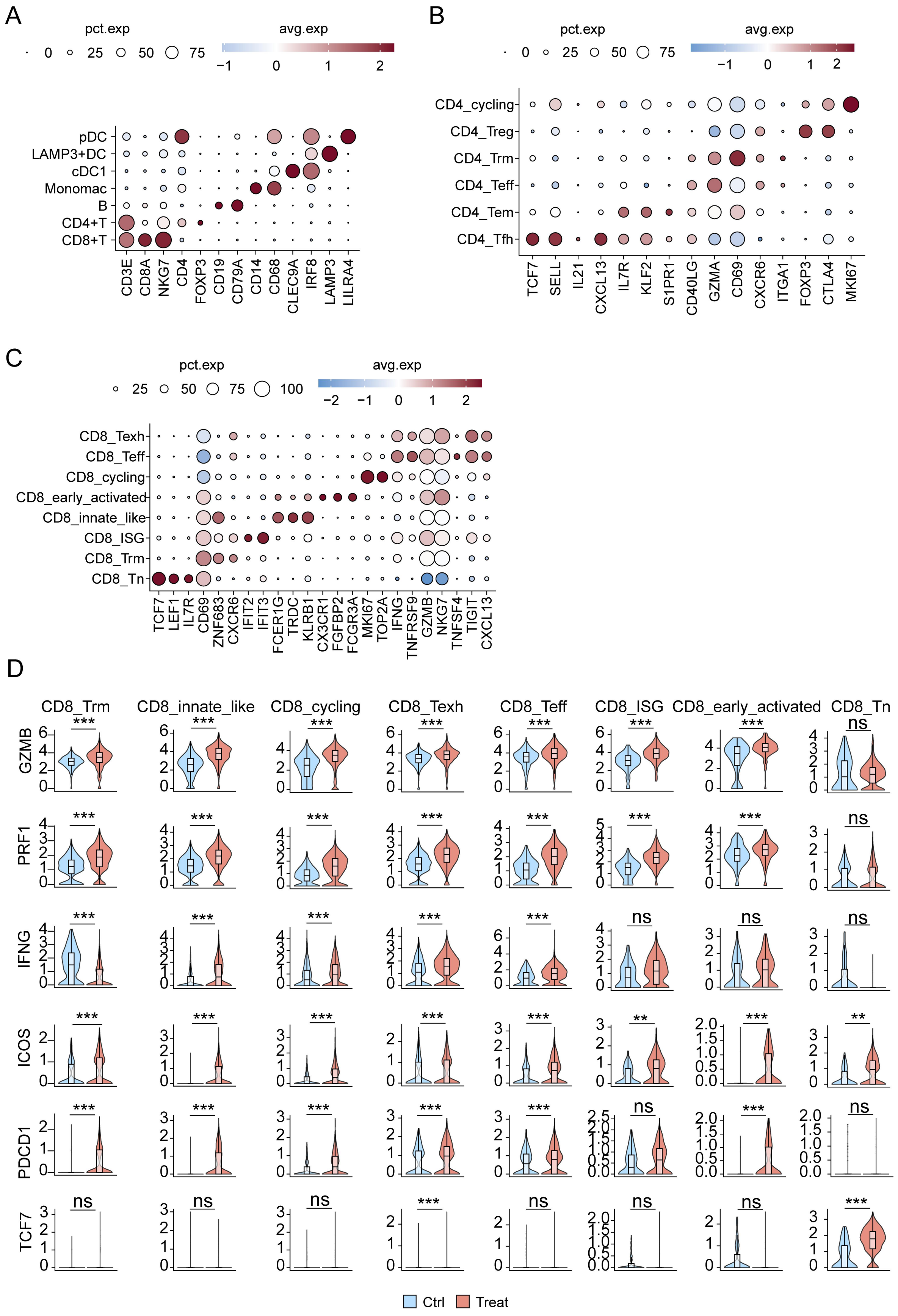
A
pct.exp
0 25 50 75
avg.exp
−1 0 1 2
pDC
LAMP3+DC
cDC1
Monomac
B
CD4+T
CD8+T
CD3E CD8A NKG7 CD4 FOXP3 CD19 CD79A CD14 CD68 CLEC9A IRF8 LAMP3 LILRA4
B
pct.exp
0 25 50 75
avg.exp
−1 0 1 2
CD4_cycling
CD4_Treg
CD4_Trm
CD4_Teff
CD4_Tem
CD4_Tfh
TCF7 SELL IL21 CXCL13 IL7R KLF2 S1PR1 CD40LG GZMA CD69 CXCR6 ITGA1 FOXP3 CTLA4 MKI67
C
pct.exp
25 50 75 100
avg.exp
−2 −1 0 1 2
CD8_Texh
CD8_Teff
CD8_cycling
CD8_early_activated
CD8_innate_like
CD8_ISG
CD8_Trm
CD8_Tn
TCF7 LEF1 IL7R CD69 ZNF683 CXCR6 IFIT2 IFIT3 FCER1G TRDC KLRB1 CX3CR1 FGFBP2 FCGR3A MKI67 TOP2A IFNG TNFRSF9 GZMB NKG7 TNFSF4 TIGIT CXCL13
D
CD8_Trm CD8_innate_like CD8_cycling CD8_Texh CD8_Teff CD8_ISG CD8_early_activated CD8_Tn
GZMB *** *** *** *** *** *** *** ns
PRF1 *** *** *** *** *** *** *** ns
IFNG *** *** *** *** *** ns ns ns
ICOS *** *** *** *** *** ** *** **
PDCD1 *** *** *** *** *** ns *** ns
TCF7 ns ns ns *** ns ns ns ***
Ctrl
Treat

**Fig. S12. Single-Cell Analysis of Human CD45$^+$ Cells in Tumor Tissues.** (**A**) Dot plot showing marker gene expression profiles used for cell type clustering. (**B**, **C**) Dot plots displaying marker gene expression for CD8$^+$ T cell (**B**) and CD4$^+$ T cell (**C**) sub clustering. (**D**) Violin plots depicting the expression levels of *GZMB*, *PRF1*, *IFNG*, *ICOS*, *PDCD1*, and *TCF7* in CD8$^+$ T cell subclusters. **D** data were analyzed using the Wilcoxon rank-sum test. **p<0.01, ***p<0.001, ns, no significant.
<u>Related to Fig. 6.</u>

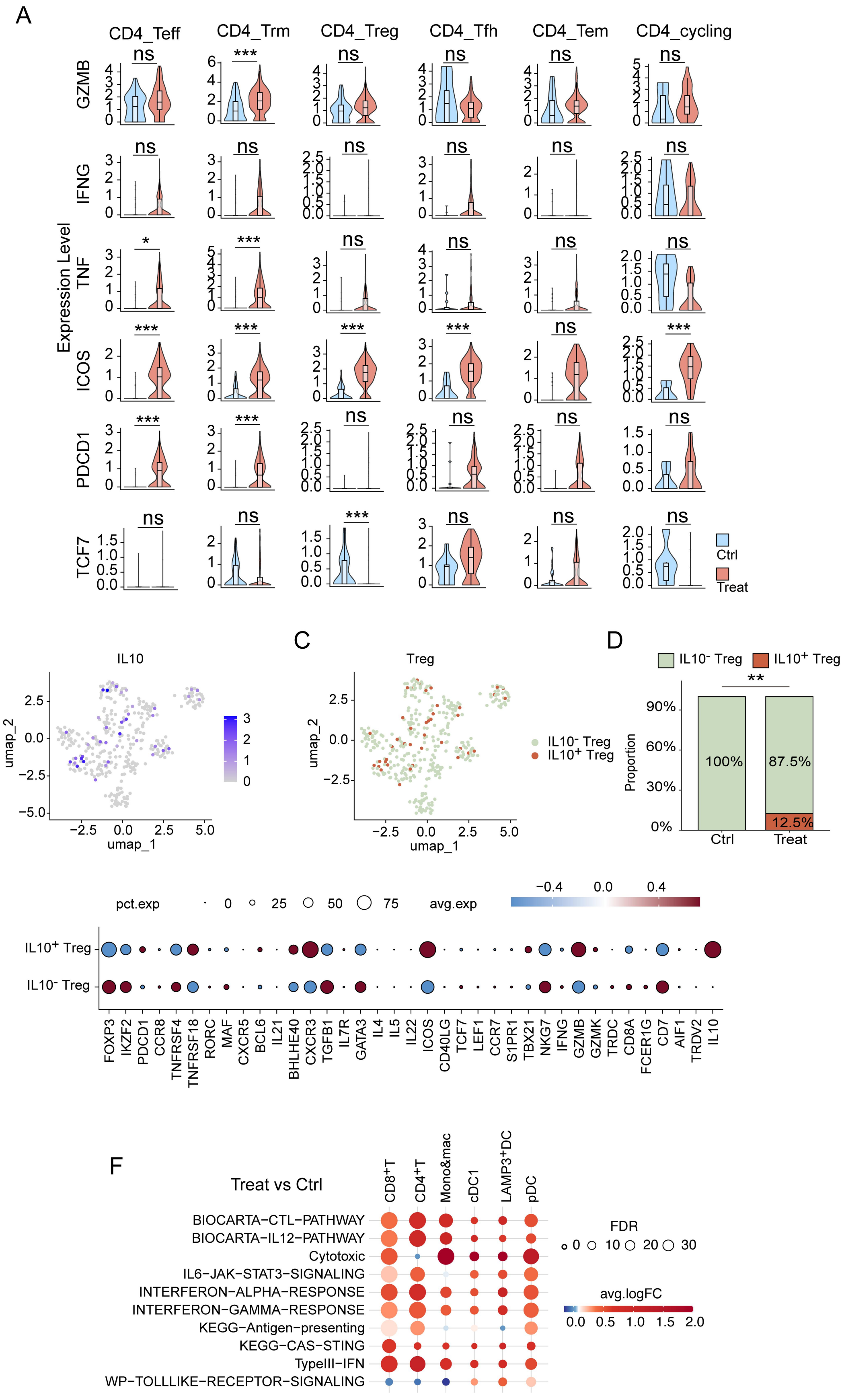
A
CD4_Teff
CD4_Trm
CD4_Treg
CD4_Tfh
CD4_Tem
CD4_cycling
GZMB
IFNG
TNF
ICOS
PDCD1
TCF7
Expression Level
Ctrl
Treat
B
IL10
umap_1
umap_2
C
Treg
IL10⁻ Treg
IL10⁺ Treg
D
Proportion
100%
87.5%
12.5%
Ctrl
Treat
E
pct.exp
avg.exp
IL10⁺ Treg
IL10⁻ Treg
FOXP3
IKZF2
PDCD1
CCR8
TNFRSF4
TNFRSF18
RORC
MAF
CXCR5
BCL6
IL21
BHLHE40
CXCR3
TGFB1
IL7R
GATA3
IL4
IL5
IL22
ICOS
CD40LG
TCF7
LEF1
CCR7
S1PR1
TBX21
NKG7
IFNG
GZMB
GZMK
TRDC
CD8A
FCER1G
CD7
AIF1
TRDV2
IL10
F
Treat vs Ctrl
CD8⁺T
CD4⁺T
Mono&mac
cDC1
LAMP3⁺DC
pDC
BIOCARTA−CTL−PATHWAY
BIOCARTA−IL12−PATHWAY
Cytotoxic
IL6−JAK−STAT3−SIGNALING
INTERFERON−ALPHA−RESPONSE
INTERFERON−GAMMA−RESPONSE
KEGG−Antigen−presenting
KEGG−CAS−STING
TypeIII−IFN
WP−TOLLLIKE−RECEPTOR−SIGNALING
FDR
avg.logFC

**Fig. S13. CBcDC1 Therapy Significantly Improves the TME.** (**A**) Violin plots depicting the expression of *GZMB*, *IFNG*, *TNF*, *ICOS*, *PDCD1*, and *TCF7* in $CD4^+$ T cell subclusters. (**B**) UMAP plot depicting the expression levels of *IL10* in Treg cells from control (Ctrl) and treatment (Treat) groups. (**C**) UMAP plot showing the distribution of $IL10^+$ and $IL10^-$ Treg subsets. (**D**) The relative proportions of Treg subsets. (**E**) The bubble plot illustrates the gene expression signatures of Treg subsets. (**F**) Enrichment analysis of immune-related pathways in distinct cell clusters from Ctrl and Treat tumors. **A** data were analyzed using the Wilcoxon rank-sum test. **D** data were analyzed using the Pearson's chi-squared ($\chi^2$) test. *$p<0.05$, **$p<0.01$, ***$p<0.001$, ns, no significant.
<u>Related to Fig. 6.</u>

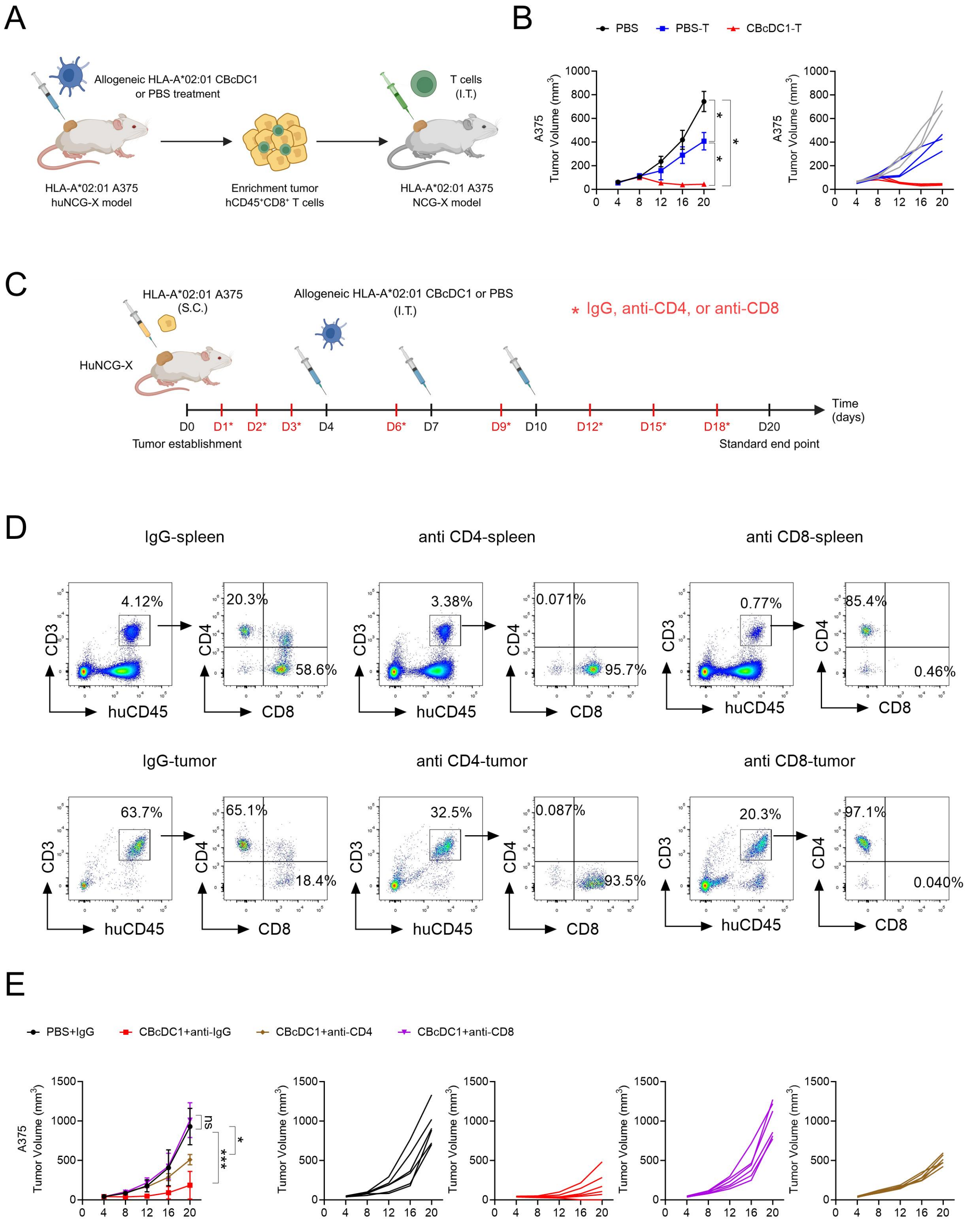


**Fig. S14. Polyclonal CD8$^+$ T Cells are Critical for CBcDC1 Mediated Tumor Control.** (**A**) Schematic representation of the experimental setup. Tumor-infiltrating human CD8$^+$ T cells (huCD45$^+$CD3$^+$CD8$^+$) were isolated from HLA-A*02:01 A375 tumor-bearing huNCG-X mice that had been treated with either PBS (PBS-T) or HLA-A*02:01 CBcDC1 (CBcDC1-T). These

CD8$^+$ T cells were subsequently I.T. transferred into recipient NCG-X mice bearing HLA-A*02:01 A375 tumors ($1\times10^6$ per mouse). (**B**) Tumor growth curves showing mean tumor growth (left) and growth of individual tumors (right). N=3. (**C**) Schematic diagram of T cell depletion experiments. HLA-A*02:01 huNCG-X mice bearing HLA-A*02:01 A375 tumors were intraperitoneally (I.P.) injected with T cell depleting antibodies at the indicated time points (red, 100μg per mouse per injection). Activated allogeneic HLA-A*02:01 CBcDC1 ($1\times10^6$ cells per mouse per injection) were injected into tumors at 4-, 7- and 10-days following tumor establishment. (**D**) Assay for T cell depletion efficiency. (**E**) Tumor growth kinetics, showing mean tumor volume (left) and growth trajectories of individual tumors (right). N=6. (**B**) and (**E**) data are presented as the mean ± SD and were analyzed by two-way ANOVA with Bonferroni post-hoc test. *$p<0.05$, ***$p<0.001$, ns, no significant.

Related to Fig. 7.

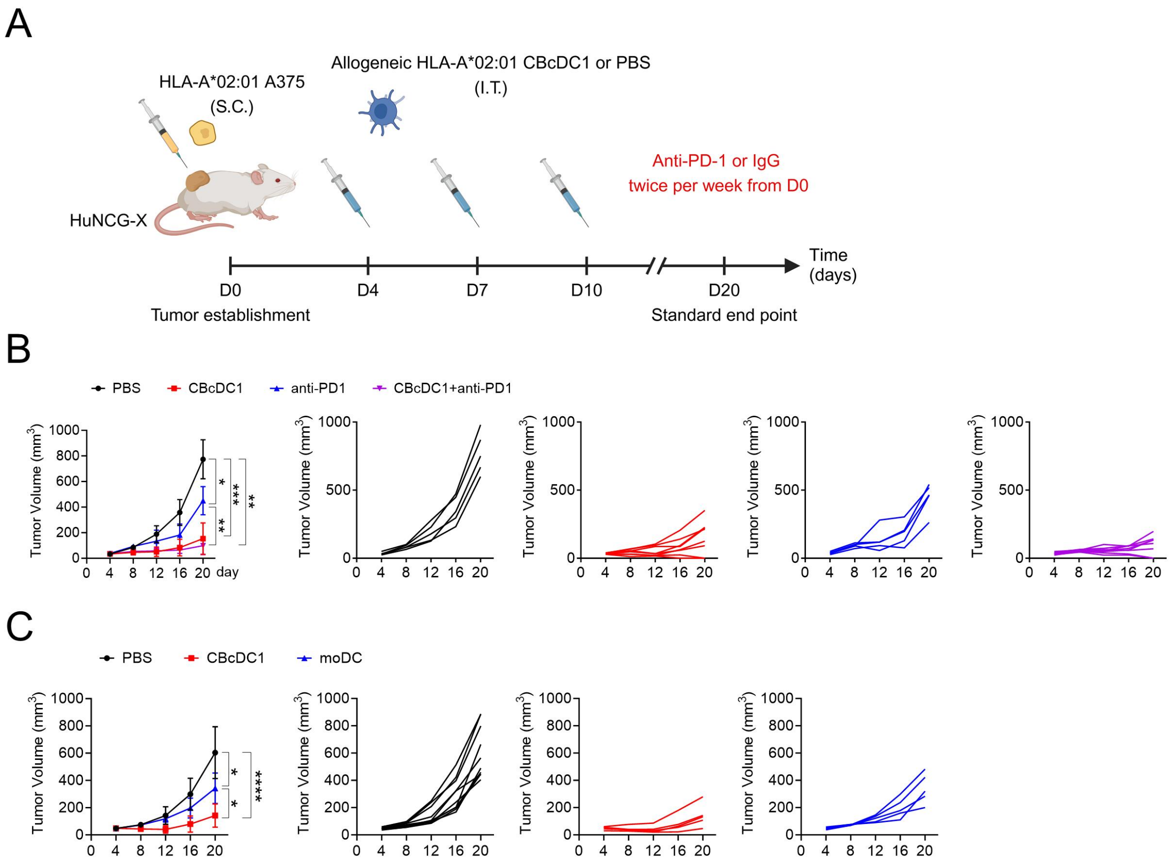


**Fig. S15. CBcDC1 is Superior in Tumor Control than Immune Checkpoint Inhibitor and MoDC.** (**A**) Schematic of the experimental design for combination therapy with CBcDC1 and immune checkpoint blockade in melanoma. Activated allogeneic HLA-A*02:01 CBcDC1 ($1\times10^6$ cells per mouse per injection) were I.T. injected into A375 tumor-bearing huNCG-X mice on day-4, -7, and -10 following tumor establishment. Anti-PD-1 antibody or IgG control (50µg per mice per injection) were I.P. injected into HLA-A*02:01 huNCG-X mice twice per week from day 0. (**B**) Tumor growth curves showing the mean tumor size (left) and growth of individual tumors (right) in the indicated groups. N=5-8. (**C**) Comparison of the anti-tumor efficacy of CBcDC1 and moDC ($1\times10^6$ cells per mouse per injection) in melanoma models. Activated allogeneic HLA-A*02:01 CBcDC1 or moDC pulsed with tumor lysis were administered I.T. on days 4, 7 and 10 post-tumors implantations. Tumor growth kinetics, showing mean tumor volume (left) and growth trajectories of individual tumors (right). N=5-10. (**B**) and (**D**) data are presented as the mean ± SD and were analyzed by two-way ANOVA with Bonferroni post-hoc test. *$p<0.05$, **$p<0.01$, ***$p<0.001$, ****$p<0.0001$.
Related to Fig. 7.

**Table S1.**

**Top 50 Differentially Expressed Genes (DEGs) for Major Immune Cell Clusters and cDC Subclusters.**

**top50_DEG_of_total_cell_cluster**

| p_val | avg_log2FC | pct.1 | pct.2 | p_val_adj | cluster | gene |
|---|---|---|---|---|---|---|
| 0 | 4.94222640293064 | 0.864 | 0.107 | 0 | CBcDC1 | C1orf54 |
| 0 | 4.43457550561405 | 0.975 | 0.239 | 0 | CBcDC1 | IRF8 |
| 0 | 5.46492855102651 | 0.724 | 0.037 | 0 | CBcDC1 | CLEC9A |
| 0 | 3.48014958338817 | 0.828 | 0.149 | 0 | CBcDC1 | NAPSB |
| 0 | 3.43864689033265 | 0.793 | 0.147 | 0 | CBcDC1 | MPEG1 |
| 0 | 2.96091603380635 | 0.945 | 0.301 | 0 | CBcDC1 | S100B |
| 0 | 3.5444059675327 | 0.737 | 0.105 | 0 | CBcDC1 | RGCC |
| 0 | 3.31657495342017 | 0.884 | 0.257 | 0 | CBcDC1 | ID2 |
| 0 | 2.20173195643222 | 0.824 | 0.205 | 0 | CBcDC1 | TGFBI |
| 0 | 4.71810104274104 | 0.658 | 0.042 | 0 | CBcDC1 | RUBCNL |
| 0 | 3.00858249667281 | 0.759 | 0.15 | 0 | CBcDC1 | LGMN |
| 0 | 3.52298436816629 | 0.892 | 0.287 | 0 | CBcDC1 | WDFY4 |
| 0 | 3.34610842764329 | 0.738 | 0.138 | 0 | CBcDC1 | RAB7B |
| 0 | 2.5530945465879 | 0.704 | 0.121 | 0 | CBcDC1 | WFDC21P |
| 0 | 1.74739503232769 | 0.898 | 0.323 | 0 | CBcDC1 | FGL2 |
| 0 | 3.54244661763831 | 0.659 | 0.088 | 0 | CBcDC1 | ZNF366 |
| 0 | 3.45634731794063 | 0.679 | 0.109 | 0 | CBcDC1 | EGLN3 |
| 0 | 3.38369670437993 | 0.666 | 0.109 | 0 | CBcDC1 | CPVL-AS2 |
| 0 | 3.44261611029293 | 0.657 | 0.1 | 0 | CBcDC1 | SHTN1 |
| 0 | 3.26895925315168 | 0.654 | 0.102 | 0 | CBcDC1 | HLA-DOB |
| 0 | 1.53321705557948 | 0.938 | 0.388 | 0 | CBcDC1 | CLEC10A |
| 0 | 2.77032770076695 | 0.928 | 0.38 | 0 | CBcDC1 | BASP1 |
| 0 | 1.5377402050 | 0.95 | 0.40 | 0 | CBcDC1 | HLA-DQA1 |

| | | | | | | |
|---|---|---|---|---|---|---|
| | 0668 | 2 | 4 | | | |
| 0 | 5.06161218339684 | 0.58 | 0.033 | 0 | CBcDC1 | CCND1 |
| 0 | 2.26882031928325 | 0.858 | 0.32 | 0 | CBcDC1 | NAAA |
| 0 | 2.40687460335181 | 0.961 | 0.424 | 0 | CBcDC1 | LMNA |
| 0 | 3.00663549537419 | 0.643 | 0.12 | 0 | CBcDC1 | NDRG2 |
| 0 | 2.40602677439193 | 0.924 | 0.403 | 0 | CBcDC1 | CRIP1 |
| 0 | 3.9605804643793 | 0.58 | 0.059 | 0 | CBcDC1 | TCEA3 |
| 0 | 2.94746333605318 | 0.624 | 0.105 | 0 | CBcDC1 | CLIC2 |
| 0 | 3.65382214345109 | 0.569 | 0.065 | 0 | CBcDC1 | CAMK2D |
| 0 | 2.19206494877354 | 0.974 | 0.473 | 0 | CBcDC1 | HLA-DRB5 |
| 0 | 3.0804600173991 | 0.598 | 0.101 | 0 | CBcDC1 | BATF3 |
| 0 | 2.22604845796856 | 0.658 | 0.161 | 0 | CBcDC1 | CLEC7A |
| 0 | 2.44292039562519 | 0.658 | 0.165 | 0 | CBcDC1 | BCL6 |
| 0 | 2.45414046718789 | 0.815 | 0.33 | 0 | CBcDC1 | TAP2 |
| 0 | 2.95095743479449 | 0.611 | 0.128 | 0 | CBcDC1 | RAB29 |
| 0 | 1.70574164298534 | 0.743 | 0.27 | 0 | CBcDC1 | ARL4C |
| 0 | 1.63936623698475 | 0.86 | 0.392 | 0 | CBcDC1 | HLA-DRB6 |
| 0 | 1.71090491295018 | 0.647 | 0.183 | 0 | CBcDC1 | HLA-DQA2 |
| 0 | 1.96073875382763 | 0.784 | 0.322 | 0 | CBcDC1 | DSE |
| 0 | 1.01277062881507 | 0.839 | 0.381 | 0 | CBcDC1 | HLA-DMB |
| 0 | 2.17492411750976 | 0.661 | 0.205 | 0 | CBcDC1 | TSPAN33 |
| 0 | 1.92635121676535 | 0.581 | 0.127 | 0 | CBcDC1 | C1QB |
| 0 | 2.60642570994352 | 0.971 | 0.518 | 0 | CBcDC1 | TAP1 |
| 0 | 2.19753872058926 | 0.978 | 0.528 | 0 | CBcDC1 | ANXA2 |
| 0 | 2.2233661217069 | 0.69 | 0.252 | 0 | CBcDC1 | LIMA1 |
| 0 | 3.15757929841113 | 0.511 | 0.074 | 0 | CBcDC1 | PLBD1 |
| 0 | 3.21468260743868 | 0.494 | 0.061 | 0 | CBcDC1 | SLAMF8 |

| | | | | | | |
|---|---|---|---|---|---|---|
| 0 | 5.48689273702558 | 0.447 | 0.014 | 0 | CBcDC1 | ENPP1 |
| 0 | 4.01060316674232 | 0.972 | 0.203 | 0 | CBcDC2 | MRC1 |
| 0 | 3.19353611209464 | 0.966 | 0.333 | 0 | CBcDC2 | CD1C |
| 0 | 3.31191450127278 | 0.847 | 0.217 | 0 | CBcDC2 | ENSG00000284697 |
| 0 | 3.16095681349777 | 0.93 | 0.305 | 0 | CBcDC2 | NCF2 |
| 0 | 3.5165390450239 | 0.735 | 0.115 | 0 | CBcDC2 | IGSF6 |
| 0 | 3.44040584784638 | 0.753 | 0.147 | 0 | CBcDC2 | CTSH |
| 0 | 3.11013178920888 | 0.711 | 0.148 | 0 | CBcDC2 | CDKN1A |
| 0 | 1.95321370364056 | 0.926 | 0.365 | 0 | CBcDC2 | FGL2 |
| 0 | 3.03527211094986 | 0.754 | 0.193 | 0 | CBcDC2 | PKIB |
| 0 | 2.38726592229694 | 0.98 | 0.425 | 0 | CBcDC2 | CLEC10A |
| 0 | 2.65346864860094 | 0.916 | 0.379 | 0 | CBcDC2 | JAML |
| 0 | 3.26160751090905 | 0.655 | 0.123 | 0 | CBcDC2 | CLEC4A |
| 0 | 4.85136524271326 | 0.575 | 0.046 | 0 | CBcDC2 | CD1B |
| 0 | 2.24244755535816 | 0.97 | 0.445 | 0 | CBcDC2 | HLA-DQA1 |
| 0 | 2.35074043994971 | 0.914 | 0.405 | 0 | CBcDC2 | HLA-DMB |
| 0 | 4.23420979617148 | 0.558 | 0.05 | 0 | CBcDC2 | CD1E |
| 0 | 2.35780695452681 | 0.795 | 0.3 | 0 | CBcDC2 | ALOX5AP |
| 0 | 3.02214233026588 | 0.604 | 0.117 | 0 | CBcDC2 | DOK2 |
| 0 | 3.15700380454405 | 0.613 | 0.127 | 0 | CBcDC2 | CD1A |
| 0 | 2.62601553850679 | 0.818 | 0.332 | 0 | CBcDC2 | GGTA1 |
| 0 | 3.10295031658795 | 0.804 | 0.319 | 0 | CBcDC2 | IFITM3 |
| 0 | 2.60802105348033 | 0.629 | 0.147 | 0 | CBcDC2 | MS4A6A |
| 0 | 3.99903279217894 | 0.544 | 0.07 | 0 | CBcDC2 | ENSG00000272980 |
| 0 | 1.69912993322858 | 0.737 | 0.27 | 0 | CBcDC2 | TGFBI |
| 0 | 4.36136286890328 | 0.515 | 0.053 | 0 | CBcDC2 | FCGR2B |
| 0 | 2.0648277080 | 0.67 | 0.21 | 0 | CBcDC2 | HLA-DQA2 |

| | | | | | | |
|---|---|---|---|---|---|---|
| | 0423 | 5 | 6 | | | |
| 0 | 2.61003358400674 | 0.625 | 0.172 | 0 | CBcDC2 | HLA-DOA |
| 0 | 2.88718918462541 | 0.573 | 0.126 | 0 | CBcDC2 | IL10RA |
| 0 | 1.57129891071758 | 0.96 | 0.516 | 0 | CBcDC2 | HLA-DRB5 |
| 0 | 1.76182427334012 | 0.871 | 0.428 | 0 | CBcDC2 | HLA-DRB6 |
| 0 | 5.53596001312342 | 0.462 | 0.02 | 0 | CBcDC2 | FPR3 |
| 0 | 2.0201515824978 | 0.698 | 0.257 | 0 | CBcDC2 | CD86 |
| 0 | 2.36920097502615 | 0.735 | 0.297 | 0 | CBcDC2 | PEA15 |
| 0 | 2.27478090967307 | 0.869 | 0.436 | 0 | CBcDC2 | FCER1G |
| 0 | 2.49751101209717 | 0.973 | 0.541 | 0 | CBcDC2 | ENSG00000268173 |
| 0 | 2.7363002158408 | 0.531 | 0.101 | 0 | CBcDC2 | CSF1R |
| 0 | 1.98878444783159 | 0.743 | 0.313 | 0 | CBcDC2 | CIITA |
| 0 | 2.85323670587195 | 0.568 | 0.142 | 0 | CBcDC2 | LGALS3 |
| 0 | 2.76166973194758 | 0.967 | 0.541 | 0 | CBcDC2 | TYROBP |
| 0 | 1.75915996113558 | 0.922 | 0.502 | 0 | CBcDC2 | SAMHD1 |
| 0 | 2.649801806 | 0.553 | 0.134 | 0 | CBcDC2 | ASGR2 |
| 0 | 1.59622657917179 | 0.748 | 0.331 | 0 | CBcDC2 | LY86 |
| 0 | 2.48169907770401 | 0.652 | 0.236 | 0 | CBcDC2 | SOAT1 |
| 0 | 2.10648771404841 | 0.651 | 0.239 | 0 | CBcDC2 | ADAM8 |
| 0 | 1.67269644916486 | 0.72 | 0.312 | 0 | CBcDC2 | ARL4C |
| 0 | 2.93142460579891 | 0.536 | 0.129 | 0 | CBcDC2 | C1QA |
| 0 | 2.49175747969222 | 0.704 | 0.299 | 0 | CBcDC2 | SLC2A3 |
| 0 | 2.01551788637981 | 0.622 | 0.22 | 0 | CBcDC2 | TYMP |
| 0 | 3.81655639362875 | 0.444 | 0.046 | 0 | CBcDC2 | IL1R1 |
| 0 | 1.84094484246124 | 0.836 | 0.443 | 0 | CBcDC2 | RAB31 |
| 0 | 1.41024249627576 | 0.816 | 0.287 | 0 | CD1C$^+$ cell | MRC1 |
| 0 | 2.03094007071106 | 0.784 | 0.299 | 0 | CD1C$^+$ cell | MKI67 |

| | | | | | | |
|---|---|---|---|---|---|---|
| 0 | 1.59481319727029 | 0.858 | 0.417 | 0 | CD1C$^+$ cell | TOP2A |
| 0 | 1.85847606801496 | 0.619 | 0.195 | 0 | CD1C$^+$ cell | MS4A6A |
| 0 | 1.65907726204153 | 0.759 | 0.34 | 0 | CD1C$^+$ cell | CCNB2 |
| 0 | 1.26033733797325 | 0.782 | 0.371 | 0 | CD1C$^+$ cell | LY86 |
| 0 | 1.69303441194632 | 0.693 | 0.293 | 0 | CD1C$^+$ cell | UBE2C |
| 0 | 1.50398603248483 | 0.668 | 0.272 | 0 | CD1C$^+$ cell | AURKB |
| 0 | 1.91873612670698 | 0.634 | 0.24 | 0 | CD1C$^+$ cell | CDC20 |
| 0 | 1.56959174804983 | 0.652 | 0.269 | 0 | CD1C$^+$ cell | CDKN3 |
| 0 | 1.44672999952096 | 0.637 | 0.264 | 0 | CD1C$^+$ cell | CCNA2 |
| 0 | 1.78074663017988 | 0.554 | 0.185 | 0 | CD1C$^+$ cell | HMMR |
| 0 | 1.48687404632083 | 0.644 | 0.278 | 0 | CD1C$^+$ cell | TPX2 |
| 0 | 1.43904822984802 | 0.637 | 0.279 | 0 | CD1C$^+$ cell | CENPF |
| 0 | 1.04166652897197 | 0.896 | 0.545 | 0 | CD1C$^+$ cell | SAMHD1 |
| 0 | 1.32405495336663 | 0.68 | 0.33 | 0 | CD1C$^+$ cell | ENSG00000285920 |
| 0 | 1.14367347785208 | 0.828 | 0.481 | 0 | CD1C$^+$ cell | CKS1B |
| 0 | 1.63329449842154 | 0.6 | 0.254 | 0 | CD1C$^+$ cell | CCNB1 |
| 0 | 1.90611321437867 | 0.506 | 0.162 | 0 | CD1C$^+$ cell | CEP55 |
| 0 | 1.41206711602547 | 0.629 | 0.287 | 0 | CD1C$^+$ cell | CDK1 |
| 0 | 1.37859912160796 | 0.614 | 0.274 | 0 | CD1C$^+$ cell | BIRC5 |
| 0 | 1.5461725162841 | 0.569 | 0.231 | 0 | CD1C$^+$ cell | NCAPG |
| 0 | 1.47768426006513 | 0.544 | 0.211 | 0 | CD1C$^+$ cell | GAS6 |
| 0 | 1.50667626453469 | 0.57 | 0.239 | 0 | CD1C$^+$ cell | PRC1 |
| 0 | 1.33226304121879 | 0.759 | 0.429 | 0 | CD1C$^+$ cell | PTTG1 |
| 0 | 1.52434238676807 | 0.475 | 0.146 | 0 | CD1C$^+$ cell | CSF1R |
| 0 | 1.37937888347782 | 0.562 | 0.234 | 0 | CD1C$^+$ cell | TASL |
| 0 | 1.34545489327713 | 0.521 | 0.193 | 0 | CD1C$^+$ cell | MS4A7 |
| 0 | 2.2610532371 | 0.41 | 0.08 | 0 | CD1C$^+$ cell | CCR2 |

| | | | | | | |
|---|---|---|---|---|---|---|
| | 9621 | 4 | 8 | | | |
| 0 | 1.59291220519766 | 0.514 | 0.192 | 0 | CD1C+ cell | KIF2C |
| 0 | 1.2107983381036 | 0.775 | 0.457 | 0 | CD1C+ cell | TACC3 |
| 0 | 1.7567806991786 | 0.499 | 0.183 | 0 | CD1C+ cell | CENPE |
| 0 | 1.82290363139729 | 0.476 | 0.162 | 0 | CD1C+ cell | PLK1 |
| 0 | 1.51698307168122 | 0.505 | 0.193 | 0 | CD1C+ cell | NUF2 |
| 0 | 1.7960281057026 | 0.457 | 0.149 | 0 | CD1C+ cell | DLGAP5 |
| 0 | 1.72163627559976 | 0.459 | 0.156 | 0 | CD1C+ cell | CDCA3 |
| 0 | 2.10098981106157 | 0.426 | 0.129 | 0 | CD1C+ cell | GTSE1 |
| 0 | 1.78571739361426 | 0.389 | 0.112 | 0 | CD1C+ cell | PLD4 |
| 0 | 1.26846041943059 | 0.878 | 0.608 | 0 | CD1C+ cell | TUBB4B |
| 0 | 1.96113806762654 | 0.297 | 0.063 | 0 | CD1C+ cell | CD180 |
| 0 | 2.00499856140962 | 0.286 | 0.064 | 0 | CD1C+ cell | TMEM236 |
| 0 | 1.26242168658779 | 0.99 | 0.807 | 0 | CD1C+ cell | HMGB2 |
| 2.69985595753762e-295 | 1.59797690470994 | 0.329 | 0.093 | 1.04935301501615e-290 | CD1C+ cell | FCER2 |
| 9.73434061642805e-291 | 1.11755161162581 | 0.507 | 0.188 | 3.78344616738709e-286 | CD1C+ cell | IGSF6 |
| 3.13215847136606e-290 | 1.35132868718366 | 0.709 | 0.404 | 1.21737603306585e-285 | CD1C+ cell | KPNA2 |
| 1.13595420413437e-289 | 1.14048640635569 | 0.896 | 0.683 | 4.41511320520906e-285 | CD1C+ cell | H4C3 |
| 1.59980469119741e-289 | 1.68175883439295 | 0.274 | 0.068 | 6.21796089327699e-285 | CD1C+ cell | CYBB |
| 3.79412386205992e-282 | 1.04879190356364 | 0.838 | 0.607 | 1.47466212146683e-277 | CD1C+ cell | ACAT2 |
| 9.09850371651844e-281 | 1.53303372708818 | 0.394 | 0.132 | 3.53631543949922e-276 | CD1C+ cell | DAB2 |
| 3.5559472426241e-280 | 1.40641223839443 | 0.538 | 0.231 | 1.38209001479071e-275 | CD1C+ cell | CLECL1 |
| 0 | 8.91711763670108 | 0.973 | 0.034 | 0 | LAMP3+ DC | CCL17 |
| 0 | 8.61239386484421 | 0.883 | 0.009 | 0 | LAMP3+ DC | LAMP3 |
| 0 | 8.81870617028292 | 0.856 | 0.011 | 0 | LAMP3+ DC | CCR7 |
| 0 | 5.67905365530682 | 0.899 | 0.063 | 0 | LAMP3+ DC | IL4I1 |
| 0 | 6.14787261861533 | 0.91 | 0.089 | 0 | LAMP3+ DC | RGS1 |

| | | | | | | |
|---|---|---|---|---|---|---|
| 0 | 5.27500149690199 | 0.771 | 0.06 | 0 | LAMP3$^+$ DC | IRF4 |
| 0 | 4.5707808892924 | 0.691 | 0.04 | 0 | LAMP3$^+$ DC | NR4A3 |
| 0 | 10.9013484135136 | 0.606 | 0.002 | 0 | LAMP3$^+$ DC | UBD |
| 0 | 5.18764635680755 | 0.628 | 0.035 | 0 | LAMP3$^+$ DC | CD274 |
| 0 | 9.18062368356098 | 0.58 | 0.004 | 0 | LAMP3$^+$ DC | CCL22 |
| 0 | 6.10827755617555 | 0.569 | 0.022 | 0 | LAMP3$^+$ DC | TNFRSF11A |
| 0 | 7.08646978015299 | 0.532 | 0.008 | 0 | LAMP3$^+$ DC | NRP2 |
| 0 | 5.71121913267117 | 0.516 | 0.021 | 0 | LAMP3$^+$ DC | LIMCH1 |
| 0 | 11.0577240378603 | 0.495 | 0.002 | 0 | LAMP3$^+$ DC | TUBB2B |
| 0 | 5.0830513137839 | 0.473 | 0.023 | 0 | LAMP3$^+$ DC | ENSG00000288528 |
| 0 | 8.51467131776112 | 0.441 | 0.003 | 0 | LAMP3$^+$ DC | TMEM176A |
| 0 | 6.76081955892786 | 0.42 | 0.007 | 0 | LAMP3$^+$ DC | TMEM176B |
| 0 | 5.59286466231135 | 0.388 | 0.008 | 0 | LAMP3$^+$ DC | SLCO5A1 |
| 0 | 4.92288462842006 | 0.34 | 0.007 | 0 | LAMP3$^+$ DC | IL32 |
| 0 | 6.42622808941789 | 0.335 | 0.007 | 0 | LAMP3$^+$ DC | LINC01857 |
| 0 | 5.72788917482556 | 0.293 | 0.007 | 0 | LAMP3$^+$ DC | TMCC3 |
| 0 | 6.79620941039168 | 0.25 | 0.003 | 0 | LAMP3$^+$ DC | TNFRSF9 |
| 0 | 11.1005669014143 | 0.245 | 0.001 | 0 | LAMP3$^+$ DC | TFPI2 |
| 0 | 10.7076776900002 | 0.239 | 0 | 0 | LAMP3$^+$ DC | CCL19 |
| 0 | 6.28595283919561 | 0.239 | 0.005 | 0 | LAMP3$^+$ DC | SOCAR |
| 0 | 7.76766063775486 | 0.223 | 0.002 | 0 | LAMP3$^+$ DC | EBI3 |
| 0 | 6.34558937860824 | 0.223 | 0.005 | 0 | LAMP3$^+$ DC | LAMB1 |
| 0 | 9.65108069676018 | 0.213 | 0 | 0 | LAMP3$^+$ DC | NCCRP1 |
| 0 | 9.31959846682434 | 0.207 | 0 | 0 | LAMP3$^+$ DC | LAD1 |
| 0 | 8.01731210107551 | 0.176 | 0.001 | 0 | LAMP3$^+$ DC | OLAH |
| 0 | 7.35537164639429 | 0.17 | 0.002 | 0 | LAMP3$^+$ DC | LORICRIN |
| 0 | 6.7726798722 | 0.16 | 0.00 | 0 | LAMP3$^+$ DC | MSC |

| | | | | | | |
|---|---|---|---|---|---|---|
| | 7047 | 5 | 2 | | | |
| 0 | 7.74281049656673 | 0.112 | 0.001 | 0 | LAMP3+ DC | CADM3 |
| 0 | 8.42349500780659 | 0.096 | 0 | 0 | LAMP3+ DC | CLLU1-AS1 |
| 0 | 7.42973931431783 | 0.085 | 0.001 | 0 | LAMP3+ DC | DCLRE1CP1 |
| 4.43348578416563e-289 | 5.64898260073051 | 0.968 | 0.148 | 1.72316291973165e-284 | LAMP3+ DC | BIRC3 |
| 1.46440860840387e-261 | 6.73351398322385 | 0.117 | 0.002 | 5.69171693828332e-257 | LAMP3+ DC | OR2I1P |
| 2.9856512599948e-260 | 4.61539914460979 | 0.457 | 0.029 | 1.16043307522218e-255 | LAMP3+ DC | PTGIR |
| 4.85899216268339e-247 | 4.77957854589478 | 0.527 | 0.041 | 1.88854448387015e-242 | LAMP3+ DC | SPON2 |
| 3.19797251687277e-242 | 6.2939313173566 | 0.181 | 0.005 | 1.24295597813294e-237 | LAMP3+ DC | HSD11B1-AS1 |
| 7.40694488824206e-242 | 4.60004512166462 | 0.372 | 0.02 | 2.87885726971304e-237 | LAMP3+ DC | GPR157 |
| 1.58676871026948e-220 | 5.52657518686997 | 0.261 | 0.011 | 6.16729394620437e-216 | LAMP3+ DC | DEPP1 |
| 7.23007338860053e-216 | 4.11202140694337 | 0.479 | 0.038 | 2.81011262394737e-211 | LAMP3+ DC | CD80 |
| 3.25134967398934e-207 | 4.20832990806213 | 0.606 | 0.065 | 1.26370207778944e-202 | LAMP3+ DC | MREG |
| 5.06289437821515e-193 | 4.17762013509929 | 0.819 | 0.143 | 1.96779515798088e-188 | LAMP3+ DC | RASSF4 |
| 5.37792625134632e-190 | 4.3856759437694 | 0.686 | 0.095 | 2.09023859611077e-185 | LAMP3+ DC | LY75 |
| 7.44586553546747e-186 | 4.82681652386892 | 0.271 | 0.014 | 2.89398455767014e-181 | LAMP3+ DC | BICDL1 |
| 3.08609002787903e-185 | 4.41909660959643 | 0.527 | 0.054 | 1.19947061113574e-180 | LAMP3+ DC | TRAF1 |
| 4.0024492932206e-185 | 4.00062443530221 | 0.644 | 0.081 | 1.55563196679605e-180 | LAMP3+ DC | PALLD |
| 7.85811173759333e-185 | 4.22841043803355 | 0.601 | 0.072 | 3.0542122890504e-180 | LAMP3+ DC | ETV3 |
| 0 | 3.61151219183921 | 0.727 | 0.093 | 0 | DC_progenitors | JCHAIN |
| 0 | 3.40780179552812 | 0.883 | 0.46 | 0 | DC_progenitors | LTB |
| 0 | 5.00131132414871 | 0.397 | 0.029 | 0 | DC_progenitors | ACY3 |
| 0 | 3.1516161959793 | 0.363 | 0.04 | 0 | DC_progenitors | ENSG00000286848 |
| 0 | 6.74427944338701 | 0.329 | 0.008 | 0 | DC_progenitors | MS4A1 |
| 0 | 6.55788391374275 | 0.235 | 0.011 | 0 | DC_progenitors | IGKC |
| 0 | 4.71825408002944 | 0.23 | 0.018 | 0 | DC_progenitors | SPIB |
| 0 | 5.17984473271147 | 0.155 | 0.007 | 0 | DC_progenitors | NIBAN3 |

| | | | | | | |
|---|---|---|---|---|---|---|
| 0 | 7.24970063947907 | 0.106 | 0.003 | 0 | DC_progenitors | AGR2 |
| 1.21960157755601e-288 | 3.09257007523523 | 0.439 | 0.088 | 4.74022545148692e-284 | DC_progenitors | BLNK |
| 2.90938695492011e-213 | 3.51955890395257 | 0.495 | 0.15 | 1.1307914277688e-208 | DC_progenitors | TCF4 |
| 7.77285486465264e-208 | 3.35327333595719 | 0.259 | 0.042 | 3.02107550024454e-203 | DC_progenitors | TNNI2 |
| 3.70642443383939e-203 | 7.21681638507934 | 0.053 | 0.001 | 1.44057598470035e-198 | DC_progenitors | GZMB |
| 8.52916834649496e-198 | 2.70118934514409 | 0.44 | 0.121 | 3.3150318612322e-193 | DC_progenitors | PLD4 |
| 1.66311557989696e-194 | 3.45159310212343 | 0.341 | 0.075 | 6.46403132438552e-190 | DC_progenitors | IGHM |
| 2.86716200614786e-192 | 2.13048987761251 | 0.62 | 0.239 | 1.11437985692949e-187 | DC_progenitors | CLECL1 |
| 4.58428892191262e-189 | 7.45073117271788 | 0.026 | 0 | 1.78177557527978e-184 | DC_progenitors | TNFRSF17 |
| 1.31957392518096e-177 | 4.6482334538641 | 0.1 | 0.007 | 5.12878797500082e-173 | DC_progenitors | KCNA5 |
| 9.86827837545833e-161 | 1.22402985186329 | 0.909 | 0.825 | 3.83550375618939e-156 | DC_progenitors | GPX1 |
| 2.79245257117892e-151 | 2.13348246323848 | 0.576 | 0.241 | 1.08534254084011e-146 | DC_progenitors | CYTH4 |
| 1.61982245548671e-131 | 2.46117274421402 | 0.114 | 0.013 | 6.29576393774021e-127 | DC_progenitors | CCR7 |
| 8.99993462308602e-129 | 1.54639431475485 | 0.67 | 0.311 | 3.49800458995484e-124 | DC_progenitors | SPINK2 |
| 1.75389310234921e-121 | 2.06743230554745 | 0.579 | 0.277 | 6.81685632090067e-117 | DC_progenitors | MEF2C |
| 1.88524849409505e-117 | 5.85707700448436 | 0.048 | 0.002 | 7.32739532199923e-113 | DC_progenitors | ITGAD |
| 2.06089602284432e-112 | 5.26226034676527 | 0.029 | 0.001 | 8.01008457198902e-108 | DC_progenitors | IGHG3 |
| 5.310072028708e-111 | 3.97625705891932 | 0.09 | 0.01 | 2.06386569539794e-106 | DC_progenitors | TLR7 |
| 2.11816795399199e-110 | 2.35998783938257 | 0.31 | 0.092 | 8.23268338678066e-106 | DC_progenitors | BMF |
| 5.00266248827771e-110 | 3.32401589678989 | 0.176 | 0.034 | 1.9443848293189e-105 | DC_progenitors | MEF2B |
| 2.00705292558509e-104 | 1.69831388922177 | 0.543 | 0.259 | 7.80081260587156e-100 | DC_progenitors | ITGB7 |
| 2.28487712544222e-102 | 1.38390892475943 | 0.733 | 0.504 | 8.88063192345627e-98 | DC_progenitors | LAT2 |
| 2.35288758301812e-101 | 2.82282631253611 | 0.194 | 0.043 | 9.14496816891652e-97 | DC_progenitors | TLR10 |
| 9.05335168181938e-98 | 3.75101640929406 | 0.067 | 0.006 | 3.51876619817274e-93 | DC_progenitors | MIR4432HG |
| 3.03925242843643e-96 | 1.06432112745706 | 0.786 | 0.534 | 1.18126624136039e-91 | DC_progenitors | LSP1 |
| 3.71125637788371e-96 | 5.68226909241345 | 0.032 | 0.001 | 1.44245401639206e-91 | DC_progenitors | ENSG00000290104 |
| 5.80437624811906 | 5.5674885844 | 0.02 | 0.00 | 2.25598691635644 | DC_progenito | ENSG000002 |

| | | | | | | |
|---|---|---|---|---|---|---|
| e-96 | 6231 | 6 | 1 | e-91 | rs | 30631 |
| 1.07136608388248e-95 | 2.99181052978542 | 0.159 | 0.031 | 4.16407855822603e-91 | DC_progenitors | DTX4 |
| 5.45839669552585e-95 | 3.11953400362246 | 0.145 | 0.027 | 2.12151504365003e-90 | DC_progenitors | LPAR3 |
| 8.19587020341675e-91 | 2.42312526553058 | 0.268 | 0.082 | 3.18548887196199e-86 | DC_progenitors | MGAT3 |
| 6.00628540713571e-90 | 1.58925841905901 | 0.557 | 0.295 | 2.33446294919144e-85 | DC_progenitors | LPXN |
| 7.04738822162474e-86 | 2.05382771811286 | 0.363 | 0.141 | 2.73910838009889e-81 | DC_progenitors | DAB2 |
| 8.65351642841643e-84 | 1.94151153614835 | 0.536 | 0.286 | 3.36336223023262e-79 | DC_progenitors | ITGAL |
| 3.12561662011502e-79 | 5.15334546671659 | 0.054 | 0.005 | 1.2148334117401e-74 | DC_progenitors | TCF7 |
| 1.14847635565472e-75 | 1.17114729375767 | 0.782 | 0.558 | 4.46378305152321e-71 | DC_progenitors | SAMHD1 |
| 9.41023844592106e-75 | 2.56933339273147 | 0.197 | 0.054 | 3.65747737677614e-70 | DC_progenitors | PPP1R16B |
| 9.4311700141717e-71 | 2.37336039208484 | 0.261 | 0.09 | 3.66561284940811e-66 | DC_progenitors | SFMBT2 |
| 1.52660034808231e-70 | 5.99906778162373 | 0.02 | 0.001 | 5.93343757289152e-66 | DC_progenitors | IGLL5 |
| 2.26312152971362e-70 | 4.29647323355831 | 0.067 | 0.008 | 8.79607444953792e-66 | DC_progenitors | KCNQ3 |
| 3.73203254022886e-68 | 1.7222469166661 | 0.487 | 0.267 | 1.45052908741075e-63 | DC_progenitors | CCDC50 |
| 2.81308022595745e-67 | 1.59147928426523 | 0.411 | 0.195 | 1.09335989142288e-62 | DC_progenitors | SCPEP1 |
| 1.0011072567722e-65 | 2.09202854449802 | 0.2 | 0.06 | 3.89100357489652e-61 | DC_progenitors | SIGLEC6 |
| 0 | 5.14319155476957 | 0.707 | 0.141 | 0 | Neutrophil_like | AZU1 |
| 0 | 4.81706512158916 | 0.777 | 0.257 | 0 | Neutrophil_like | PRTN3 |
| 0 | 3.38826621437296 | 0.643 | 0.125 | 0 | Neutrophil_like | RNASE2 |
| 0 | 5.45448691495808 | 0.654 | 0.149 | 0 | Neutrophil_like | ELANE |
| 0 | 2.70206869864196 | 0.615 | 0.219 | 0 | Neutrophil_like | CFD |
| 0 | 2.89477108373619 | 0.527 | 0.142 | 0 | Neutrophil_like | CTSG |
| 0 | 2.61192186442162 | 0.636 | 0.269 | 0 | Neutrophil_like | CSF3R |
| 0 | 2.93989287313672 | 0.512 | 0.145 | 0 | Neutrophil_like | PRAM1 |
| 0 | 3.53558878683441 | 0.417 | 0.066 | 0 | Neutrophil_like | CSTA |
| 0 | 2.7526732160931 | 0.53 | 0.185 | 0 | Neutrophil_like | LYST |
| 0 | 2.4574998923555 | 0.511 | 0.181 | 0 | Neutrophil_like | CEBPD |

| | | | | | | |
|---|---|---|---|---|---|---|
| 0 | 2.79543348569953 | 0.523 | 0.206 | 0 | Neutrophil_like | FNDC3B |
| 0 | 1.61276643320941 | 0.69 | 0.373 | 0 | Neutrophil_like | PLAC8 |
| 0 | 3.56756327258712 | 0.977 | 0.683 | 0 | Neutrophil_like | MPO |
| 0 | 2.13100081951949 | 0.953 | 0.665 | 0 | Neutrophil_like | LYZ |
| 0 | 1.98031987732482 | 0.575 | 0.289 | 0 | Neutrophil_like | ZEB2 |
| 0 | 1.72328189675624 | 0.794 | 0.514 | 0 | Neutrophil_like | PRSS57 |
| 0 | 2.05679895988309 | 0.489 | 0.224 | 0 | Neutrophil_like | FAM107B |
| 0 | 1.9253581469875 | 0.481 | 0.218 | 0 | Neutrophil_like | NKG7 |
| 0 | 3.68662232266714 | 0.294 | 0.039 | 0 | Neutrophil_like | TRGC2 |
| 0 | 3.41871224595194 | 0.33 | 0.086 | 0 | Neutrophil_like | RETN |
| 0 | 1.69795076260676 | 0.518 | 0.292 | 0 | Neutrophil_like | TENT5A |
| 0 | 1.13181990884417 | 0.883 | 0.659 | 0 | Neutrophil_like | NUCB2 |
| 0 | 2.52978387925896 | 0.333 | 0.111 | 0 | Neutrophil_like | CLEC12A |
| 0 | 2.06066995478483 | 0.432 | 0.21 | 0 | Neutrophil_like | ERLIN1 |
| 0 | 1.63648540548641 | 0.402 | 0.182 | 0 | Neutrophil_like | MNDA |
| 0 | 1.55039029402693 | 0.454 | 0.236 | 0 | Neutrophil_like | CEBPA |
| 0 | 5.17597340043555 | 0.232 | 0.014 | 0 | Neutrophil_like | MS4A3 |
| 0 | 2.8679895744295 | 0.266 | 0.061 | 0 | Neutrophil_like | C16orf74 |
| 0 | 1.02808085870237 | 0.682 | 0.478 | 0 | Neutrophil_like | GIHCG |
| 0 | 2.4450921964254 | 0.346 | 0.142 | 0 | Neutrophil_like | EGR1 |
| 0 | 1.82293773923055 | 0.434 | 0.23 | 0 | Neutrophil_like | ENSG00000257764 |
| 0 | 4.64220027230745 | 0.218 | 0.015 | 0 | Neutrophil_like | SERPINB10 |
| 0 | 1.53529471932596 | 0.542 | 0.343 | 0 | Neutrophil_like | DENND10 |
| 0 | 1.79178251705411 | 0.415 | 0.216 | 0 | Neutrophil_like | SPNS3 |
| 0 | 1.47504757156293 | 0.577 | 0.378 | 0 | Neutrophil_like | CST7 |
| 0 | 1.81922213554988 | 0.498 | 0.301 | 0 | Neutrophil_like | MFSD10 |
| 0 | 2.3192965383 | 0.37 | 0.17 | 0 | Neutrophil_li | ATP8B4 |

| | | | | | | |
|---|---|---|---|---|---|---|
| | 5224 | 4 | 9 | | ke | |
| 0 | 1.13355388284283 | 0.77 | 0.576 | 0 | Neutrophil_like | ZFP36L2 |
| 0 | 3.08565813799701 | 0.254 | 0.063 | 0 | Neutrophil_like | SLC22A15 |
| 0 | 1.77038806427573 | 0.42 | 0.23 | 0 | Neutrophil_like | PXK |
| 0 | 1.84845046591472 | 0.337 | 0.149 | 0 | Neutrophil_like | SMIM3 |
| 0 | 3.16760458023853 | 0.221 | 0.037 | 0 | Neutrophil_like | TRGC1 |
| 0 | 1.7577550623294 | 0.348 | 0.165 | 0 | Neutrophil_like | GGT5 |
| 0 | 2.09145996258217 | 0.397 | 0.215 | 0 | Neutrophil_like | LRMDA |
| 0 | 3.23255375144953 | 0.227 | 0.048 | 0 | Neutrophil_like | LINC00926 |
| 0 | 1.90577998331274 | 0.434 | 0.255 | 0 | Neutrophil_like | MIR142HG |
| 0 | 1.18876772408427 | 0.588 | 0.41 | 0 | Neutrophil_like | TNFSF13B |
| 0 | 2.43822083802306 | 0.273 | 0.096 | 0 | Neutrophil_like | FUT4 |
| 0 | 1.55377463620682 | 0.815 | 0.64 | 0 | Neutrophil_like | SERPINB1 |
| 0 | 2.38168061690895 | 0.839 | 0.223 | 0 | GP_like | SPINK2 |
| 0 | 2.31595004882132 | 0.908 | 0.37 | 0 | GP_like | IGLL1 |
| 0 | 1.70169439413822 | 0.799 | 0.282 | 0 | GP_like | SMIM24 |
| 0 | 1.25460276773961 | 0.814 | 0.343 | 0 | GP_like | PLAC8 |
| 0 | 1.29331740989793 | 0.732 | 0.296 | 0 | GP_like | ITM2A |
| 0 | 1.60256529612129 | 0.597 | 0.189 | 0 | GP_like | CD34 |
| 0 | 1.22368303003032 | 0.708 | 0.301 | 0 | GP_like | RFLNB |
| 0 | 2.14010314866907 | 0.517 | 0.119 | 0 | GP_like | CLDN10 |
| 0 | 1.82240409072128 | 0.544 | 0.154 | 0 | GP_like | TRBC2 |
| 0 | 1.29234228584419 | 0.695 | 0.309 | 0 | GP_like | CDCA7 |
| 0 | 1.27055404554167 | 0.896 | 0.513 | 0 | GP_like | TYMS |
| 0 | 1.46425696940477 | 0.867 | 0.5 | 0 | GP_like | GYPC |
| 0 | 1.42120499430986 | 0.597 | 0.234 | 0 | GP_like | HMGN5 |
| 0 | 1.17832286249495 | 0.718 | 0.359 | 0 | GP_like | GGH |

| | | | | | | |
|---|---|---|---|---|---|---|
| 0 | 1.41363136210448 | 0.543 | 0.186 | 0 | GP_like | PTPRCAP |
| 0 | 1.16713477433395 | 0.788 | 0.433 | 0 | GP_like | BEX3 |
| 0 | 1.03282896712384 | 0.716 | 0.374 | 0 | GP_like | MCM2 |
| 0 | 1.23003289285785 | 0.692 | 0.35 | 0 | GP_like | DHFR |
| 0 | 1.56715457991548 | 0.593 | 0.251 | 0 | GP_like | MYC |
| 0 | 1.22764221819244 | 0.978 | 0.637 | 0 | GP_like | NUCB2 |
| 0 | 1.09653919982094 | 0.687 | 0.347 | 0 | GP_like | MAD2L1 |
| 0 | 1.24808667368041 | 0.46 | 0.123 | 0 | GP_like | HOPX |
| 0 | 1.02563429301721 | 0.783 | 0.447 | 0 | GP_like | LAPTM4B |
| 0 | 1.10598263810023 | 0.799 | 0.463 | 0 | GP_like | TM7SF3 |
| 0 | 1.29154027467318 | 0.881 | 0.551 | 0 | GP_like | XBP1 |
| 0 | 1.00767580197458 | 0.689 | 0.363 | 0 | GP_like | SYNGR1 |
| 0 | 1.10286187915004 | 0.516 | 0.196 | 0 | GP_like | CYTL1 |
| 0 | 1.03764649578461 | 0.675 | 0.356 | 0 | GP_like | SVIP |
| 0 | 1.53738052567367 | 0.483 | 0.167 | 0 | GP_like | PHGDH |
| 0 | 1.05277213269223 | 0.686 | 0.371 | 0 | GP_like | EBPL |
| 0 | 1.17156396030345 | 0.621 | 0.306 | 0 | GP_like | SNHG3 |
| 0 | 1.35379684335241 | 0.49 | 0.175 | 0 | GP_like | EGFL7 |
| 0 | 1.0611501736347 | 0.559 | 0.245 | 0 | GP_like | TCEAL9 |
| 0 | 1.04660792869216 | 0.74 | 0.426 | 0 | GP_like | LBR |
| 0 | 2.43049849064785 | 0.366 | 0.057 | 0 | GP_like | C1QTNF4 |
| 0 | 1.14767299874752 | 0.501 | 0.196 | 0 | GP_like | SPNS3 |
| 0 | 1.1361080126591 | 0.55 | 0.246 | 0 | GP_like | GINS2 |
| 0 | 1.04520790495731 | 0.711 | 0.41 | 0 | GP_like | MGST1 |
| 0 | 1.10793417294174 | 0.577 | 0.276 | 0 | GP_like | CENPH |
| 0 | 1.29150246321143 | 0.486 | 0.189 | 0 | GP_like | SNHG19 |
| 0 | 1.1014951409 | 0.57 | 0.28 | 0 | GP_like | DANCR |

| | | | | | | |
|---|---|---|---|---|---|---|
| | 4176 | 9 | 3 | | | |
| 0 | 1.00901516487742 | 0.628 | 0.332 | 0 | GP_like | GMNN |
| 0 | 1.78487499267308 | 0.383 | 0.091 | 0 | GP_like | MZB1 |
| 0 | 1.02987758981911 | 0.507 | 0.217 | 0 | GP_like | FAM30A |
| 0 | 1.04580074546426 | 0.672 | 0.384 | 0 | GP_like | NPM3 |
| 0 | 1.25838259911242 | 0.532 | 0.244 | 0 | GP_like | GCSH |
| 0 | 1.21130820206912 | 0.488 | 0.201 | 0 | GP_like | FAM216A |
| 0 | 1.07666199337423 | 0.493 | 0.212 | 0 | GP_like | CCDC34 |
| 0 | 2.19825532378557 | 0.359 | 0.078 | 0 | GP_like | ENSG00000224277 |
| 0 | 1.00871492059489 | 0.574 | 0.296 | 0 | GP_like | CMSS1 |
| 0 | 4.08230572974953 | 0.812 | 0.128 | 0 | HSPC | HOPX |
| 0 | 2.46938656167323 | 0.885 | 0.275 | 0 | HSPC | SPINK2 |
| 0 | 3.04243427867308 | 0.793 | 0.222 | 0 | HSPC | FAM30A |
| 0 | 2.5817124258848 | 0.719 | 0.247 | 0 | HSPC | HOXA9 |
| 0 | 2.15788054374298 | 0.671 | 0.22 | 0 | HSPC | CD34 |
| 0 | 1.70646098543082 | 0.777 | 0.33 | 0 | HSPC | SMIM24 |
| 0 | 2.448398539 | 0.654 | 0.215 | 0 | HSPC | CYTL1 |
| 0 | 4.63693132449368 | 0.494 | 0.057 | 0 | HSPC | MAL |
| 0 | 1.88336743370808 | 0.599 | 0.186 | 0 | HSPC | TRBC2 |
| 0 | 2.04640793409157 | 0.604 | 0.214 | 0 | HSPC | PTPRCAP |
| 0 | 3.65181389045558 | 0.441 | 0.053 | 0 | HSPC | IGHM |
| 0 | 2.85617914304512 | 0.513 | 0.126 | 0 | HSPC | HOXA7 |
| 0 | 5.36742121041718 | 0.403 | 0.024 | 0 | HSPC | KIAA0087 |
| 0 | 3.63521997562494 | 0.438 | 0.069 | 0 | HSPC | SNX10-AS1 |
| 0 | 1.53483679511053 | 0.759 | 0.393 | 0 | HSPC | OCIAD2 |
| 0 | 2.48624164710101 | 0.457 | 0.112 | 0 | HSPC | MZB1 |
| 0 | 1.38666072618557 | 0.684 | 0.339 | 0 | HSPC | ITM2A |

| | | | | | | |
|---|---|---|---|---|---|---|
| 0 | 1.38281699445877 | 0.802 | 0.476 | 0 | HSPC | LAPTM4B |
| 0 | 2.1932457340217 | 0.508 | 0.188 | 0 | HSPC | GNG11 |
| 0 | 1.84130192545172 | 0.503 | 0.183 | 0 | HSPC | ERG |
| 0 | 2.79147041874204 | 0.464 | 0.152 | 0 | HSPC | IL4R |
| 0 | 2.04693381849196 | 0.46 | 0.151 | 0 | HSPC | RAB27B |
| 0 | 4.10106717297711 | 0.391 | 0.084 | 0 | HSPC | JCHAIN |
| 0 | 1.49501226313751 | 0.764 | 0.46 | 0 | HSPC | PECAM1 |
| 0 | 2.19346891253844 | 0.473 | 0.169 | 0 | HSPC | SORL1 |
| 0 | 1.15781467087265 | 0.762 | 0.467 | 0 | HSPC | BEX3 |
| 0 | 1.70750432466605 | 0.467 | 0.173 | 0 | HSPC | PADI4 |
| 0 | 1.70592244503951 | 0.496 | 0.203 | 0 | HSPC | EGFL7 |
| 0 | 2.45039473356752 | 0.383 | 0.094 | 0 | HSPC | ADGRG1 |
| 0 | 2.28898387199859 | 0.398 | 0.117 | 0 | HSPC | SOCS2 |
| 0 | 1.77756225085546 | 0.427 | 0.161 | 0 | HSPC | CLDN10 |
| 0 | 2.3628033035873 | 0.346 | 0.085 | 0 | HSPC | ANGPT1 |
| 0 | 1.12995073135494 | 0.845 | 0.588 | 0 | HSPC | ITM2C |
| 0 | 2.39927580451562 | 0.338 | 0.082 | 0 | HSPC | BAALC |
| 0 | 2.31390526290125 | 0.349 | 0.093 | 0 | HSPC | BEX2 |
| 0 | 1.7891688528757 | 0.426 | 0.171 | 0 | HSPC | DUSP6 |
| 0 | 2.35695501190318 | 0.488 | 0.242 | 0 | HSPC | EPB41L2 |
| 0 | 2.17227126898482 | 0.327 | 0.088 | 0 | HSPC | C1QTNF4 |
| 0 | 4.27138799763078 | 0.26 | 0.03 | 0 | HSPC | ENSG00000286848 |
| 0 | 2.32092407705464 | 0.295 | 0.068 | 0 | HSPC | HEMGN |
| 0 | 2.19845068224983 | 0.343 | 0.118 | 0 | HSPC | IQCJ-SCHIP1 |
| 0 | 3.13595374703624 | 0.264 | 0.043 | 0 | HSPC | RBPMS |
| 0 | 4.04536856515067 | 0.244 | 0.024 | 0 | HSPC | MMP28 |
| 0 | 2.0984987192 | 0.30 | 0.08 | 0 | HSPC | F2R |

| | | | | | | |
|---|---|---|---|---|---|---|
| | 2378 | 3 | 6 | | | |
| 0 | 2.2116247491798 | 0.299 | 0.083 | 0 | HSPC | CD79B |
| 0 | 5.50542577096522 | 0.222 | 0.011 | 0 | HSPC | NPTX2 |
| 0 | 3.46353681880754 | 0.241 | 0.036 | 0 | HSPC | HOXA5 |
| 0 | 2.93385179654111 | 0.266 | 0.067 | 0 | HSPC | GZMA |
| 0 | 3.23990968001055 | 0.231 | 0.043 | 0 | HSPC | CRHBP |
| 0 | 3.31208935997617 | 0.216 | 0.036 | 0 | HSPC | F12 |
| 0 | 4.83887946326798 | 0.824 | 0.127 | 0 | MEP_like | GATA2 |
| 0 | 5.04775087620777 | 0.724 | 0.081 | 0 | MEP_like | HDC |
| 0 | 4.36411234949991 | 0.687 | 0.088 | 0 | MEP_like | SLC40A1 |
| 0 | 5.58913378049075 | 0.638 | 0.042 | 0 | MEP_like | CNRIP1 |
| 0 | 2.96179871239115 | 0.863 | 0.273 | 0 | MEP_like | CD82 |
| 0 | 4.47658672633246 | 0.673 | 0.088 | 0 | MEP_like | ITGA2B |
| 0 | 4.42196381316716 | 0.771 | 0.193 | 0 | MEP_like | HPGDS |
| 0 | 3.93517187274943 | 0.753 | 0.207 | 0 | MEP_like | CPA3 |
| 0 | 3.52005078838235 | 0.663 | 0.134 | 0 | MEP_like | TESPA1 |
| 0 | 3.13959290445247 | 0.809 | 0.329 | 0 | MEP_like | TFRC |
| 0 | 3.92051970162496 | 0.555 | 0.086 | 0 | MEP_like | CLU |
| 0 | 3.28215435613838 | 0.6 | 0.131 | 0 | MEP_like | ALDH1A1 |
| 0 | 3.41043726284387 | 0.571 | 0.108 | 0 | MEP_like | ST8SIA6 |
| 0 | 5.49334982275445 | 0.485 | 0.052 | 0 | MEP_like | KRT1 |
| 0 | 4.72998584226046 | 0.476 | 0.043 | 0 | MEP_like | ENSG00000261067 |
| 0 | 5.50343820032615 | 0.595 | 0.174 | 0 | MEP_like | HPGD |
| 0 | 1.99940601976299 | 0.579 | 0.158 | 0 | MEP_like | FCER1A |
| 0 | 5.53014333998978 | 0.438 | 0.023 | 0 | MEP_like | ENSG00000233968 |
| 0 | 4.96297358994847 | 0.466 | 0.061 | 0 | MEP_like | VWA5A |
| 0 | 2.31769809164926 | 0.605 | 0.212 | 0 | MEP_like | STXBP5 |

| | | | | | | |
|---|---|---|---|---|---|---|
| 0 | 4.40600370697343 | 0.426 | 0.047 | 0 | MEP_like | CAVIN2 |
| 0 | 5.02281139205971 | 0.396 | 0.028 | 0 | MEP_like | GRAP2 |
| 0 | 1.39832969324462 | 0.791 | 0.424 | 0 | MEP_like | GIHCG |
| 0 | 3.62476025155027 | 0.442 | 0.076 | 0 | MEP_like | ENSG00000251002 |
| 0 | 2.91004850823252 | 0.498 | 0.134 | 0 | MEP_like | TSC22D1 |
| 0 | 3.94251676486648 | 0.825 | 0.464 | 0 | MEP_like | LMO4 |
| 0 | 1.570403064 | 0.623 | 0.266 | 0 | MEP_like | MLC1 |
| 0 | 3.03564571283876 | 0.468 | 0.118 | 0 | MEP_like | PDLIM1 |
| 0 | 2.38309019078022 | 0.538 | 0.188 | 0 | MEP_like | CSF2RB |
| 0 | 2.96795139189562 | 0.433 | 0.091 | 0 | MEP_like | GCSAML |
| 0 | 3.69700814530296 | 0.411 | 0.069 | 0 | MEP_like | NMT2 |
| 0 | 1.53793787510514 | 0.725 | 0.384 | 0 | MEP_like | CDK6 |
| 0 | 1.64324053205683 | 0.787 | 0.459 | 0 | MEP_like | IQGAP2 |
| 0 | 4.75493956338052 | 0.345 | 0.02 | 0 | MEP_like | TIMP3 |
| 0 | 1.83429055908096 | 0.565 | 0.247 | 0 | MEP_like | PLIN2 |
| 0 | 4.86129943504858 | 0.333 | 0.016 | 0 | MEP_like | GATA1 |
| 0 | 1.57381709269085 | 0.579 | 0.263 | 0 | MEP_like | RAB37 |
| 0 | 4.92626121431433 | 0.338 | 0.023 | 0 | MEP_like | IL1RL1 |
| 0 | 3.50530289017179 | 0.362 | 0.048 | 0 | MEP_like | ZNF521 |
| 0 | 4.05781104186892 | 0.439 | 0.128 | 0 | MEP_like | RHEX |
| 0 | 3.36842874639437 | 0.359 | 0.052 | 0 | MEP_like | MTURN |
| 0 | 5.4522700244201 | 0.349 | 0.044 | 0 | MEP_like | TPSB2 |
| 0 | 1.63530542824541 | 0.565 | 0.263 | 0 | MEP_like | STING1 |
| 0 | 2.56383946525986 | 0.402 | 0.101 | 0 | MEP_like | EMID1 |
| 0 | 4.13764331899299 | 0.338 | 0.038 | 0 | MEP_like | SLC45A3 |
| 0 | 3.16137237010495 | 0.413 | 0.113 | 0 | MEP_like | ABCC4 |
| 0 | 1.4363009889 | 0.55 | 0.25 | 0 | MEP_like | MYB |

| | | | | | | |
|---|---|---|---|---|---|---|
| | 7109 | 7 | 8 | | | |
| 0 | 1.91517823152272 | 0.483 | 0.184 | 0 | MEP_like | PRKCQ-AS1 |
| 0 | 2.15344544093106 | 0.471 | 0.172 | 0 | MEP_like | ATP6V0A2 |
| 0 | 3.75420959801446 | 0.334 | 0.036 | 0 | MEP_like | TFR2 |

**top50_DEG_of_CBcDC1_subclusters**

| p_val | avg_log2FC | pct.1 | pct.2 | p_val_adj | cluster | gene |
|---|---|---|---|---|---|---|
| 0 | 2.17229736004564 | 0.81 | 0.23 | 0 | CBcDC1_C0 | AR |
| 0 | 2.0702007002236 | 0.736 | 0.223 | 0 | CBcDC1_C0 | SLC4A8 |
| 0 | 1.89862549887031 | 0.75 | 0.41 | 0 | CBcDC1_C0 | RNASE6 |
| 0 | 1.57297781027136 | 0.699 | 0.373 | 0 | CBcDC1_C0 | SERPINF1 |
| 0 | 1.79630339614545 | 0.883 | 0.577 | 0 | CBcDC1_C0 | LTB |
| 1.46704175447846e-245 | 2.04771218647616 | 0.46 | 0.171 | 5.70195118713145e-241 | CBcDC1_C0 | AQP3 |
| 4.49195172689165e-219 | 1.0857174524611 | 0.76 | 0.504 | 1.74588687769098e-214 | CBcDC1_C0 | TCEA3 |
| 7.08098575644681e-209 | 2.02312981194441 | 0.373 | 0.122 | 2.75216673395818e-204 | CBcDC1_C0 | LBH |
| 7.42573762001148e-194 | 1.15157394810018 | 0.677 | 0.415 | 2.88616144076986e-189 | CBcDC1_C0 | GPR171 |
| 2.93673711319238e-182 | 1.69572171106753 | 0.457 | 0.203 | 1.14142161378448e-177 | CBcDC1_C0 | TNF |
| 2.03534492177839e-151 | 1.21100191600523 | 0.547 | 0.31 | 7.91077510747605e-147 | CBcDC1_C0 | FFAR4 |
| 2.26740645372567e-134 | 1.45329154191124 | 0.375 | 0.168 | 8.81272866369558e-130 | CBcDC1_C0 | PTGER3 |
| 1.41792211459748e-122 | 1.5489039546165 | 0.305 | 0.121 | 5.51103788280601e-118 | CBcDC1_C0 | ENSG00000272980 |
| 2.7646122957203e-121 | 1.23505488959468 | 0.482 | 0.269 | 1.07452186097761e-116 | CBcDC1_C0 | FOSB |
| 7.35997205156275e-103 | 1.08621901945752 | 0.462 | 0.271 | 2.8606003372809e-98 | CBcDC1_C0 | NFAT5 |
| 4.62279848519026e-98 | 1.03494468272495 | 0.477 | 0.295 | 1.7967430872389e-93 | CBcDC1_C0 | PIK3CG |
| 8.91008463494634e-98 | 1.42735207499569 | 0.256 | 0.102 | 3.46308259506459e-93 | CBcDC1_C0 | NLRP1 |
| 9.71215871576079e-93 | 1.14344595291609 | 0.4 | 0.223 | 3.77482472805475e-88 | CBcDC1_C0 | TNFAIP8L2 |
| 9.14936623342771e-82 | 1.25986810341394 | 0.325 | 0.17 | 3.55608417394635e-77 | CBcDC1_C0 | SCN9A |
| 2.5387255404603e-81 | 1.95879022120788 | 0.273 | 0.129 | 9.86726455810703e-77 | CBcDC1_C0 | FCER1A |
| 2.84386102310458e-81 | 1.45620023262118 | 0.231 | 0.096 | 1.10532346385006e-76 | CBcDC1_C0 | PHETA1 |

| | | | | | | |
|---|---|---|---|---|---|---|
| 3.12041215039363e-81 | 1.01113470034817 | 0.441 | 0.275 | 1.21281059049349e-76 | CBcDC1_C0 | SLFN12L |
| 1.04846888415944e-76 | 1.07121798869288 | 0.397 | 0.237 | 4.07508401206248e-72 | CBcDC1_C0 | FILIP1L |
| 3.09988705321007e-68 | 1.13206313080556 | 0.401 | 0.251 | 1.20483310097116e-63 | CBcDC1_C0 | MAML2 |
| 2.26771884230616e-65 | 1.26247555739558 | 0.241 | 0.116 | 8.81394282439136e-61 | CBcDC1_C0 | NEIL1 |
| 3.37009475457809e-65 | 1.05016836348377 | 0.308 | 0.17 | 1.30985472826187e-60 | CBcDC1_C0 | RIPOR1 |
| 4.36131147253341e-65 | 1.05391409056252 | 0.329 | 0.189 | 1.69511093002956e-60 | CBcDC1_C0 | ZFP64 |
| 1.67040560379835e-64 | 1.21771727513767 | 0.252 | 0.125 | 6.49236546028306e-60 | CBcDC1_C0 | GNAO1 |
| 3.25012017687767e-64 | 1.08929409205561 | 0.285 | 0.15 | 1.26322420914704e-59 | CBcDC1_C0 | GPR82 |
| 4.34843187302008e-61 | 1.1603041452369 | 0.29 | 0.161 | 1.69010501608671e-56 | CBcDC1_C0 | SEPTIN3 |
| 1.06288539136258e-53 | 1.37509991281319 | 0.131 | 0.046 | 4.13111665060893e-49 | CBcDC1_C0 | CD300E |
| 1.93918944274109e-53 | 1.31947144370677 | 0.175 | 0.076 | 7.53704760710179e-49 | CBcDC1_C0 | HCAR1 |
| 3.64822302077567e-53 | 1.05667221921789 | 0.29 | 0.167 | 1.41795484148488e-48 | CBcDC1_C0 | ENSG00000285417 |
| 1.60770619948072e-51 | 1.13954686310245 | 0.237 | 0.126 | 6.2486716855217e-47 | CBcDC1_C0 | KLF8 |
| 1.64606759307047e-51 | 1.18344993748461 | 0.242 | 0.128 | 6.39777091398701e-47 | CBcDC1_C0 | EGR1 |
| 9.62471465626353e-51 | 1.12367832909941 | 0.225 | 0.116 | 3.74083784544995e-46 | CBcDC1_C0 | NCKAP5 |
| 2.94378851142237e-46 | 1.24148398957835 | 0.197 | 0.099 | 1.14416228073453e-41 | CBcDC1_C0 | TLR10 |
| 1.74209813604262e-45 | 1.62343338547383 | 0.129 | 0.052 | 6.77101282535683e-41 | CBcDC1_C0 | CD300C |
| 1.90566587847691e-45 | 1.57640310923641 | 0.174 | 0.084 | 7.40675156987619e-41 | CBcDC1_C0 | SNORD13 |
| 2.40275284162176e-45 | 1.39684329253337 | 0.169 | 0.08 | 9.33877946953129e-41 | CBcDC1_C0 | CLDN23 |
| 1.60394573560061e-44 | 1.0818437561558 | 0.218 | 0.118 | 6.2340558905589e-40 | CBcDC1_C0 | PNPLA6 |
| 3.47780974290496e-43 | 1.08088452918053 | 0.178 | 0.087 | 1.35172031277487e-38 | CBcDC1_C0 | TMEM150A |
| 1.34748425783138e-42 | 1.08259157575689 | 0.211 | 0.114 | 5.23726706491321e-38 | CBcDC1_C0 | CALHM2 |
| 1.39291288866221e-42 | 1.55247896192243 | 0.128 | 0.052 | 5.41383452436342e-38 | CBcDC1_C0 | ENSG00000258875 |
| 1.60050952293256e-42 | 1.01220034392551 | 0.208 | 0.112 | 6.22070036278198e-38 | CBcDC1_C0 | PIGZ |
| 2.1663863024037e-42 | 1.19780589753078 | 0.219 | 0.121 | 8.42009364155246e-38 | CBcDC1_C0 | TPM1 |
| 2.6149005376336e-41 | 1.22693709631601 | 0.144 | 0.064 | 1.01633339196205e-36 | CBcDC1_C0 | TMEM273 |
| 8.68744660578647e | 1.31479196236 | 0.12 | 0.05 | 3.37654987227103e | CBcDC1_ | SLAMF1 |

| | | | | | | |
|---|---|---|---|---|---|---|
| -41 | 985 | 3 | | -36 | C0 | |
| 1.17253705042561e-40 | 1.17564947717048 | 0.158 | 0.075 | 4.55729975388923e-36 | CBcDC1_C0 | SEL1L3 |
| 6.43481547427913e-37 | 1.29681069115803 | 0.138 | 0.064 | 2.50101973038807e-32 | CBcDC1_C0 | PLAAT3 |
| 0 | 2.32403135921049 | 0.704 | 0.181 | 0 | CBcDC1_C1 | MKI67 |
| 0 | 1.92193315758591 | 0.792 | 0.302 | 0 | CBcDC1_C1 | SMC4 |
| 0 | 1.95822358255579 | 0.746 | 0.257 | 0 | CBcDC1_C1 | TOP2A |
| 0 | 2.07991904235607 | 0.676 | 0.197 | 0 | CBcDC1_C1 | CCNB2 |
| 0 | 1.83939418002677 | 0.754 | 0.281 | 0 | CBcDC1_C1 | TACC3 |
| 0 | 2.03525644276378 | 0.628 | 0.157 | 0 | CBcDC1_C1 | ZWINT |
| 0 | 1.70466607981917 | 0.677 | 0.209 | 0 | CBcDC1_C1 | MAD2L1 |
| 0 | 1.53119675595526 | 0.863 | 0.397 | 0 | CBcDC1_C1 | PCLAF |
| 0 | 1.46019489853624 | 0.872 | 0.415 | 0 | CBcDC1_C1 | CKS1B |
| 0 | 2.23330357340971 | 0.625 | 0.173 | 0 | CBcDC1_C1 | FEN1 |
| 0 | 2.02781132925822 | 0.621 | 0.175 | 0 | CBcDC1_C1 | PRC1 |
| 0 | 2.33572749949553 | 0.579 | 0.136 | 0 | CBcDC1_C1 | CDKN3 |
| 0 | 1.52565440812712 | 0.706 | 0.266 | 0 | CBcDC1_C1 | CENPW |
| 0 | 1.91299075483457 | 0.618 | 0.18 | 0 | CBcDC1_C1 | CDK1 |
| 0 | 1.94149808344609 | 0.622 | 0.184 | 0 | CBcDC1_C1 | RAD51AP1 |
| 0 | 2.28393011118833 | 0.592 | 0.154 | 0 | CBcDC1_C1 | CDC20 |
| 0 | 1.9302216519009 | 0.602 | 0.164 | 0 | CBcDC1_C1 | CENPU |
| 0 | 2.07287490660214 | 0.571 | 0.134 | 0 | CBcDC1_C1 | CEP55 |
| 0 | 1.94579078500878 | 0.6 | 0.165 | 0 | CBcDC1_C1 | TPX2 |
| 0 | 1.74237478958239 | 0.642 | 0.216 | 0 | CBcDC1_C1 | ENSG00000285920 |
| 0 | 1.89574443595713 | 0.635 | 0.209 | 0 | CBcDC1_C1 | UBE2C |
| 0 | 2.1764516101139 | 0.553 | 0.128 | 0 | CBcDC1_C1 | NCAPG |
| 0 | 1.42463249208621 | 0.769 | 0.344 | 0 | CBcDC1_C1 | TMPO |
| 0 | 1.93714205404374 | 0.586 | 0.162 | 0 | CBcDC1_C1 | AURKB |

| | | | | | | |
|---|---|---|---|---|---|---|
| 0 | 2.09091413413452 | 0.543 | 0.125 | 0 | CBcDC1_C1 | CCNA2 |
| 0 | 2.61090392562363 | 0.561 | 0.145 | 0 | CBcDC1_C1 | CCNB1 |
| 0 | 1.96558869804922 | 0.58 | 0.164 | 0 | CBcDC1_C1 | RRM2 |
| 0 | 1.7224176228958 | 0.642 | 0.232 | 0 | CBcDC1_C1 | TYMS |
| 0 | 1.9330918613183 | 0.554 | 0.147 | 0 | CBcDC1_C1 | BIRC5 |
| 0 | 1.46952667854319 | 0.607 | 0.202 | 0 | CBcDC1_C1 | GGH |
| 0 | 2.28780489285204 | 0.522 | 0.119 | 0 | CBcDC1_C1 | PLK1 |
| 0 | 1.89304101956399 | 0.581 | 0.18 | 0 | CBcDC1_C1 | GMNN |
| 0 | 1.6226248311496 | 0.597 | 0.197 | 0 | CBcDC1_C1 | KIFC1 |
| 0 | 1.83368433828937 | 0.577 | 0.179 | 0 | CBcDC1_C1 | ORC6 |
| 0 | 2.21504091004668 | 0.526 | 0.132 | 0 | CBcDC1_C1 | CENPF |
| 0 | 1.93228736684326 | 0.51 | 0.116 | 0 | CBcDC1_C1 | ASF1B |
| 0 | 1.61240559915526 | 0.587 | 0.194 | 0 | CBcDC1_C1 | DTYMK |
| 0 | 1.5688234072638 | 0.664 | 0.274 | 0 | CBcDC1_C1 | UBE2S |
| 0 | 1.74276862808112 | 0.513 | 0.137 | 0 | CBcDC1_C1 | CENPM |
| 0 | 2.02243351766466 | 0.48 | 0.107 | 0 | CBcDC1_C1 | NUF2 |
| 0 | 1.64875096840009 | 0.896 | 0.527 | 0 | CBcDC1_C1 | TUBB4B |
| 0 | 2.10208583577071 | 0.495 | 0.127 | 0 | CBcDC1_C1 | HMMR |
| 0 | 2.02844022223974 | 0.468 | 0.11 | 0 | CBcDC1_C1 | KIF11 |
| 0 | 2.41652565306448 | 0.447 | 0.092 | 0 | CBcDC1_C1 | SGO2 |
| 0 | 2.23728133221519 | 0.439 | 0.087 | 0 | CBcDC1_C1 | NDC80 |
| 0 | 2.16082035602619 | 0.445 | 0.108 | 0 | CBcDC1_C1 | CENPE |
| 0 | 2.3950002904096 | 0.413 | 0.079 | 0 | CBcDC1_C1 | CDCA3 |
| 0 | 2.67057480914901 | 0.39 | 0.066 | 0 | CBcDC1_C1 | CENPA |
| 0 | 1.41142109518197 | 0.98 | 0.656 | 0 | CBcDC1_C1 | STMN1 |
| 0 | 2.53503005950485 | 0.394 | 0.072 | 0 | CBcDC1_C1 | ASPM |
| 0 | 1.54202208995 | 0.89 | 0.33 | 0 | CBcDC1_ | EPB41L3 |

| | | | | | | |
|---|---|---|---|---|---|---|
| | 719 | | 7 | | C2 | |
| 0 | 1.91189075533456 | 0.725 | 0.328 | 0 | CBcDC1_C2 | DDIT4 |
| 2.13546937763339e-287 | 1.12756219755699 | 0.893 | 0.851 | 8.29992883004769e-283 | CBcDC1_C2 | MCL1 |
| 2.21001198674322e-252 | 2.03781119890536 | 0.617 | 0.29 | 8.58965358887486e-248 | CBcDC1_C2 | SLC2A3 |
| 5.25097805906362e-199 | 2.0992673003802 | 0.55 | 0.272 | 2.04089764221626e-194 | CBcDC1_C2 | ANKRD37 |
| 3.25983944469264e-165 | 1.11962453304352 | 0.785 | 0.725 | 1.26700179696869e-160 | CBcDC1_C2 | RAB7B |
| 7.47692636900868e-147 | 1.81471089481499 | 0.576 | 0.353 | 2.9060569718426e-142 | CBcDC1_C2 | RGS2 |
| 1.82294438252729e-144 | 1.52424954432499 | 0.619 | 0.45 | 7.08523793156881e-140 | CBcDC1_C2 | VDR |
| 9.53147820425528e-140 | 1.90275652332328 | 0.523 | 0.31 | 3.7045996336479e-135 | CBcDC1_C2 | DUSP4 |
| 1.20022470641957e-122 | 1.41809740097906 | 0.61 | 0.449 | 4.66491336644094e-118 | CBcDC1_C2 | TYMP |
| 2.0586727233106e-115 | 1.47336185358507 | 0.494 | 0.271 | 8.00144327369132e-111 | CBcDC1_C2 | HILPDA |
| 4.73946364432924e-113 | 2.6768421902381 | 0.227 | 0.07 | 1.84208733464145e-108 | CBcDC1_C2 | NRARP |
| 5.69746136109538e-97 | 1.13588055193942 | 0.66 | 0.584 | 2.21443230721694e-92 | CBcDC1_C2 | IVNS1ABP |
| 2.18709518159244e-96 | 1.12373021666238 | 0.643 | 0.533 | 8.50058284229532e-92 | CBcDC1_C2 | CYTIP |
| 7.27102375433582e-87 | 1.89217320907076 | 0.463 | 0.297 | 2.8260288025977e-82 | CBcDC1_C2 | TACSTD2 |
| 1.23919466598889e-84 | 1.46984382175449 | 0.435 | 0.263 | 4.81637790829902e-80 | CBcDC1_C2 | SOCS1 |
| 1.65954079368486e-82 | 1.2396811345289 | 0.579 | 0.47 | 6.45013720281493e-78 | CBcDC1_C2 | ST8SIA4 |
| 9.17797074213595e-82 | 1.74458553469882 | 0.324 | 0.16 | 3.56720188834598e-77 | CBcDC1_C2 | TENT5C |
| 3.10328604743144e-78 | 1.37631662596375 | 0.458 | 0.296 | 1.20615418805518e-73 | CBcDC1_C2 | SLC2A1 |
| 6.90533724459991e-74 | 1.63775638034551 | 0.383 | 0.225 | 2.68389742685865e-69 | CBcDC1_C2 | BCL2L11 |
| 1.27835455968145e-69 | 1.16720584818631 | 0.526 | 0.373 | 4.96858066711391e-65 | CBcDC1_C2 | C1QA |
| 3.33388054499576e-66 | 1.14065359120994 | 0.546 | 0.427 | 1.2957793514235e-61 | CBcDC1_C2 | CHD6 |
| 3.99692464831837e-65 | 1.76824201577063 | 0.358 | 0.214 | 1.5534847030619e-60 | CBcDC1_C2 | SGK1 |
| 1.11909434292146e-64 | 2.55444221996896 | 0.114 | 0.029 | 4.34958398263283e-60 | CBcDC1_C2 | TAMALIN |
| 6.50803027367205e-56 | 1.24395898811569 | 0.541 | 0.445 | 2.52947612646811e-51 | CBcDC1_C2 | CD83 |
| 3.15552306717951e-55 | 2.02157521687485 | 0.188 | 0.08 | 1.22645715052066e-50 | CBcDC1_C2 | NR4A1 |
| 2.88758089153847e-54 | 1.12424350535972 | 0.51 | 0.401 | 1.12231606511426e-49 | CBcDC1_C2 | HMOX1 |

| | | | | | | |
|---|---|---|---|---|---|---|
| 1.45805974951389e-53 | 1.4161246007223 | 0.309 | 0.176 | 5.66704082843564e-49 | CBcDC1_C2 | RNF144B |
| 7.83203605447591e-52 | 1.51390383843823 | 0.262 | 0.137 | 3.04407745329315e-47 | CBcDC1_C2 | PIM2 |
| 8.12251077525848e-52 | 1.6269320146413 | 0.325 | 0.197 | 3.15697626301971e-47 | CBcDC1_C2 | ENSG00000198211 |
| 5.69762186588106e-51 | 1.0582619949083 | 0.564 | 0.486 | 2.21449469061199e-46 | CBcDC1_C2 | APOL1 |
| 2.73514813229806e-49 | 1.71823620898764 | 0.363 | 0.238 | 1.06307002458029e-44 | CBcDC1_C2 | HSPA1B |
| 3.45433952532566e-47 | 1.20878862149967 | 0.473 | 0.371 | 1.34259814330832e-42 | CBcDC1_C2 | PLAUR |
| 3.86841427390534e-45 | 1.62485251093447 | 0.207 | 0.102 | 1.50353657583879e-40 | CBcDC1_C2 | DUSP5 |
| 1.7050538444992e-44 | 1.29653372241548 | 0.404 | 0.292 | 6.62703277741505e-40 | CBcDC1_C2 | PFKFB3 |
| 5.62391617100277e-43 | 1.93154762241774 | 0.171 | 0.077 | 2.18584749818365e-38 | CBcDC1_C2 | ID3 |
| 7.64276982886453e-43 | 1.73749489686601 | 0.155 | 0.066 | 2.97051534938478e-38 | CBcDC1_C2 | SEMA7A |
| 3.1480131758237e-42 | 1.2815978642962 | 0.467 | 0.369 | 1.2235382810474e-37 | CBcDC1_C2 | ATF5 |
| 1.10468178748111e-40 | 1.6033161547729 | 0.225 | 0.122 | 4.29356670340281e-36 | CBcDC1_C2 | PER1 |
| 2.37062323920562e-40 | 1.80714536605082 | 0.24 | 0.137 | 9.21390134382049e-36 | CBcDC1_C2 | DUSP2 |
| 4.76073269576475e-40 | 1.1535780524504 | 0.32 | 0.204 | 1.85035397686288e-35 | CBcDC1_C2 | HK2 |
| 1.55109862019188e-39 | 1.10769445800934 | 0.438 | 0.33 | 6.02865500709977e-35 | CBcDC1_C2 | CDKN1A |
| 6.60621931908026e-38 | 1.36914506305811 | 0.268 | 0.16 | 2.56763926274693e-33 | CBcDC1_C2 | RAPGEF2 |
| 8.15424747834487e-38 | 2.32548410790991 | 0.082 | 0.025 | 3.1693113674083e-33 | CBcDC1_C2 | NR4A3 |
| 2.72782184334614e-37 | 1.84906747091146 | 0.114 | 0.044 | 1.06022251585334e-32 | CBcDC1_C2 | ARID5B |
| 8.51787594364963e-36 | 1.15663028312005 | 0.387 | 0.287 | 3.3106428430183e-31 | CBcDC1_C2 | N4BP2L1 |
| 1.87569280921118e-34 | 1.02228294055914 | 0.475 | 0.387 | 7.29025524156109e-30 | CBcDC1_C2 | P4HA1 |
| 6.73951806005309e-34 | 1.52551745018166 | 0.214 | 0.121 | 2.61944848440084e-29 | CBcDC1_C2 | FOSL2 |
| 9.77044106676962e-34 | 1.2134794658364 | 0.356 | 0.252 | 3.79747732942135e-29 | CBcDC1_C2 | IDO1 |
| 8.53152074517056e-32 | 1.07129468530511 | 0.387 | 0.292 | 3.31594616802544e-27 | CBcDC1_C2 | GPR146 |
| 8.2636087047318e-300 | 1.13891904139819 | 0.979 | 0.89 | 3.21181679526811e-295 | CBcDC1_C3 | TUBB |
| 3.54789467064428e-260 | 1.14701885339002 | 0.977 | 0.741 | 1.37896022163931e-255 | CBcDC1_C3 | HMGB2 |
| 3.08340309712652e-246 | 1.19807595371066 | 0.907 | 0.391 | 1.19842628176016e-241 | CBcDC1_C3 | EPB41L3 |
| 7.15915290758174e | 1.47019335106 | 0.82 | 0.48 | 2.78254796058979e | CBcDC1_ | CKS1B |

| | | | | | | |
|---|---|---|---|---|---|---|
| -232 | 035 | 9 | 1 | -227 | C3 | |
| 2.98472064224083e-231 | 1.1153693849585 | 0.953 | 0.702 | 1.16007137201974e-226 | CBcDC1_C3 | STMN1 |
| 1.34549378773465e-228 | 1.26889335851758 | 0.91 | 0.626 | 5.22953070478828e-224 | CBcDC1_C3 | H4C3 |
| 1.45103225794578e-193 | 1.26949672298349 | 0.755 | 0.364 | 5.63972707695787e-189 | CBcDC1_C3 | DDIT4 |
| 8.23184374567568e-185 | 1.63535992773982 | 0.627 | 0.266 | 3.19947070863177e-180 | CBcDC1_C3 | UBE2C |
| 6.05600919185792e-179 | 1.46009967752197 | 0.695 | 0.329 | 2.35378909259942e-174 | CBcDC1_C3 | TOP2A |
| 6.62745127504566e-172 | 1.65307732962304 | 0.588 | 0.242 | 2.575891487072e-167 | CBcDC1_C3 | CDK1 |
| 5.19799986261836e-157 | 1.22394831346239 | 0.779 | 0.471 | 2.02030660660388e-152 | CBcDC1_C3 | PCLAF |
| 2.13869863646699e-130 | 1.21147359992857 | 0.776 | 0.541 | 8.31247999035627e-126 | CBcDC1_C3 | PCNA |
| 1.39746609592214e-123 | 1.54361015896273 | 0.515 | 0.229 | 5.43153147502058e-119 | CBcDC1_C3 | RRM2 |
| 2.84498524240332e-110 | 1.1568687055641 | 0.563 | 0.283 | 1.1057604141649e-105 | CBcDC1_C3 | HILPDA |
| 3.16240199063022e-110 | 1.42305566777569 | 0.504 | 0.23 | 1.22913078169825e-105 | CBcDC1_C3 | AURKB |
| 3.88163273810885e-104 | 3.17198733780175 | 0.116 | 0.014 | 1.50867419632077e-99 | CBcDC1_C3 | ENSG00000224647 |
| 3.80194301873174e-103 | 1.34962709304476 | 0.553 | 0.289 | 1.47770119309046e-98 | CBcDC1_C3 | MAD2L1 |
| 8.75392047257834e-92 | 1.01023550472552 | 0.767 | 0.624 | 3.40238627007702e-87 | CBcDC1_C3 | DUT |
| 1.13261514042755e-80 | 1.61884018690311 | 0.399 | 0.188 | 4.40213526629976e-76 | CBcDC1_C3 | ENSG00000288637 |
| 2.24967850136257e-80 | 1.2396571396595 | 0.525 | 0.29 | 8.74382543124589e-76 | CBcDC1_C3 | ENSG00000285920 |
| 2.56349926378725e-79 | 1.0943791886421 | 0.619 | 0.395 | 9.96355258856189e-75 | CBcDC1_C3 | MCM7 |
| 3.54744496896283e-74 | 1.27629350027902 | 0.432 | 0.213 | 1.37878543608678e-69 | CBcDC1_C3 | CEP55 |
| 5.38811739748135e-69 | 1.56472025605158 | 0.344 | 0.155 | 2.09419958887908e-64 | CBcDC1_C3 | KIF2C |
| 3.17386663025185e-66 | 1.24437318378697 | 0.509 | 0.305 | 1.23358674317999e-61 | CBcDC1_C3 | KIF22 |
| 5.02256298056697e-66 | 1.39392434217115 | 0.415 | 0.215 | 1.95211955365696e-61 | CBcDC1_C3 | UBE2T |
| 1.38463486010615e-64 | 1.0225853763306 | 0.501 | 0.286 | 5.38166031077459e-60 | CBcDC1_C3 | CCNB2 |
| 7.6363421579798e-63 | 1.1136528596018 | 0.511 | 0.305 | 2.96801710654201e-58 | CBcDC1_C3 | TYMS |
| 1.61023275158379e-62 | 1.09093156287213 | 0.559 | 0.368 | 6.25849163558073e-58 | CBcDC1_C3 | PHF19 |
| 2.53271316319913e-61 | 1.46721990344228 | 0.502 | 0.319 | 9.84389625140605e-57 | CBcDC1_C3 | LINC01588 |
| 9.22180407343549e-61 | 1.03127315641889 | 0.439 | 0.235 | 3.58423858922217e-56 | CBcDC1_C3 | CDC20 |

| | | | | | | |
|---|---|---|---|---|---|---|
| 1.89801112187404e-60 | 1.47433813545419 | 0.343 | 0.163 | 7.37699982738784e-56 | CBcDC1_C3 | BUB1 |
| 4.09481793792801e-60 | 1.0073835995168 | 0.23 | 0.083 | 1.59153288793448e-55 | CBcDC1_C3 | ADM |
| 5.14110918449769e-60 | 1.21054041263266 | 0.445 | 0.247 | 1.99819490673872e-55 | CBcDC1_C3 | ZWINT |
| 4.42453129410849e-58 | 1.27679822202138 | 0.426 | 0.241 | 1.71968257808115e-53 | CBcDC1_C3 | HSPA1B |
| 2.20267681937876e-57 | 1.40256572312535 | 0.364 | 0.185 | 8.56114399387944e-53 | CBcDC1_C3 | TK1 |
| 6.13243252451476e-57 | 1.44549909236471 | 0.335 | 0.162 | 2.38349254930315e-52 | CBcDC1_C3 | SPAG5 |
| 8.12486068464439e-57 | 2.45034399603824 | 0.071 | 0.01 | 3.15788960230074e-52 | CBcDC1_C3 | TNFSF9 |
| 1.13452746315242e-55 | 1.13409116543064 | 0.381 | 0.193 | 4.40956789103449e-51 | CBcDC1_C3 | PLK1 |
| 3.12669553615324e-55 | 1.2468807943655 | 0.388 | 0.203 | 1.21525275403668e-50 | CBcDC1_C3 | CCNA2 |
| 9.26373048806515e-55 | 1.28737680861616 | 0.386 | 0.205 | 3.60053412879628e-50 | CBcDC1_C3 | CENPM |
| 1.03134806753469e-54 | 1.11805112523582 | 0.522 | 0.343 | 4.00854053408706e-50 | CBcDC1_C3 | NCAPD2 |
| 4.64669181502019e-54 | 1.0931406360949 | 0.439 | 0.246 | 1.8060297077439e-49 | CBcDC1_C3 | TPX2 |
| 6.67866913605359e-54 | 1.05481649836308 | 0.528 | 0.345 | 2.59579833310995e-49 | CBcDC1_C3 | UBE2S |
| 4.07488720294811e-53 | 2.4013114115283 | 0.141 | 0.042 | 1.58378640916984e-48 | CBcDC1_C3 | GSG1 |
| 2.35077096195421e-52 | 1.39230822386077 | 0.348 | 0.176 | 9.13674149782742e-48 | CBcDC1_C3 | NUF2 |
| 1.43296879362392e-49 | 1.61555795593836 | 0.158 | 0.052 | 5.56951981017808e-45 | CBcDC1_C3 | HSPA6 |
| 1.10995230085039e-48 | 1.4121524345643 | 0.306 | 0.151 | 4.31405160771522e-44 | CBcDC1_C3 | CDCA5 |
| 1.28720574203515e-48 | 2.39878006678658 | 0.123 | 0.035 | 5.002982557568e-44 | CBcDC1_C3 | MROH8 |
| 2.72638791517722e-46 | 1.22174679450478 | 0.373 | 0.208 | 1.05966519099193e-41 | CBcDC1_C3 | SHCBP1 |
| 8.46427314907371e-46 | 1.13427807136198 | 0.434 | 0.264 | 3.28980904485048e-41 | CBcDC1_C3 | CENPN |
| 0 | 3.05079675477395 | 0.673 | 0.149 | 0 | CBcDC1_C4 | RNF130 |
| 0 | 3.94824692572954 | 0.378 | 0.05 | 0 | CBcDC1_C4 | PLAC8 |
| 3.77921751406288e-301 | 2.97096596054932 | 0.512 | 0.107 | 1.46886847119082e-296 | CBcDC1_C4 | SAMSN1 |
| 5.37293805574614e-291 | 3.26505300145128 | 0.374 | 0.051 | 2.08829983412685e-286 | CBcDC1_C4 | ZEB2 |
| 4.0280014869211e-287 | 3.28763663469189 | 0.672 | 0.242 | 1.56556333792162e-282 | CBcDC1_C4 | NUCB2 |
| 5.83903174688157e-279 | 4.09643257262768 | 0.329 | 0.041 | 2.26945646906046e-274 | CBcDC1_C4 | RNASE2 |
| 4.01149069574372e | 3.45327504649 | 0.48 | 0.12 | 1.55914608871471e | CBcDC1_ | PRSS57 |

| | | | | | | |
|---|---|---|---|---|---|---|
| -250 | 993 | 3 | 1 | -245 | C4 | |
| 9.52944410581271e-241 | 2.1346956937173 | 0.971 | 0.85 | 3.70380904060623e-236 | CBcDC1_C4 | SRGN |
| 4.10800462630704e-203 | 3.06267975034881 | 0.324 | 0.057 | 1.59665815810676e-198 | CBcDC1_C4 | NFE2 |
| 3.93176060312219e-202 | 2.50384617570532 | 0.355 | 0.069 | 1.5281573936155e-197 | CBcDC1_C4 | CHST11 |
| 2.01213873903568e-186 | 3.0062001252624 | 0.566 | 0.225 | 7.82057963700998e-182 | CBcDC1_C4 | MRC1 |
| 5.09079757197502e-185 | 3.98717573731573 | 0.239 | 0.032 | 1.97864029229953e-180 | CBcDC1_C4 | CTSG |
| 6.4569204222068e-174 | 2.9106202219838 | 0.316 | 0.063 | 2.50961126049912e-169 | CBcDC1_C4 | NKG7 |
| 1.91787131815233e-171 | 2.57019652532475 | 0.355 | 0.081 | 7.45419045226266e-167 | CBcDC1_C4 | GBP1 |
| 4.09332394374116e-171 | 3.38896545522472 | 0.218 | 0.028 | 1.59095221721388e-166 | CBcDC1_C4 | MLC1 |
| 2.46708328487239e-169 | 3.11608523450699 | 0.213 | 0.027 | 9.5888126033135e-165 | CBcDC1_C4 | LAIR1 |
| 2.79547802982136e-168 | 2.93456589673296 | 0.265 | 0.044 | 1.08651844585067e-163 | CBcDC1_C4 | STING1 |
| 9.33897759115412e-165 | 3.92251521471167 | 0.455 | 0.152 | 3.62978042035387e-160 | CBcDC1_C4 | AZU1 |
| 4.15021557859102e-162 | 2.40114006234875 | 0.295 | 0.057 | 1.61306428893097e-157 | CBcDC1_C4 | RAB37 |
| 1.90800352569132e-161 | 3.41844228490468 | 0.845 | 0.664 | 7.41583730330445e-157 | CBcDC1_C4 | MPO |
| 2.30162190336673e-159 | 3.37834131546483 | 0.21 | 0.028 | 8.94571385181545e-155 | CBcDC1_C4 | PRAM1 |
| 1.93146282262386e-157 | 2.39483540186205 | 0.361 | 0.089 | 7.50701655269214e-153 | CBcDC1_C4 | LAMP1 |
| 4.24017817365582e-157 | 2.60204437399454 | 0.434 | 0.14 | 1.64803005075481e-152 | CBcDC1_C4 | GIHCG |
| 5.98070911013229e-154 | 2.75436819720302 | 0.252 | 0.043 | 2.32452220983512e-149 | CBcDC1_C4 | TFEC |
| 1.15710260652099e-153 | 2.10302307444988 | 0.553 | 0.23 | 4.49731070076513e-149 | CBcDC1_C4 | ZFP36L2 |
| 1.15744773748418e-152 | 3.07511141071468 | 0.199 | 0.026 | 4.49865212127975e-148 | CBcDC1_C4 | P2RX1 |
| 4.52815450784268e-147 | 3.4599967268856 | 0.208 | 0.031 | 1.75995781256321e-142 | CBcDC1_C4 | MAOA |
| 6.7613741802608e-146 | 1.82321017911931 | 0.589 | 0.262 | 2.62794330264197e-141 | CBcDC1_C4 | HCST |
| 1.44348781822387e-144 | 4.06782221740984 | 0.565 | 0.265 | 5.61040410309073e-140 | CBcDC1_C4 | PRTN3 |
| 3.26123469048568e-144 | 2.52567381313161 | 0.26 | 0.05 | 1.26754408715107e-139 | CBcDC1_C4 | SIRPA |
| 1.44685257862648e-143 | 3.43612724095511 | 0.242 | 0.044 | 5.62348191734753e-139 | CBcDC1_C4 | ITM2A |
| 1.05835316069986e-141 | 3.10601395055852 | 0.18 | 0.023 | 4.11350122969216e-137 | CBcDC1_C4 | SMIM3 |
| 4.67684414298388e-141 | 3.81925262330396 | 0.243 | 0.046 | 1.81774901305354e-136 | CBcDC1_C4 | CPA3 |

| | | | | | | |
|---|---|---|---|---|---|---|
| 3.82195845187669e-140 | 2.61521425656208 | 0.236 | 0.042 | 1.48548059149091e-135 | CBcDC1_C4 | CSF1R |
| 8.63329690922569e-139 | 1.82278370676855 | 0.568 | 0.246 | 3.35550350970875e-134 | CBcDC1_C4 | GLUL |
| 1.39858923345481e-136 | 4.51706467686181 | 0.431 | 0.156 | 5.43589677366881e-132 | CBcDC1_C4 | ELANE |
| 7.0211327667824e-134 | 2.39216735013301 | 0.339 | 0.091 | 2.72890367246532e-129 | CBcDC1_C4 | LYST |
| 3.17580923675469e-132 | 1.26151058899188 | 0.869 | 0.692 | 1.23434177604944e-127 | CBcDC1_C4 | MT-TL1 |
| 2.19690427680123e-130 | 2.67601456336888 | 0.199 | 0.031 | 8.53870785264334e-126 | CBcDC1_C4 | TBXAS1 |
| 3.25592633272475e-130 | 2.28009604894659 | 0.363 | 0.106 | 1.26548088774013e-125 | CBcDC1_C4 | TESC |
| 7.75942672447953e-130 | 3.0419768057531 | 0.239 | 0.047 | 3.01585638500346e-125 | CBcDC1_C4 | CSTA |
| 1.29337735463592e-127 | 2.68915051201712 | 0.298 | 0.074 | 5.02696976426344e-123 | CBcDC1_C4 | CEBPA |
| 1.54684767734699e-127 | 2.96170170502094 | 0.181 | 0.026 | 6.01213286754453e-123 | CBcDC1_C4 | PTAFR |
| 1.41341157229027e-125 | 3.02915121278509 | 0.278 | 0.067 | 5.4935067580206e-121 | CBcDC1_C4 | CD82 |
| 1.58745742337469e-125 | 2.36344788332092 | 0.291 | 0.071 | 6.16997076743042e-121 | CBcDC1_C4 | GGT5 |
| 5.24500311032533e-124 | 2.85157637040172 | 0.18 | 0.027 | 2.03857535889015e-119 | CBcDC1_C4 | CD300LF |
| 6.47917855842403e-122 | 3.80225880762135 | 0.125 | 0.011 | 2.51826233030267e-117 | CBcDC1_C4 | TRGC2 |
| 1.16974942943102e-121 | 3.05159203331082 | 0.15 | 0.018 | 4.54646510736955e-117 | CBcDC1_C4 | CYBB |
| 2.43990015107436e-119 | 2.76818679763091 | 0.208 | 0.038 | 9.48315991718072e-115 | CBcDC1_C4 | RRAGD |
| 4.87661226666914e-118 | 3.72531171931311 | 0.12 | 0.011 | 1.89539288968629e-113 | CBcDC1_C4 | CCR1 |

**top50_DEG_of_CBcDC2_subclusters**

| p_val | avg_log2FC | pct.1 | pct.2 | p_val_adj | cluster | gene |
|---|---|---|---|---|---|---|
| 0 | 3.5524500392585 | 0.718 | 0.074 | 0 | CBcDC2_C0 | AR |
| 0 | 3.38017980003545 | 0.649 | 0.076 | 0 | CBcDC2_C0 | SLC4A8 |
| 0 | 1.09914158783021 | 0.921 | 0.68 | 0 | CBcDC2_C0 | ALOX5AP |
| 1.11992111469459e-282 | 1.41558676781918 | 0.869 | 0.567 | 4.35279739648345e-278 | CBcDC2_C0 | LTB |
| 2.60581949945411e-272 | 1.58529654298297 | 0.832 | 0.526 | 1.01280386485283e-267 | CBcDC2_C0 | FOS |
| 3.42287184847268e-234 | 1.7999951125841 | 0.599 | 0.24 | 1.33036760134588e-229 | CBcDC2_C0 | AQP3 |
| 3.64556135739805e-202 | 1.52967499128408 | 0.584 | 0.238 | 1.4169203327799e-197 | CBcDC2_C0 | ALOX5 |
| 1.434832697423e- | 1.49586280546 | 0.62 | 0.32 | 5.57676424507398e | CBcDC2_ | RCBTB2 |

| | | | | | | |
|---|---|---|---|---|---|---|
| 186 | 044 | 8 | | -182 | C0 | |
| 1.95495127061997e-186 | 1.09936536897553 | 0.723 | 0.381 | 7.59830910351862e-182 | CBcDC2_C0 | ENSG00000272980 |
| 2.83632456393996e-180 | 1.07248252472549 | 0.777 | 0.5 | 1.10239426826654e-175 | CBcDC2_C0 | BASP1 |
| 4.20461995086687e-171 | 1.49685506542192 | 0.616 | 0.306 | 1.63420963630343e-166 | CBcDC2_C0 | IER3 |
| 5.64654732923699e-158 | 1.14048744577396 | 0.714 | 0.426 | 2.19464355045454e-153 | CBcDC2_C0 | RNF24 |
| 8.65575654473046e-151 | 1.0464446622926 | 0.71 | 0.439 | 3.36423289624039e-146 | CBcDC2_C0 | PON2 |
| 3.65087838513445e-148 | 1.22800501506541 | 0.563 | 0.267 | 1.4189869019502e-143 | CBcDC2_C0 | SERPINF1 |
| 1.02858627113208e-144 | 1.59999785355344 | 0.499 | 0.218 | 3.99780626000906e-140 | CBcDC2_C0 | FCER1A |
| 6.92306790076678e-136 | 1.62562825764285 | 0.448 | 0.183 | 2.69078880099102e-131 | CBcDC2_C0 | FOSB |
| 5.36228992167813e-134 | 2.124113633 | 0.338 | 0.1 | 2.08416122385864e-129 | CBcDC2_C0 | EGR1 |
| 3.24752734966428e-128 | 1.7106605686357 | 0.34 | 0.105 | 1.26221645499402e-123 | CBcDC2_C0 | PTGER3 |
| 2.49270117299808e-121 | 1.32178286031325 | 0.504 | 0.248 | 9.68838164909164e-117 | CBcDC2_C0 | GPR171 |
| 1.40320820684069e-120 | 1.49891811618767 | 0.383 | 0.142 | 5.4538493375277e-116 | CBcDC2_C0 | PLBD1 |
| 1.77362591726382e-120 | 1.38918482234221 | 0.428 | 0.181 | 6.8935518526293e-116 | CBcDC2_C0 | P2RY6 |
| 2.41313635595021e-120 | 1.03299847368578 | 0.578 | 0.307 | 9.37913707467169e-116 | CBcDC2_C0 | NAPSB |
| 2.38033234013085e-119 | 1.39331069988537 | 0.421 | 0.175 | 9.25163770638657e-115 | CBcDC2_C0 | SCN9A |
| 3.4518385715067e-115 | 1.69075115948696 | 0.345 | 0.121 | 1.34162609758751e-110 | CBcDC2_C0 | EGR2 |
| 5.534620462493e-113 | 1.05177699351757 | 0.519 | 0.261 | 2.15114093515715e-108 | CBcDC2_C0 | P2RY14 |
| 8.40501077454048e-108 | 1.76627406920818 | 0.322 | 0.112 | 3.26677553774065e-103 | CBcDC2_C0 | TCEA3 |
| 1.87140641172102e-103 | 1.12772526721244 | 0.556 | 0.317 | 7.2735953004361e-99 | CBcDC2_C0 | CD69 |
| 1.94390470226415e-99 | 1.00520493037698 | 0.532 | 0.293 | 7.55537440629009e-95 | CBcDC2_C0 | OPN3 |
| 6.33631325114825e-98 | 1.1086981436964 | 0.471 | 0.24 | 2.46273487132379e-93 | CBcDC2_C0 | NDRG2 |
| 1.15207015026069e-97 | 1.15840917166964 | 0.444 | 0.219 | 4.47775105301821e-93 | CBcDC2_C0 | CBX6 |
| 2.24911988872086e-97 | 1.04476865370982 | 0.456 | 0.225 | 8.74165427149136e-93 | CBcDC2_C0 | ENSG00000232628 |
| 2.28155849388726e-97 | 1.04907796579266 | 0.487 | 0.258 | 8.8677333981916e-93 | CBcDC2_C0 | SYS1-DBNDD2 |
| 1.96931157372821e-96 | 1.08908676025636 | 0.493 | 0.262 | 7.65412329360943e-92 | CBcDC2_C0 | HLA-DOB |
| 9.07222947383622e-92 | 2.38543169282205 | 0.196 | 0.042 | 3.52610342959592e-87 | CBcDC2_C0 | ENC1 |

| | | | | | | |
|---|---|---|---|---|---|---|
| 9.47711461014151e-92 | 1.19481597740013 | 0.423 | 0.206 | 3.6834701355237e-87 | CBcDC2_C0 | PIM1 |
| 3.29529327378671e-81 | 1.95433167911697 | 0.212 | 0.058 | 1.28078163672268e-76 | CBcDC2_C0 | CD2 |
| 7.35858627845815e-76 | 1.4264058846327 | 0.271 | 0.104 | 2.86006172884833e-71 | CBcDC2_C0 | RFXAP |
| 6.53560355408375e-74 | 1.30267016093682 | 0.298 | 0.125 | 2.54019303336573e-69 | CBcDC2_C0 | MT-TT |
| 1.80107858907861e-71 | 1.96624200665467 | 0.209 | 0.065 | 7.00025215217184e-67 | CBcDC2_C0 | DUSP6 |
| 5.43951795287986e-70 | 1.11425687275097 | 0.332 | 0.156 | 2.11417744274582e-65 | CBcDC2_C0 | MTFMT |
| 6.50055331767495e-69 | 1.04403433261729 | 0.329 | 0.151 | 2.52657005798072e-64 | CBcDC2_C0 | FILIP1L |
| 6.02329692421332e-66 | 1.24731546728312 | 0.345 | 0.176 | 2.34107481553399e-61 | CBcDC2_C0 | MSI2 |
| 8.52898977671257e-62 | 1.29861896577539 | 0.263 | 0.114 | 3.31496245651488e-57 | CBcDC2_C0 | PRMT7 |
| 1.65377870108531e-60 | 1.41796177359692 | 0.223 | 0.084 | 6.42774167750827e-56 | CBcDC2_C0 | DPP4 |
| 9.85633195677332e-59 | 1.65655804677011 | 0.179 | 0.057 | 3.83086054163908e-54 | CBcDC2_C0 | ENSG00000258875 |
| 3.95606893052473e-58 | 1.10558145751093 | 0.321 | 0.162 | 1.53760531122705e-53 | CBcDC2_C0 | FPR1 |
| 1.69982579112239e-57 | 1.73236565883589 | 0.161 | 0.047 | 6.60671290235539e-53 | CBcDC2_C0 | ADAMTS14 |
| 5.13693183565611e-57 | 1.55911746681539 | 0.196 | 0.07 | 1.99657129656446e-52 | CBcDC2_C0 | NEIL1 |
| 9.94603136803115e-57 | 1.16726870586568 | 0.246 | 0.104 | 3.86572401181267e-52 | CBcDC2_C0 | DEPTOR |
| 7.46070118928495e-55 | 2.50365781230853 | 0.103 | 0.016 | 2.89975073123938e-50 | CBcDC2_C0 | ENSG00000261222 |
| 0 | 2.8660406809065 | 0.878 | 0.201 | 0 | CBcDC2_C1 | EPB41L3 |
| 0 | 3.30286554653125 | 0.619 | 0.151 | 0 | CBcDC2_C1 | HILPDA |
| 0 | 2.87508983445447 | 0.718 | 0.251 | 0 | CBcDC2_C1 | DDIT4 |
| 0 | 3.84837700520161 | 0.424 | 0.045 | 0 | CBcDC2_C1 | ADM |
| 0 | 2.64402734572752 | 0.894 | 0.568 | 0 | CBcDC2_C1 | SLC2A3 |
| 2.9120529711643e-264 | 1.26172296610741 | 0.982 | 0.972 | 1.13182762830243e-259 | CBcDC2_C1 | TXN |
| 4.6424647712514e-226 | 1.62876595014992 | 0.75 | 0.537 | 1.80438678264228e-221 | CBcDC2_C1 | STAT1 |
| 8.41860762396161e-215 | 1.04591379191679 | 0.868 | 0.795 | 3.27206022520516e-210 | CBcDC2_C1 | PSME2 |
| 1.01140887321708e-198 | 2.44794532406849 | 0.514 | 0.221 | 3.93104286753282e-194 | CBcDC2_C1 | ANKRD37 |
| 1.13606198788642e-197 | 1.51398230000828 | 0.725 | 0.548 | 4.41553212831816e-193 | CBcDC2_C1 | TYMP |
| 1.04859324461694e | 1.74180159471 | 0.67 | 0.40 | 4.07556736385267e | CBcDC2_ | C1QB |

| | | | | | | |
|---|---|---|---|---|---|---|
| -183 | 56 | 3 | 4 | -179 | C1 | |
| 1.32043933111021e-179 | 1.54943199033893 | 0.689 | 0.427 | 5.13215154822604e-175 | CBcDC2_C1 | C1QA |
| 2.26340037457832e-179 | 1.43506036618289 | 0.813 | 0.713 | 8.79715823587355e-175 | CBcDC2_C1 | FABP5 |
| 4.22708144475346e-176 | 1.38154965645475 | 0.721 | 0.568 | 1.64293974513233e-171 | CBcDC2_C1 | LAP3 |
| 1.01454568349123e-159 | 1.5395031834945 | 0.663 | 0.481 | 3.94323470802535e-155 | CBcDC2_C1 | WARS1 |
| 8.18625795040867e-147 | 2.19441739461645 | 0.469 | 0.219 | 3.18175287758534e-142 | CBcDC2_C1 | NAMPT |
| 4.03848024043315e-139 | 2.04730442330934 | 0.494 | 0.262 | 1.56963611504915e-134 | CBcDC2_C1 | SLC2A1 |
| 1.40720394885762e-135 | 1.16325758752072 | 0.853 | 0.837 | 5.46937958802492e-131 | CBcDC2_C1 | CTSB |
| 6.3428154032536e-134 | 2.6204062217784 | 0.335 | 0.109 | 2.46526206278258e-129 | CBcDC2_C1 | FOSL2 |
| 6.84428000930442e-134 | 2.24607914672481 | 0.397 | 0.158 | 2.66016631121635e-129 | CBcDC2_C1 | BIRC3 |
| 7.66845583510422e-133 | 2.88544540621178 | 0.341 | 0.114 | 2.98049872942996e-128 | CBcDC2_C1 | HSPA1B |
| 1.52397677548463e-129 | 2.66273466841218 | 0.298 | 0.085 | 5.9232405332761e-125 | CBcDC2_C1 | NR4A3 |
| 1.60976722734545e-127 | 1.86469660698663 | 0.426 | 0.185 | 6.25668228252358e-123 | CBcDC2_C1 | CXCR4 |
| 1.07322288851293e-125 | 1.60471443669754 | 0.684 | 0.558 | 4.17129540078319e-121 | CBcDC2_C1 | CD83 |
| 2.91391244144692e-118 | 1.83048789774518 | 0.43 | 0.198 | 1.13255034861717e-113 | CBcDC2_C1 | ITGAX |
| 1.2366522511259e-114 | 2.09368374352516 | 0.476 | 0.265 | 4.80649630445102e-110 | CBcDC2_C1 | RGS2 |
| 3.61589698409464e-108 | 1.1123487075165 | 0.736 | 0.652 | 1.40539068080806e-103 | CBcDC2_C1 | PLEK |
| 1.20649132610845e-107 | 2.27025823867944 | 0.334 | 0.13 | 4.68926983718572e-103 | CBcDC2_C1 | SNAPC1 |
| 4.05600280074566e-107 | 2.72390578237725 | 0.362 | 0.158 | 1.57644660856582e-102 | CBcDC2_C1 | SPP1 |
| 4.09698036810353e-105 | 2.3053872902256 | 0.319 | 0.12 | 1.5923733596708e-100 | CBcDC2_C1 | MXD1 |
| 7.79794267152699e-102 | 1.17905127518293 | 0.643 | 0.521 | 3.03082637814239e-97 | CBcDC2_C1 | CYTIP |
| 2.97961898898655e-101 | 1.80623026534389 | 0.412 | 0.204 | 1.1580885124494e-96 | CBcDC2_C1 | GBP2 |
| 6.69203221044824e-101 | 1.44773633997999 | 0.535 | 0.342 | 2.60099215923492e-96 | CBcDC2_C1 | GBP1 |
| 1.36951710158115e-99 | 2.16558587566105 | 0.359 | 0.163 | 5.32290211871544e-95 | CBcDC2_C1 | PER1 |
| 2.93351859893607e-93 | 1.9035321860545 | 0.387 | 0.188 | 1.14017067384848e-88 | CBcDC2_C1 | C15orf48 |
| 1.63201400181052e-90 | 1.90490709634163 | 0.352 | 0.162 | 6.34314882083695e-86 | CBcDC2_C1 | AVPI1 |
| 1.09902475916221e-88 | 2.21802739017147 | 0.2 | 0.048 | 4.27157953143575e-84 | CBcDC2_C1 | ARRDC3 |

| | | | | | | |
|---|---|---|---|---|---|---|
| 5.50786560325554e-84 | 1.30496682627548 | 0.572 | 0.437 | 2.14074212401733e-79 | CBcDC2_C1 | PDXK |
| 1.84622334464227e-80 | 3.19191496180217 | 0.127 | 0.016 | 7.17571627362111e-76 | CBcDC2_C1 | IFI27 |
| 6.68920851667612e-77 | 1.06002671010026 | 0.649 | 0.556 | 2.59989467417651e-72 | CBcDC2_C1 | PIM3 |
| 2.2276639918791e-76 | 1.82287941278376 | 0.313 | 0.14 | 8.65826163723651e-72 | CBcDC2_C1 | IL4I1 |
| 3.2553375432464e-76 | 1.63246145255096 | 0.372 | 0.192 | 1.26525204293358e-71 | CBcDC2_C1 | NFKBID |
| 5.60195167298296e-70 | 2.31488396084022 | 0.327 | 0.164 | 2.17731055673829e-65 | CBcDC2_C1 | LINC01588 |
| 1.29776642537855e-68 | 1.54154646232809 | 0.369 | 0.202 | 5.04402876551883e-64 | CBcDC2_C1 | GPR146 |
| 2.6789266939606e-68 | 1.87377482639865 | 0.255 | 0.102 | 1.04121843814167e-63 | CBcDC2_C1 | SERPING1 |
| 8.78425764143953e-68 | 2.43512251159186 | 0.179 | 0.052 | 3.4141774174983e-63 | CBcDC2_C1 | SLAMF7 |
| 8.62230681690914e-67 | 1.88212984050877 | 0.267 | 0.115 | 3.35123199052808e-62 | CBcDC2_C1 | CD80 |
| 1.54525023572242e-66 | 1.6547851275536 | 0.347 | 0.183 | 6.00592409118233e-62 | CBcDC2_C1 | IL18BP |
| 5.16621968774129e-66 | 2.65134415540736 | 0.152 | 0.037 | 2.00795460603441e-61 | CBcDC2_C1 | MREG |
| 1.09799776222913e-65 | 1.4024148947541 | 0.469 | 0.324 | 4.26758790245596e-61 | CBcDC2_C1 | HK2 |
| 1.22127562032071e-214 | 3.03289915223187 | 0.528 | 0.102 | 4.7467319535005e-210 | CBcDC2_C2 | MCM4 |
| 2.8477374062833e-191 | 2.79545787877299 | 0.588 | 0.15 | 1.10683009770013e-186 | CBcDC2_C2 | TYMS |
| 8.9955601788229e-183 | 2.84578155334779 | 0.456 | 0.086 | 3.4963043747031e-178 | CBcDC2_C2 | MCM2 |
| 9.57926889257186e-182 | 3.22950138117408 | 0.328 | 0.039 | 3.72317444047591e-177 | CBcDC2_C2 | GINS2 |
| 9.97571659455743e-172 | 1.90396412205878 | 0.862 | 0.519 | 3.87726176880664e-167 | CBcDC2_C2 | STMN1 |
| 2.20512172879336e-171 | 3.17909012611545 | 0.531 | 0.136 | 8.57064662330115e-167 | CBcDC2_C2 | PRSS57 |
| 1.3590625387944e-166 | 3.96651930348991 | 0.284 | 0.031 | 5.28226836953218e-162 | CBcDC2_C2 | CTSG |
| 2.6495028356469e-164 | 2.66005251439154 | 0.372 | 0.059 | 1.02978226713088e-159 | CBcDC2_C2 | CDCA7 |
| 4.25802506005685e-159 | 2.28624987628383 | 0.625 | 0.197 | 1.6549666000923e-154 | CBcDC2_C2 | PCLAF |
| 9.48319969678664e-157 | 2.91233392649885 | 0.422 | 0.085 | 3.68583522615006e-152 | CBcDC2_C2 | FEN1 |
| 1.12051440479685e-156 | 2.60738666822753 | 0.704 | 0.307 | 4.35510333712393e-152 | CBcDC2_C2 | PCNA |
| 3.31512915865309e-156 | 2.43279713450428 | 0.55 | 0.152 | 1.2884912500937e-151 | CBcDC2_C2 | MCM7 |
| 5.56791748620584e-155 | 2.42517723301714 | 0.55 | 0.154 | 2.16408248936362e-150 | CBcDC2_C2 | MCM3 |
| 2.62065562417532e | 2.10402214501 | 0.75 | 0.36 | 1.01857022144822e | CBcDC2_ | NUCB2 |

| | | | | | | |
|---|---|---|---|---|---|---|
| -142 | 207 | 9 | 8 | -137 | C2 | |
| 7.53219484771899e-142 | 2.92639702844289 | 0.375 | 0.072 | 2.92753817146294e-137 | CBcDC2_C2 | PLAC8 |
| 4.09662800830812e-138 | 2.69518286439933 | 0.313 | 0.049 | 1.59223640798912e-133 | CBcDC2_C2 | CDT1 |
| 8.03709281886506e-137 | 3.15891496318568 | 0.251 | 0.03 | 3.12377686590828e-132 | CBcDC2_C2 | CLSPN |
| 9.51397493979441e-136 | 1.24014071757678 | 0.956 | 0.808 | 3.69779663984989e-131 | CBcDC2_C2 | HMGN2 |
| 1.29559886312717e-128 | 2.85288845229654 | 0.321 | 0.057 | 5.03560410131639e-124 | CBcDC2_C2 | CENPU |
| 3.6419126975142e-125 | 3.45319799863269 | 0.204 | 0.021 | 1.41550220814284e-120 | CBcDC2_C2 | CDC6 |
| 1.73321276230848e-118 | 2.51993660880644 | 0.335 | 0.067 | 6.73647804326438e-114 | CBcDC2_C2 | ZWINT |
| 3.45429930742327e-113 | 1.38846839299085 | 0.888 | 0.718 | 1.3425825118162e-108 | CBcDC2_C2 | H2AZ1 |
| 1.00315503306549e-111 | 2.1886113672477 | 0.368 | 0.084 | 3.89896266701565e-107 | CBcDC2_C2 | GGH |
| 4.51601435116787e-110 | 2.50590444467404 | 0.416 | 0.114 | 1.75523929786842e-105 | CBcDC2_C2 | GIHCG |
| 1.08908419506793e-106 | 4.55962057842789 | 0.14 | 0.01 | 4.23294354097054e-102 | CBcDC2_C2 | FAM111B |
| 4.01420261939393e-105 | 1.09345817721328 | 0.934 | 0.789 | 1.56020013207984e-100 | CBcDC2_C2 | HMGB1 |
| 3.13573805715887e-104 | 2.10551524822074 | 0.481 | 0.157 | 1.21876731067594e-99 | CBcDC2_C2 | MCM6 |
| 7.95017699214869e-103 | 3.16899918975599 | 0.221 | 0.032 | 3.08999529153843e-98 | CBcDC2_C2 | TCF19 |
| 3.08525194706853e-102 | 2.68961419949528 | 0.207 | 0.028 | 1.19914487426713e-97 | CBcDC2_C2 | HELLS |
| 5.67332927696281e-100 | 3.05727613292382 | 0.846 | 0.646 | 2.20505289007714e-95 | CBcDC2_C2 | MPO |
| 1.91145705922144e-94 | 2.77597116251809 | 0.213 | 0.032 | 7.42926015207595e-90 | CBcDC2_C2 | ASF1B |
| 2.04584745149926e-94 | 2.92129902893707 | 0.187 | 0.024 | 7.95159528974217e-90 | CBcDC2_C2 | DTL |
| 2.69925252021593e-92 | 1.49643439240385 | 0.766 | 0.477 | 1.04911847703232e-87 | CBcDC2_C2 | DUT |
| 4.87737523653165e-90 | 1.36015452619462 | 0.835 | 0.579 | 1.89568943318276e-85 | CBcDC2_C2 | HMGB2 |
| 5.19068669661586e-90 | 3.21806837691101 | 0.163 | 0.019 | 2.01746419837368e-85 | CBcDC2_C2 | E2F1 |
| 5.85651948558203e-89 | 2.68129055490097 | 0.232 | 0.042 | 2.27625342846117e-84 | CBcDC2_C2 | UNG |
| 1.18298067103939e-86 | 3.44734458201726 | 0.128 | 0.011 | 4.59789097412881e-82 | CBcDC2_C2 | ZNF367 |
| 1.17761660341526e-85 | 2.65397071812953 | 0.206 | 0.033 | 4.5770424524941e-81 | CBcDC2_C2 | MYB |
| 5.84602398404378e-85 | 2.37875776382736 | 0.331 | 0.087 | 2.2721741418783e-80 | CBcDC2_C2 | RNASE2 |
| 1.3382454909694e-84 | 1.80980653941806 | 0.503 | 0.2 | 5.20135874975077e-80 | CBcDC2_C2 | DNAJC9 |

| | | | | | | |
|---|---|---|---|---|---|---|
| 1.43751657619497e-84 | 2.57069965215776 | 0.24 | 0.046 | 5.587195676697e-80 | CBcDC2_C2 | MLC1 |
| 7.76745948951971e-84 | 3.3875990758437 | 0.141 | 0.015 | 3.01897847979162e-79 | CBcDC2_C2 | CCNE2 |
| 3.14309760271661e-82 | 2.75390335416811 | 0.178 | 0.026 | 1.22162774524787e-77 | CBcDC2_C2 | MYBL2 |
| 1.35100894854526e-80 | 3.10204126934164 | 0.162 | 0.021 | 5.25096648031087e-76 | CBcDC2_C2 | WDR76 |
| 2.06140794184443e-80 | 1.76778445015679 | 0.375 | 0.111 | 8.01207424756677e-76 | CBcDC2_C2 | CFAP44 |
| 2.77955182371129e-80 | 2.36953717492878 | 0.278 | 0.065 | 1.08032840732187e-75 | CBcDC2_C2 | ORC6 |
| 3.33620629512523e-79 | 3.30951855641301 | 0.124 | 0.011 | 1.29668330072632e-74 | CBcDC2_C2 | MCM10 |
| 6.78018240439265e-79 | 2.30460239542266 | 0.215 | 0.039 | 2.63525349511529e-74 | CBcDC2_C2 | CENPM |
| 5.04212035577888e-77 | 2.31027490636607 | 0.288 | 0.073 | 1.95972091868058e-72 | CBcDC2_C2 | MKI67 |
| 1.91027696874581e-76 | 1.95345924065594 | 0.381 | 0.123 | 7.42467349442433e-72 | CBcDC2_C2 | MSH6 |
| 0 | 8.02261207360508 | 0.579 | 0.004 | 0 | CBcDC2_C3 | MIR4432HG |
| 0 | 10.2079843464996 | 0.395 | 0 | 0 | CBcDC2_C3 | ENSG00000213557 |
| 9.84626682117445e-233 | 7.86093891477435 | 0.342 | 0.002 | 3.82694852538588e-228 | CBcDC2_C3 | FCRLA |
| 3.83238965272622e-223 | 7.55581270024776 | 0.553 | 0.008 | 1.4895348863251e-218 | CBcDC2_C3 | GPR153 |
| 3.43691151332025e-171 | 7.93439297893347 | 0.211 | 0.001 | 1.33582439788218e-166 | CBcDC2_C3 | LINC01724 |
| 3.61295376065224e-165 | 5.95709033832207 | 0.816 | 0.029 | 1.40424673815271e-160 | CBcDC2_C3 | CX3CR1 |
| 9.25854788440133e-164 | 5.63292503277354 | 0.474 | 0.009 | 3.59851980623026e-159 | CBcDC2_C3 | MSC-AS1 |
| 3.01461275720762e-142 | 6.32949025909045 | 0.263 | 0.003 | 1.17168954034389e-137 | CBcDC2_C3 | BDKRB2 |
| 5.73407870764222e-142 | 5.71501543395832 | 0.474 | 0.01 | 2.2286643712993e-137 | CBcDC2_C3 | EPHB1 |
| 1.95488012867837e-138 | 5.99804169381499 | 0.579 | 0.017 | 7.59803259613423e-134 | CBcDC2_C3 | ENSG00000286848 |
| 8.55282805081791e-127 | 6.35644470301868 | 0.316 | 0.005 | 3.3242276785114e-122 | CBcDC2_C3 | AXL |
| 7.93229256557338e-115 | 5.78312433048023 | 0.921 | 0.059 | 3.0830441514614e-110 | CBcDC2_C3 | JCHAIN |
| 2.0397444149225e-92 | 5.67721833727357 | 0.526 | 0.021 | 7.92787461747928e-88 | CBcDC2_C3 | TSPAN13 |
| 1.25720684155909e-78 | 5.21951159319223 | 0.658 | 0.042 | 4.88638583108772e-74 | CBcDC2_C3 | ENSG00000285417 |
| 1.95988582342958e-78 | 6.20564160771946 | 0.184 | 0.003 | 7.61748822992375e-74 | CBcDC2_C3 | SPEG |
| 1.31244340132914e-77 | 5.30632318379829 | 0.553 | 0.029 | 5.10107376794597e-73 | CBcDC2_C3 | SIGLEC6 |
| 1.62727087702516e | 5.11146424453 | 0.23 | 0.00 | 6.32471371773369e | CBcDC2_ | KCNK10 |

| | | | | | | |
|---|---|---|---|---|---|---|
| -74 | 452 | 7 | 5 | -70 | C3 | |
| 7.29137536751045e-74 | 4.83878711407238 | 0.395 | 0.015 | 2.83393886409029e-69 | CBcDC2_C3 | ECE1 |
| 4.20950566859087e-72 | 7.7875452905068 | 0.105 | 0.001 | 1.63610856821121e-67 | CBcDC2_C3 | KDR |
| 4.75784928421551e-72 | 5.38126999809613 | 0.289 | 0.008 | 1.84923328129604e-67 | CBcDC2_C3 | CAMP |
| 1.07153391300435e-69 | 5.7490906420396 | 0.184 | 0.003 | 4.164730859674e-65 | CBcDC2_C3 | RGS13 |
| 3.64641078356637e-65 | 4.28960256576613 | 0.395 | 0.017 | 1.41725047924874e-60 | CBcDC2_C3 | PROC |
| 1.29859666873963e-63 | 4.66466904785363 | 0.447 | 0.023 | 5.04725567239031e-59 | CBcDC2_C3 | STEAP4 |
| 2.23389088098629e-59 | 4.98170785138161 | 0.421 | 0.022 | 8.68246368712941e-55 | CBcDC2_C3 | SCAMP5 |
| 2.32948541247512e-59 | 7.11747922167497 | 0.132 | 0.002 | 9.05401095266703e-55 | CBcDC2_C3 | GZMB |
| 4.61580472021081e-59 | 6.96939563274821 | 0.105 | 0.001 | 1.79402482060433e-54 | CBcDC2_C3 | CH25H |
| 4.61885456400138e-59 | 6.33865749339289 | 0.211 | 0.005 | 1.79521020339042e-54 | CBcDC2_C3 | EBF4 |
| 6.09459943739926e-59 | 5.28905899332552 | 0.342 | 0.014 | 2.36878796333397e-54 | CBcDC2_C3 | EPHA2 |
| 2.08863060083131e-57 | 3.937009092 | 0.579 | 0.042 | 8.11788055625104e-53 | CBcDC2_C3 | SPIB |
| 1.39624596088897e-54 | 5.04278756783048 | 0.316 | 0.013 | 5.42678917618717e-50 | CBcDC2_C3 | SCD5 |
| 1.80196948090482e-54 | 4.40184589933406 | 0.684 | 0.067 | 7.00371478143278e-50 | CBcDC2_C3 | IGLON5 |
| 1.60731213512015e-51 | 5.04818440568882 | 0.211 | 0.006 | 6.24714007557148e-47 | CBcDC2_C3 | CLEC4C |
| 2.66821372915716e-46 | 4.5937324068644 | 0.158 | 0.004 | 1.03705463011151e-41 | CBcDC2_C3 | CCDC68 |
| 4.21733258956957e-46 | 4.14967324461847 | 0.579 | 0.054 | 1.639150657588e-41 | CBcDC2_C3 | TOGARAM2 |
| 2.11304511293659e-44 | 4.87571742526799 | 0.289 | 0.014 | 8.21277244045066e-40 | CBcDC2_C3 | NCAM1 |
| 2.29367280543392e-43 | 4.13168209185322 | 0.921 | 0.177 | 8.91481809288002e-39 | CBcDC2_C3 | TCF4 |
| 2.92301395272533e-43 | 5.52395835591457 | 0.105 | 0.002 | 1.13608783300575e-38 | CBcDC2_C3 | HHIP-AS1 |
| 5.79294236885957e-43 | 3.80236692429608 | 0.632 | 0.07 | 2.25154291050465e-38 | CBcDC2_C3 | DUSP4 |
| 9.01229137132296e-43 | 5.0713073141828 | 0.237 | 0.009 | 3.5028072872921e-38 | CBcDC2_C3 | CXCR2 |
| 8.12412798611906e-42 | 8.51153048009065 | 0.026 | 0 | 3.15760482436489e-37 | CBcDC2_C3 | FAM43B |
| 8.12412798611906e-42 | 8.89954367988263 | 0.026 | 0 | 3.15760482436489e-37 | CBcDC2_C3 | ENSG00000273487 |
| 8.12412798611906e-42 | 8.26733528629218 | 0.026 | 0 | 3.15760482436489e-37 | CBcDC2_C3 | E2F3P2 |
| 8.12412798611906e-42 | 8.1441466159563 | 0.026 | 0 | 3.15760482436489e-37 | CBcDC2_C3 | SCYL2P1 |

| | | | | | | |
|---|---|---|---|---|---|---|
| 8.12412798611906e-42 | 8.24910110868692 | 0.026 | 0 | 3.15760482436489e-37 | CBcDC2_C3 | RPL23AP31 |
| 8.12412798611906e-42 | 8.19260521497935 | 0.026 | 0 | 3.15760482436489e-37 | CBcDC2_C3 | PPIAP9 |
| 8.12412798611906e-42 | 8.95039600414974 | 0.026 | 0 | 3.15760482436489e-37 | CBcDC2_C3 | PAPOLB |
| 8.12412798611906e-42 | 7.95696601659719 | 0.026 | 0 | 3.15760482436489e-37 | CBcDC2_C3 | ENSG00000253389 |
| 8.12412798611906e-42 | 8.89954367988263 | 0.026 | 0 | 3.15760482436489e-37 | CBcDC2_C3 | ENSG00000230729 |
| 8.12412798611906e-42 | 8.1020793384517 | 0.026 | 0 | 3.15760482436489e-37 | CBcDC2_C3 | LINC02578 |
| 8.12412798611906e-42 | 8.16349636629498 | 0.026 | 0 | 3.15760482436489e-37 | CBcDC2_C3 | LINC01080 |
| 3.17957296461868e-273 | 8.69575341655974 | 0.522 | 0.003 | 1.23580462415834e-268 | CBcDC2_C4 | METTL7B |
| 1.31420814539478e-195 | 7.64861309326613 | 0.478 | 0.004 | 5.10793279870591e-191 | CBcDC2_C4 | MAFB |
| 1.75724212939486e-122 | 6.91216775851347 | 0.435 | 0.006 | 6.829872984319e-118 | CBcDC2_C4 | CD163 |
| 2.18512989095448e-90 | 7.36609287458061 | 0.304 | 0.004 | 8.49294434717279e-86 | CBcDC2_C4 | SCIN |
| 5.52226962872645e-86 | 6.53826984278626 | 0.609 | 0.02 | 2.14634053659711e-81 | CBcDC2_C4 | CD14 |
| 4.62666138911569e-68 | 9.14316743636872 | 0.043 | 0 | 1.7982444821076e-63 | CBcDC2_C4 | ENSG00000272510 |
| 4.62666138911569e-68 | 10.3074793162655 | 0.043 | 0 | 1.7982444821076e-63 | CBcDC2_C4 | GPR61 |
| 4.62666138911569e-68 | 9.0979298238908 | 0.043 | 0 | 1.7982444821076e-63 | CBcDC2_C4 | FABP5P6 |
| 4.62666138911569e-68 | 10.5726295781794 | 0.043 | 0 | 1.7982444821076e-63 | CBcDC2_C4 | ENSG00000255459 |
| 4.62666138911569e-68 | 10.6342776644941 | 0.043 | 0 | 1.7982444821076e-63 | CBcDC2_C4 | PCDH9-AS4 |
| 4.62666138911569e-68 | 9.87833142219433 | 0.043 | 0 | 1.7982444821076e-63 | CBcDC2_C4 | ENSG00000263017 |
| 3.27030306104538e-48 | 6.08989304516701 | 0.304 | 0.009 | 1.27106869073651e-43 | CBcDC2_C4 | JAKMIP2 |
| 9.88396962813725e-48 | 4.87147786428205 | 0.957 | 0.101 | 3.84160247536811e-43 | CBcDC2_C4 | CD9 |
| 3.08148356759737e-46 | 6.43365118659927 | 0.435 | 0.019 | 1.19768021821807e-41 | CBcDC2_C4 | PID1 |
| 8.29735316247311e-39 | 5.73032853029968 | 0.217 | 0.005 | 3.22493225365842e-34 | CBcDC2_C4 | BAALC |
| 7.02599626811702e-37 | 5.09464319599253 | 0.826 | 0.094 | 2.73079396952904e-32 | CBcDC2_C4 | GPNMB |
| 6.28714318636225e-36 | 6.87337962125199 | 0.13 | 0.002 | 2.44362394224342e-31 | CBcDC2_C4 | ACVRL1 |
| 1.23730843090059e-35 | 5.12712740222188 | 0.391 | 0.02 | 4.80904667838131e-31 | CBcDC2_C4 | DNASE2 |
| 1.03591301901451e-34 | 8.62612367054426 | 0.043 | 0 | 4.02628313100369e-30 | CBcDC2_C4 | PFN1P2 |
| 1.03591301901451e | 8.86317219906 | 0.04 | 0 | 4.02628313100369e | CBcDC2_ | TNNT2 |

| | | | | | | |
|---|---|---|---|---|---|---|
| -34 | 584 | 3 | | -30 | C4 | |
| 1.03591301901451e-34 | 8.59510094085664 | 0.043 | 0 | 4.02628313100369e-30 | CBcDC2_C4 | ENSG00000248802 |
| 1.03591301901451e-34 | 9.98788924123129 | 0.043 | 0 | 4.02628313100369e-30 | CBcDC2_C4 | ENSG00000258130 |
| 1.03591301901451e-34 | 8.26306385649602 | 0.043 | 0 | 4.02628313100369e-30 | CBcDC2_C4 | ENSG00000261541 |
| 1.03591301901451e-34 | 9.11064001139989 | 0.043 | 0 | 4.02628313100369e-30 | CBcDC2_C4 | LINC01549 |
| 1.08209797438011e-34 | 7.57512022429428 | 0.043 | 0 | 4.20579019702317e-30 | CBcDC2_C4 | ENSG00000285998 |
| 1.08209797438011e-34 | 8.23115417998844 | 0.043 | 0 | 4.20579019702317e-30 | CBcDC2_C4 | ENSG00000287550 |
| 5.17355515738521e-32 | 4.96265414484067 | 0.391 | 0.022 | 2.01080568302091e-27 | CBcDC2_C4 | LRP1 |
| 6.93144375227625e-29 | 4.70292351026212 | 0.565 | 0.053 | 2.69404424319721e-24 | CBcDC2_C4 | AOAH |
| 1.17243276790717e-26 | 4.39694524100202 | 0.435 | 0.033 | 4.55689443902481e-22 | CBcDC2_C4 | C3AR1 |
| 4.31541539268693e-26 | 5.14543839357117 | 0.217 | 0.008 | 1.67727250067563e-21 | CBcDC2_C4 | VCAN |
| 3.25302604944857e-25 | 6.94455910672907 | 0.087 | 0.001 | 1.26435363463918e-20 | CBcDC2_C4 | SLC1A7 |
| 3.51951992252736e-25 | 6.25360434962006 | 0.087 | 0.001 | 1.36793180828871e-20 | CBcDC2_C4 | ENSG00000287677 |
| 1.47721991410222e-23 | 8.35702214340318 | 0.043 | 0 | 5.7415106401411e-19 | CBcDC2_C4 | LINC02485 |
| 1.47721991410222e-23 | 9.42564657296788 | 0.043 | 0 | 5.7415106401411e-19 | CBcDC2_C4 | RNU1-73P |
| 1.47721991410222e-23 | 8.56480442475556 | 0.043 | 0 | 5.7415106401411e-19 | CBcDC2_C4 | ENSG00000254065 |
| 1.47721991410222e-23 | 8.11379621691406 | 0.043 | 0 | 5.7415106401411e-19 | CBcDC2_C4 | ENSG00000275488 |
| 1.47721991410222e-23 | 7.93113583283565 | 0.043 | 0 | 5.7415106401411e-19 | CBcDC2_C4 | LINC02821 |
| 1.47721991410222e-23 | 8.28645983165507 | 0.043 | 0 | 5.7415106401411e-19 | CBcDC2_C4 | IGBP1P1 |
| 1.52081649199749e-23 | 7.96642545722806 | 0.043 | 0 | 5.91095745944664e-19 | CBcDC2_C4 | CFL1P2 |
| 1.5656868552936e-23 | 7.26387284160066 | 0.043 | 0 | 6.08535510046964e-19 | CBcDC2_C4 | OR10J5 |
| 1.5656868552936e-23 | 6.71863090008487 | 0.043 | 0 | 6.08535510046964e-19 | CBcDC2_C4 | ENSG00000287391 |
| 1.5656868552936e-23 | 7.43003978018873 | 0.043 | 0 | 6.08535510046964e-19 | CBcDC2_C4 | LINC01597 |
| 1.71341351107717e-23 | 5.45338559515423 | 0.261 | 0.014 | 6.65952429350364e-19 | CBcDC2_C4 | LNCAROD |
| 1.01173252034386e-22 | 4.18939017422224 | 0.609 | 0.078 | 3.93230078682048e-18 | CBcDC2_C4 | GAPT |
| 5.85729262139528e-21 | 4.32062822083513 | 0.391 | 0.034 | 2.2765539231577e-16 | CBcDC2_C4 | CCL3 |
| 5.61037274579432e-20 | 6.66044008258643 | 0.087 | 0.002 | 2.18058357510788e-15 | CBcDC2_C4 | VCAN-AS1 |

| | | | | | | |
|---|---|---|---|---|---|---|
| 5.89972284289669e-19 | 5.38884118084015 | 0.174 | 0.008 | 2.29304527734866e-14 | CBcDC2_C4 | CTSV |
| 9.95498639627264e-19 | 5.544747898 | 0.174 | 0.008 | 3.86920456263929e-14 | CBcDC2_C4 | GIMAP4 |
| 5.81608891964189e-18 | 7.89116943950063 | 0.043 | 0 | 2.26053928039721e-13 | CBcDC2_C4 | ENSG00000227815 |
| 5.81608891964189e-18 | 7.87047681406017 | 0.043 | 0 | 2.26053928039721e-13 | CBcDC2_C4 | ENSG00000228251 |

**Table S2.**

**The Number of Cells in Each Single-cell Subpopulations.**

| | Before quality control | Post quality control |
|---|---|---|
| total | 54306 | 48729 |

| Total_Cell Clusters | | | | | | | | |
|---|---|---|---|---|---|---|---|---|
| CBcDC1 | CBcDC2 | MEP_like | Neutrophil_like | GP_like | HSPC | CD1C+ cell | DC_progenitors | LAMP3+ DC |
| 10397 | 7015 | 10678 | 6394 | 7438 | 3318 | 2444 | 857 | 188 |

| CBcDC1 Subclusters | | | | |
|---|---|---|---|---|
| CBcDC1_C0 | CBcDC1_C1 | CBcDC1_C2 | CBcDC1_C3 | CBcDC1_C4 |
| 3098 | 2553 | 2290 | 1372 | 1084 |

| CBcDC2 Subclusters | | | | |
|---|---|---|---|---|
| CBcDC2_C0 | CBcDC2_C1 | CBcDC2_C2 | CBcDC2_C3 | CBcDC2_C4 |
| 3347 | 2927 | 680 | 38 | 23 |

**Table S3. Cell Counts in Tumor scRNA-seq Analysis.**

| | $CD8^+$ T | $CD4^+$ T | B | Monomac | cDC1 | $LAMP3^+$ DC | pDC |
|---|---|---|---|---|---|---|---|
| Ctrl | 3615 | 309 | 1510 | 1817 | 426 | 61 | 1177 |
| Treat | 7636 | 1185 | 653 | 262 | 29 | 93 | 233 |

| | CD8_Tn | CD8_Trm | CD8_ISG | CD8_innate_like | CD8_early_activated | CD8_cycling | CD8_Teff | CD8_Texh |
|---|---|---|---|---|---|---|---|---|
| Ctrl | 89 | 500 | 114 | 975 | 108 | 634 | 439 | 756 |
| Treat | 633 | 1214 | 398 | 697 | 215 | 2253 | 1100 | 1126 |

| | CD4_Tfh | CD4_Tem | CD4_Teff | CD4_Trm | CD4_Treg | CD4_cycling |
|---|---|---|---|---|---|---|
| Ctrl | 12 | 35 | 72 | 135 | 49 | 6 |
| Treat | 165 | 213 | 195 | 249 | 329 | 34 |

**Table S4. Materials and Reagents.**

| Reagent | Source | Cat# |
|---|---|---|
| CliniMACS® CD34 Reagent CR/GMP | Miltenyi Biotech | 220-001-831 |
| CryoStor CS10 | Biolife Solutions | 210102 |
| human naive CD8+ T cell isolation kit | Miltenyi Biotech | 130-093-244 |
| StemSpan™-AOF medium | STEMCELL Technologies | #100-0130 |
| Recombinant Human SCF GMP Protein | R&D systems | BT-SCF-GMP |
| Recombinant Human Thrombopoietin/TPO GMP Protein | R&D systems | BT-TPO-GMP |
| Recombinant Human Flt-3 Ligand/FLT3L GMP Protein | R&D systems | BT-FT3L-GMP |
| SR-1 | STEMCELL Technologies | #72342 |
| Recombinant Human IFN-gamma GMP Protein | R&D systems | 285-GMP |
| Recombinant Human GM-CSF GMP Protein | R&D systems | 215-GMP |
| Recombinant Human IL-4 GMP Protein | R&D systems | BT-004-GMP |
| Lonza X-VIVO 15 medium | Lonza Bioscience | BEBP02-061Q |
| CliniMACS® CD141 (BDCA-3) Reagent CR/GMP kit | Miltenyi Biotech | 220-001-832 |
| Poly I:C | Invivogen | tlrl-pic |
| Pam3CSK4 | Invivogen | tlrl-pms |
| R848 | Invivogen | tlrl-r848-1 |
| ODN2216 | Invivogen | tlrl-2216 |
| human CD3/CD28 T cell activation beads | BioLegend | 422604 |
| FcR Blocking Reagent | Miltenyi Biotec | 130-059-901 |

| | | |
|---|---|---|
| Cell Activation Cocktail containing Brefeldin A | BioLegend | 423304 |
| BD Cytofix/Cytoperm™ Fixation/Permeabilization Kit | BD bioscience | 554714 |
| BD Perm/Wash buffer | BD bioscience | 554723 |
| InVivoMAb anti-human CD8α | Bio X Cell | #BE0004-2 |
| InVivoMAb anti-human CD4 | Bio X Cell | #BE0003-2 |
| InVivoMAb mouse IgG2 isotype control | Bio X Cell | #BE0085 |
| GenCRISPR™ Ultra eSpCas9-2NLS-GMP | GenScript | Z03624-GMP |
| Neon™ Transfection System 10 μL Kit | ThermoFisher Scientific | MPK1096 |
| NY-ESO-1 peptide, human | Miltenyi Biotech | 130-095-381 |
| CellTrace™ Violet | ThermoFisher Scientific | C34571 |
| BD™ Cytometric Bead Array (CBA) Human IL-12p70 Flex Set | BD bioscience | 558283 |
| BD™ Cytometric Bead Array (CBA) Human IL-6 Flex Set | BD bioscience | 558276 |
| BD™ Cytometric Bead Array (CBA) Human IFN-γ Flex Set | BD bioscience | 558269 |
| BD™ Cytometric Bead Array (CBA) Human IP-10 Flex Set | BD bioscience | 558280 |
| Human IL-29 ELISA Kit | absin | abs551325 |
| APC anti-human CD3 Antibody | BioLegend | 300412 |
| PE/Cyanine7 anti-human CD3 Antibody | BioLegend | 300420 |
| FITC anti-human CD4 Antibody | BioLegend | 317408 |
| Brilliant Violet 421™ anti-human CD4 Antibody | BioLegend | 317434 |

| | | |
|---|---|---|
| PerCP/Cyanine5.5 anti-human CD8 Antibody | BioLegend | 344710 |
| APC/Cyanine7 anti-human CD45 Antibody | BioLegend | 368516 |
| FITC anti-mouse CD45 Antibody | BioLegend | 103108 |
| PE/Cyanine7 anti-human CD141 (Thrombomodulin) Antibody | BioLegend | 344110 |
| APC anti-human CD141 (Thrombomodulin) Antibody | BioLegend | 344106 |
| PE anti-human CD1c Antibody | BioLegend | 331506 |
| Brilliant Violet 650™ anti-human CD1c Antibody | BioLegend | 331542 |
| Brilliant Violet 510™ anti-human HLA-DR Antibody | BioLegend | 307646 |
| FITC anti-human HLA-DR Antibody | BioLegend | 307604 |
| Pacific Blue™ anti-human Lineage Cocktail (CD3, CD14, CD16, CD19, CD20, CD56) | BioLegend | 348805 |
| PE anti-human CD370 (CLEC9A/DNGR1) Antibody | BioLegend | 353804 |
| Pacific Blue™ anti-human CD123 Antibody | BioLegend | 306043 |
| Brilliant Violet 421™ anti-human AXL Antibody | BioLegend | 386510 |
| APC anti-human XCR1 Antibody | BioLegend | 372606 |
| PerCP/Cyanine5.5 anti-human CD11c Antibody | BioLegend | 337210 |
| Brilliant Violet 421™ anti-human CD80 Antibody | BioLegend | 305222 |
| FITC anti-human CD40 Antibody | BioLegend | 334306 |
| PE anti-human CD86 Antibody | BioLegend | 305406 |
| Brilliant Violet 510™ anti-human CD14 Antibody | BioLegend | 301842 |
| APC anti-human CD172a (SIRPα) Antibody | BioLegend | 372106 |
| PE anti-human CD103 (Integrin αE) Antibody | BioLegend | 350206 |
| CADM1 Antibody (41N3D3) [Biotin] | Novus Biologicals | NBP2-14841B |
| Brilliant Violet 510™ Streptavidin | BioLegend | 405234 |

| | | |
|---|---|---|
| PE anti-human/mouse Granzyme B Recombinant Antibody | BioLegend | 396406 |
| APC anti-human IFN-γ Antibody | BioLegend | 502512 |
| PE anti-human TNF-α Antibody | BioLegend | 502909 |
| APC anti-human HLA-A2 Antibody | BioLegend | 343308 |
| FITC anti-mouse CD326 (Ep-CAM) Antibody | BioLegend | 118208 |
| PE anti-human CD326 (EpCAM) Antibody | BioLegend | 324206 |
| PerCP/Cyanine5.5 anti-human TCR γ/δ Antibody | BioLegend | 331224 |
| Cell Staining Buffer | BioLegend | 420210 |
| HLA-A*02:01 NY-ESO-1 Tetramer-SLLMWITQC-PE | MBL Life Science | TS-M011-1 |
| Ficoll-Paque™ PREMIUM density gradient media | Cytiva | 17544203 |
| Red blood cell lysis buffer | ThermoFisher Scientific | 00-4333-57 |
| DPBS | BasalMedia | B210KJ |
| DMEM (Dulbecco's Modified Eagle Medium) | ThermoFisher Scientific | 11965092 |
| RPMI 1640 Medium | ThermoFisher Scientific | 61870010 |
| Penicillin-Streptomycin | ThermoFisher Scientific | 15070063 |
| Fetal Bovine Serum (FBS) | ThermoFisher Scientific | A5670801 |
| BD Horizon™ Fixable Viability Stain 575V | BD Bioscience | 565694 |
| McCoy's 5A medium | ThermoFisher Scientific | 36600021 |

| | | |
|---|---|---|
| Leibovitz L-15 medium | ThermoFisher Scientific | 11415064 |
| Trypsin-EDTA | ThermoFisher Scientific | 25200072 |
| Collagenase IV | Sigma-Aldrich | C5138 |
| DNase I | Roche | 10104159001 |
| A375 cell line | National Collection of Authenticated Cell Cultures | SCSP-533 |
| SKOV3 cell line | National Collection of Authenticated Cell Cultures | SCSP-5214 |
| MDA-MB-231 cell line | National Collection of Authenticated Cell Cultures | SCSP-5043 |

**Table S5. SgRNA Sequences for Gene Editing.**

| | Sequence(5’-3’) |
|---|---|
| sgRNA_NC | CTGAAAAAGGAAGGAGTTGA |
| sgRNA_TCF4 | TGGCGAGTCCCTATTGTAGT |
| sgRNA_IRF8 | AGGCCTGGGCAGTTTTTAAA |
| sgRNA_IFNGR1 | GGTCCCTGTTTTTACCGTAG |
| sgRNA_BATF3 | GAAGGCTGACAAGCTCCATG |
| sgRNA_IRF4 | TCTTGTTCAAAGCGCACCGC |
| sgRNA_ZEB2 | TGTTTGCGCCTCTTGCACCG |
